# Electrical Unfolding of Cytochrome c During Translocation Through a Nanopore Constriction

**DOI:** 10.1101/2021.02.10.430607

**Authors:** Prabhat Tripathi, Abdelkrim Benabbas, Behzad Mehrafrooz, Hirohito Yamazaki, Aleksei Aksimentiev, Paul M. Champion, Meni Wanunu

## Abstract

Many small proteins move across cellular compartments through narrow pores. In order to thread a protein through a constriction, free energy must be overcome to either deform or completely unfold the protein. In principle, the diameter of the pore, along with the effective driving force for unfolding the protein, as well as its barrier to translocation, should be critical factors that govern whether the process proceeds via squeezing, unfolding/threading, or both. To probe this for a well-established protein system, we studied the electric-field-driven translocation behavior of cytochrome c (cyt c) through ultrathin silicon nitride (SiN_x_) solid-state nanopores of diameters ranging from 1.5 to 5.5 nm. For a 2.5 nm diameter pore we find that, in a threshold electric field regime of ∼30-100 MV/m, cyt c is able to squeeze through the pore. As electric fields inside the pore are increased, the unfolded state of cyt c is thermodynamically stabilized, facilitating its translocation. In contrast, for 1.5 nm and 2.0 nm diameter pores, translocation occurs only by threading of the fully unfolded protein after it transitions through a higher energy unfolding intermediate state at the mouth of the pore. The relative energies between the metastable, intermediate, and unfolded protein states are extracted using a simple thermodynamic model that is dictated by the relatively slow (∼ms) protein translocation times for passing through the nanopore. These experiments map the various modes of protein translocation through a constriction, which opens new avenues for exploring protein folding structures, internal contacts, and electric field-induced deformability.

**Significance Statement:** Can localized electric fields drive the complete unfolding of a protein molecule? Protein unfolding prior to its translocation through a nanopore constriction is an important step in protein transport across biological membranes and also an important step in nanopore-based protein sequencing. We studied here the electric-field-driven translocation behavior of a model protein (cyt c) through nanopores of diameters ranging from 1.5 to 5.5 nm. These single molecule measurements show that electric fields at the nanopore constriction can select both partially and fully unfolded protein conformations. Zero-field free energy gaps between these conformations, found using a simple thermodynamic model, are in remarkable agreement with previously reported studies of cyt c unfolding energetics.

## Main Text

### Introduction

Protein unfolding during its translocation through a nano-constriction, and its subsequent refolding after translocation, are two ubiquitous processes in biology (1-6). In order to fully understand the two processes, a plethora of experiments that use nanopores as mimics of a real biological constriction have been performed (7-24). In these studies, voltage applied across a pore electrokinetically pulls a protein into and subsequentially through it. These studies are also critical to overcoming technical challenges associated with protein sequencing using nanopores, where tertiary and secondary structures must be unfolded to allow single-file threading of a protein through the nanopore sensor. Interactions of partially and fully unfolded proteins with biological and solid-state pores in the presence of chemical denaturants have been studied extensively (7-10). The use of an enzymatic motor to achieve processive threading of unfolded proteins was demonstrated recently (11). In other pioneering experiments, electric field-driven unidirectional threading was demonstrated by tagging the end terminal of the model protein thioredoxin with an oligonucleotide (12, 13). In these studies, the size of the pore was smaller than that of a fully folded protein (d_pore_ < d_protein_) and translocation of protein necessarily required denaturing agents, an enzyme, or an oligo tag. Nanopores with larger diameters than the folded protein (d_pore_ > d_protein_) were also employed (8, 14-22), and due to the extremely fast translocation times of folded protein transport, only a tiny fraction of the translocated population was detected(14, 15, 17, 19), predominantly the longer events associated with protein sticking to the pore walls (15, 19). Slowing protein translocation by tethering to a lipid coating on the pore walls achieved orders of magnitude reduction in the translocation times and allowed efficient protein detection (18). High-bandwidth measurements combined with the use of “tighter” pore diameters have been used to capture the fast translocation events of folded proteins and further utilized to estimate size (16, 20, 25), conformation (21), and conformational flexibility (22).

Here we report that a fully folded heme protein cytochrome c (cyt c, Fig. 1A) can pass through an ultra-thin solid-state nanopore (d_pore_ < d_protein_) and translocate without requiring chemical denaturants, an unfolding enzyme, or an oligonucleotide tag. Instead, protein unfolding during translocation can be achieved by controlling the electric field across the pore. To the best of our knowledge, this is the first report on how solid-state nanopores can unfold a protein during its translocation through a nano-constriction that is smaller than the protein diameter. The nano-constriction allows studying the protein in a “trapped” mode where the native state of the protein is not allowed to pass through, but partial unfolding and re-equilibration of the protein can take place at the pore. This leads to an outcome where unfolding of the remaining secondary *α*-helical structure can be induced by an electric field during translocation. Although this experiment is fundamentally different from DNA unzipping (26-28), stretching (29), and translocation (30), there are certain similar outcomes in the analysis. For example, we observed that there is a voltage or electric field threshold to observe transitions in the protein conformation, and we find that the translocation rate is conformation dependent.

**Figure 1.**
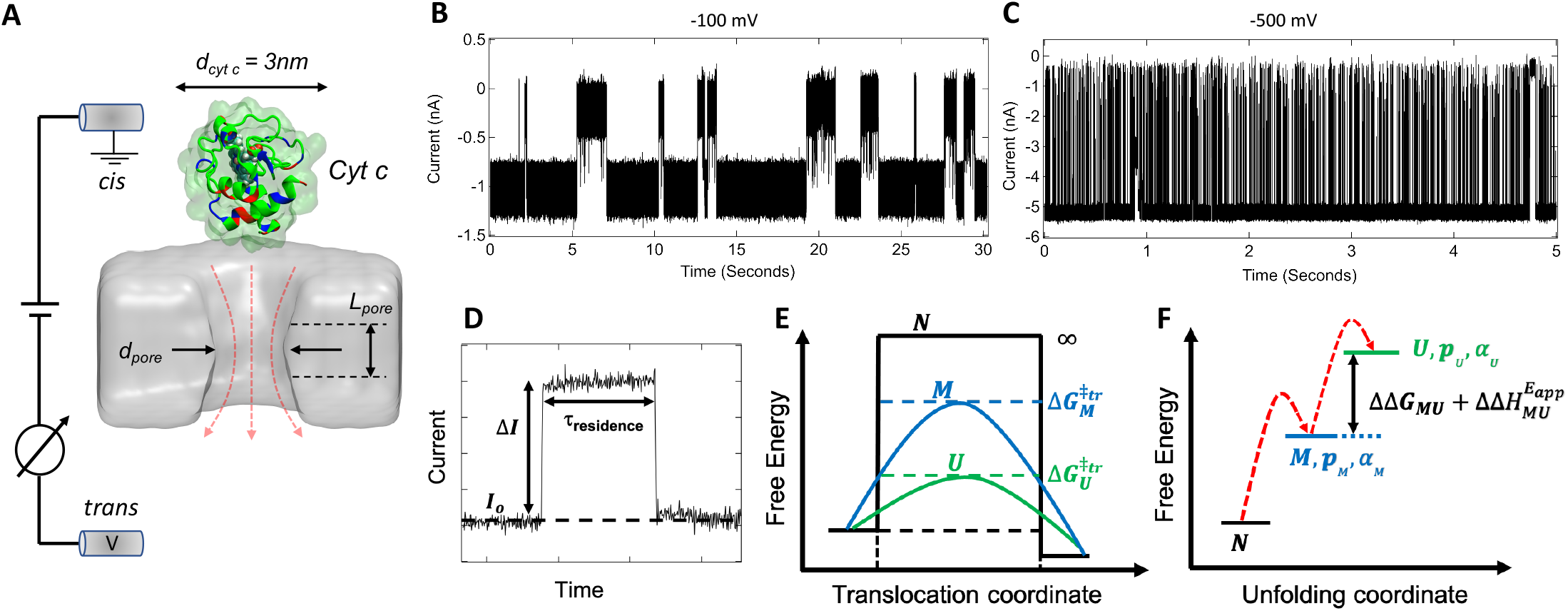
Cytochrome c squeezing/unfolding through a solid-state silicon nitride nanopore. **(A)** Schematic of our experimental setup: Application of a negative voltage to the *trans* chamber electrokinetically captures cyt c molecules at the pore vicinity. **(B, C)** Representative ionic current recordings for a d_pore_ = 2.5 nm (L_pore_ = 3.4 nm), in 1 M KCl, 10 mM HEPES, pH 7.5 buffer at −100 mV and (C) at −500 mV, respectively. **(D)** Description of the parameters in a typical current blockade event. **(E)** Schematic energy diagram for translocation of cyt c. In the native state (*N*), the energy barrier for protein translocation through the pore is assumed to be pseudo-infinity due to steric constraints, whereas the energy barrier of a partially unfolded metastable squeezed state (M), which still has intact a-helices, remains relatively large compared to the completely unfolded state, *U*. Due to interaction with an external electric field and fast *α*-helical conformational transitions, the low-barrier *U*-state population increases leading to larger average translocation rates. **(F)** The conformational populations of the protein are governed by thermal excitations, the zero-field free energy gap between conformations (ΔΔ*G*_*MU*_), and the effect of the external electric field 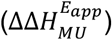.

#### Cytochrome c Interaction with a Nanopore

We chose cyt c as our model protein (SI: Sec.2, Fig. S1 (i) and (ii)) because of ample investigations concerning the energetics of its conformationally-excited metastable and unfolded states (31-33). In its native state, cyt c is relatively small (MW = 12 kDa, d_protein_ ∼ 3 nm), positively charged (+8 at physiological pH), its structure can be modulated under the action of electric field (34), and it can translocate through mitochondrial pores (6, 35, 36) and other small constrictions (37) (d_pore_ < 3 nm). Moreover, some conformational changes of cyt c are known to involve the removal of the Met 80 heme ligand (38-42). These conformational changes are known to be functionally relevant: for example, upon interaction with the mitochondrial membrane, cyt c loses the Met 80 heme ligand and transitions from an electron transport protein to a peroxidase, initiating the process of apoptosis (43-46).

Earlier studies on cyt c reported that the energy required to unfold cyt c is > 21.6 *k*_*B*_*T*_*0*_ (12.8 kcal/mol with *T*_*0*_ =298 K) (32), a significantly high energy (> 500 meV) to achieve on a lipid-supported biological nanopore platform, even at maximally-supported voltages of up to 350 mV (47). In contrast, solid-state nanopores allow us to perform experiments at high voltage (∼1 V) and also robustly analyze the same molecule with different pore diameters while maintaining the electrostatic and hydrophobic environment of the inner pore lumen (48-50). The arrangement of a solid-state nanopore within the flow cell is shown in Fig. 1A where a negative potential is applied to the *trans* compartment. The resulting steady-state ion current is transiently interrupted only by cyt c molecules interacting with the pore. The electric field intensity is maximum inside the pore, and decays rapidly outside the pore entrance (29, 50). The positively-charged cyt c is captured at the pore entrance *via* a drift-diffusion mechanism (24, 50), and following its trapping at the pore mouth, depending upon the applied potential, it can either undergo conformational transitions and translocate through the pore or escape back to the *cis* compartment. Given the dipolar nature of cyt c, we expect that in the applied voltage regime used here, the resulting electric field in the pore vicinity can preferentially orient and separate the differentially-charged segments of cyt c prior to its translocation (SI: Sec. 5). The orientation-dependent energetics of the protein, due to permanent and induced dipole interactions with the electric field (and its gradient) are, for simplicity, treated here using scalar fitting parameters. We also emphasize that rapid timescale (0.1-10 μs) *α*-helical unfolding/folding processes (51-56) are key aspects of the secondary structure of cyt c, and that these relatively fast conformational transitions appear to govern the much slower (∼ms) translocation of the protein through pores with diameters of 2.0-2.5 nm.

Example traces at −100 mV and −500 mV are shown in Fig. 1B and Fig. 1C, respectively (for other voltages see SI: Sec. 6 Fig. S3(i-vii)). A single interaction of a protein molecule with the pore, shown in Fig. 1 D, can be characterized by a reduction in the open pore current (I_0_) by some amplitude (ΔI), as well as by the duration of this interaction time (*τ*_*residence*_). As can be discerned from previous work (19, 57), as well as from the molecular dynamics (MD) simulations presented below, the scaleless fractional blockade ΔI/I_*0*_ provides information about the nature of the protein occupancy in the pore (e.g., squeezed, stretched, or unfolded), while the value of *τ*_*residence*_ provides information about the rate of protein translocation and its associated free energy barrier. We reason that translocation for d_pore_ < d_protein_ requires excitation of the native protein, *N*), to higher energy, partially unfolded, conformations. Thus, in addition to *N*), we consider a metastable squeezed state, *M*, a fully unfolded state, *U*, and an intermediate state,*U*, on the pathway between *M* and *U*. The residual *α*-helical secondary structures associated with the *M*- and *I*-states are described in more detail below. As the electric field increases, these states can interconvert, either in the nanopore or at the mouth of the pore, making the protein more flexible and reducing both the fractional blockade (ΔI/I_*0*_) and the energetic barrier to translocation, 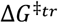.

#### Thermodynamic model

In the studies reported here, translocation of cyt c through a 2.5 nm pore takes place on ∼ms timescales. This means that protein secondary structures that re-equilibrate on much faster (≲ 10 μs) timescales can be considered within a thermodynamic quasi-equilibrium framework (i.e., using a pre-equilibrium kinetic assumption). The thermodynamic analysis describes the translocation process under near-equilibrium conditions for the *α*-helix state transitions, while the MD simulations described below probe highly non-equilibrium transport on ∼ns timescales. This two-pronged approach allows us to consider the non-equilibrium processes that are likely to occur, even on the experimentally observed (∼ms) translocation timescale, as well as to analyze the thermally driven conformational re-equilibration processes that are constantly taking place on much faster timescales.

As an example, in Fig. 1E we illustrate the energy diagram for translocation within a pore that denies access to the)-state but allows entrance and “squeezing” of a partially unfolded *M*-state, which has the three residual *α*-helices of cyt c intact (32, 58, 59). For smaller pores, which do not allow access to *M*, conformational excitations due to the electric field at the mouth of the pore can still take place and affect the ion current. Thus, for the experiments using a 2.0 nm diameter pore described below, a two-state analysis was used to describe interconversions between *M* and an intermediate *I*-state that lies on the pathway to U where one of the α-helices has unfolded (32, 58, 59). In a larger 2.5 nm pore, where the *M*-state of cyt c can enter by squeezing, both two-state (*M* ↔ *U*) and three-state (*M* ↔ *I* ↔ *U*) models were used for analysis. These models are discussed in more detail in sections 7 and 15 of the SI. In both models, a flexible dynamically interconverting thermodynamic state mixture, associated with rapid (51-56) *α*-helix folding/unfolding transitions, facilitates translocation through the pore and affects the observed fractional blockades.

The conformational excitation of an electrically polarized (or polarizable) protein to a state of higher electric dipole moment can be induced by the action of an applied electric field. This occurs because the applied field reduces the energy gap between conformational states with differing net dipole moments. As an example, using just the two states *M* and *U* (in addition to)), we depict in Fig. 1F an inherent zero-field free energy gap, ΔΔ*G*_*MU*_ that is reduced by a field-dependent interaction energy. The field-dependence of the energy gap is defined as: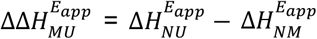. More specifically, the electric field-dependent energy gap between the states *M* and *U* can be written as:

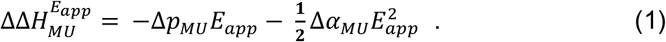

Here, Δ*p*_*MU*_ = *p*_*U*_ *- p*_*M*_ and Δ*α*_*MU*_ = *α*_*U*_ − *α*_*M*,_ are taken to be positive fitting parameters that include dipole-field orientation as well as any associated dielectrophoretic effects (29) where the gradient is expanded in powers of applied field. The field primarily interacts with either the permanent dipole moment (*p*_*M*_ and *p*_*U*_) of each state or with the conformation-dependent induced dipoles (34) that depend on both the polarizability and the applied field (i.e., *α*_*M*_ *E*_*app*_ and *α*_*U*_ *E*_*app*_). As the electric field is increased, we expect the dipoles to orient along the field so that *p*_*U*_ >*p*_*M*_ and *α*_*U*_ E_*app*_ > *α*_*M*_ E_*app*_, due to the protein elongation (and the charges within it) in the *U*-state conformation. Under this condition, the potential energy level associated with *U* will decrease relative to *M* as the magnitude of the field increases.

Thus, as the applied field increases, an energy level degeneracy occurs, where the thermodynamic population probabilities of *M* and *U* become equal, and *M* is no longer a thermodynamically-stabilized state. The resulting rapid (0.1-1 μs) timescale (51-56) thermodynamic *α*-helix unfolding/folding leads to a population mixture between *M* and *U* that affects the fractional blockade. It also facilitates translocation through the pore, but on much slower (ms) timescales than the underlying *α*-helix interconversion times. At still higher fields, level inversion takes place and the unfolded conformation (*U*) becomes more thermodynamically stable than *M*. If only one of the three *α*-helicies unfolds (32, 58, 59), a sequential intermediate state, *I*, should be included in the thermodynamic analysis. Finally, we note that, so long as the timescale separation that allows thermodynamic averaging is maintained, we do not need to explicitly consider the frictional forces that might affect the transition rates between the folded and unfolded *α*-helices linking the *M, I*, and *U* states.

### Results and Analysis

We present here extensive measurements of the fractional change in current amplitudes and residence times for many *single-molecule passage events of* cyt c through various solid-state pores under native and denaturing conditions. These experiments shed light on the kinetics and energetics of cyt c translocation by using a statistical analysis of the molecular ensemble. We use the thermodynamic model to analyze the electric field-dependent fractional blockades and, for the 2.5 nm pore, we extract the zero-field free energy gap, ΔΔ*G*_*MU*_ = Δ*G*_*NU*_ − Δ*G*_*NM*_. The results from the 2.0 nm pore also allow ΔΔ*G*_*M*I_ to be found. Thus, by use of both the pore size and the external electric field, the relative energies of different partially folded conformations of cyt c can be probed. As described below, the zero-field free energy gaps obtained with this single molecule nanopore technique are in remarkable agreement with prior studies of cyt c unfolding energetics (32, 58). Finally, the zero-field translocation energy barriers for the *M*-state 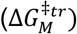, *U*-state, 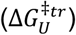, and the *I*-state 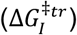 can also be estimated from the residence time (kinetic) measurements.

#### Smaller pore imposes a free energy barrier during cyt c translocation

Scatter plots of fractional blockades vs. residence times for different pore diameters are shown in Fig. 2A. Example traces for different pore diameters are shown in SI: Sec. 8, Fig. S4. For the 5.5 nm pore (V = 100 mV), we observe short-lived events (∼200 µs) with relatively shallow fractional blockades (∼0.36). Decreasing the pore size to 3.5 nm resulted in similar event durations (∼120 µs) yet deeper fractional blockades (∼0.8), as expected. Further decreasing the pore diameters to 3.0 and 2.5 nm increased the fractional blockades to near unity, and greatly increased event durations to 10-100 ms, orders of magnitude greater than for the larger pore sizes. Moreover, in the experiments with 3.0 nm and 2.5 nm pores, higher voltages were required to observe translocation events than for the larger pores. Given the molecular dimensions of cyt c, these results suggest that decreasing the pore diameter to below 3 nm imposes a significant free energy barrier for cyt c translocation in its native state, i.e., conformational excitation is required for the protein to move through the pore.

**Figure 2.**
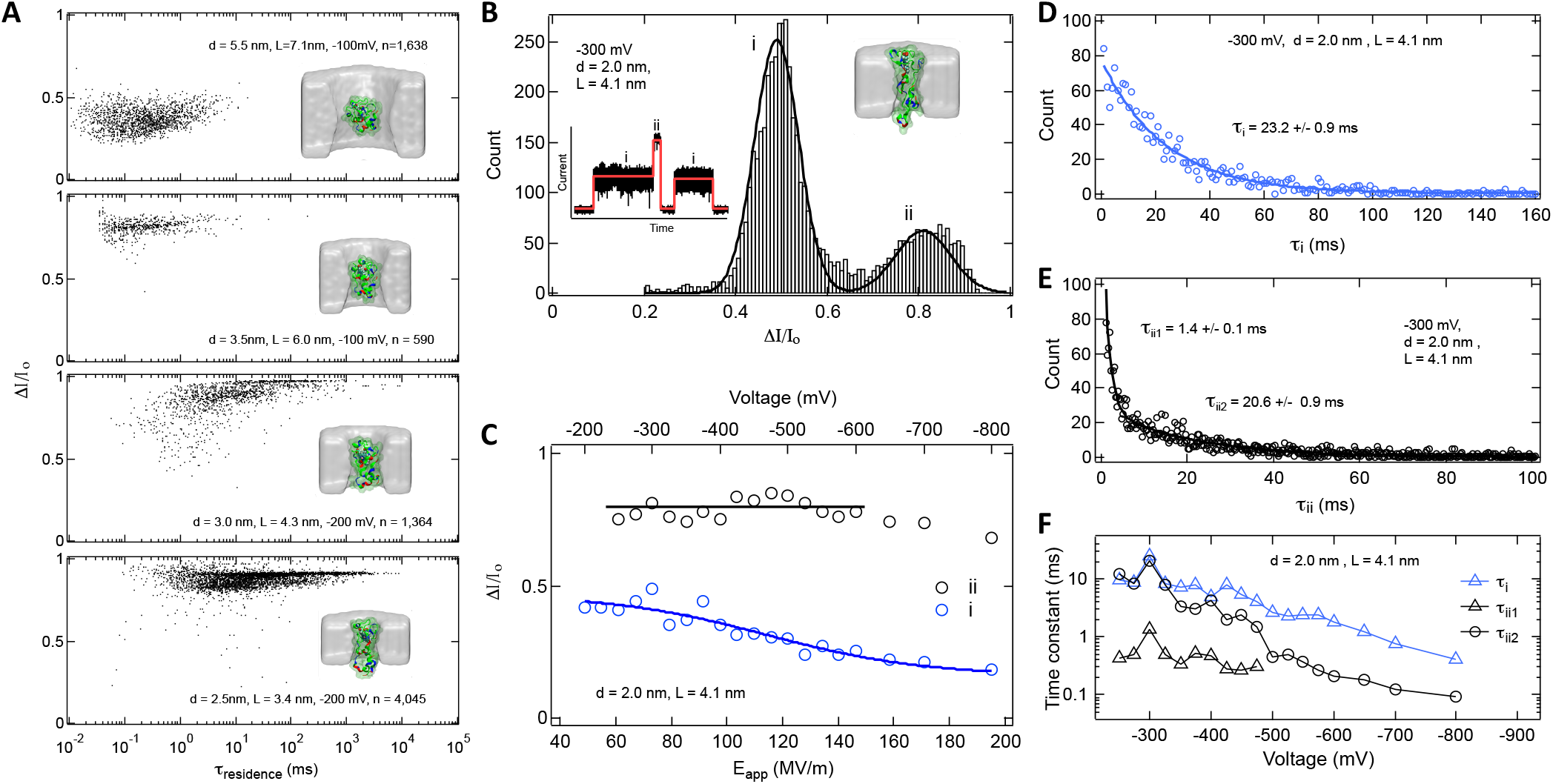
Conformationally excited states of cyt c during translocation. **(A)** Scatter plots of fractional current blockades and residence times for different pore dimensions. Insets: Snapshots from all-atom MD simulations. **(B)** Current blockade distribution for cyt c in a 2.0 nm pore shows two distinct populations of blockade events. Black curve represents a double-Gaussian fit. Inset: Example multi-level events and two-level fits (red curve) reveal levels *i* and *ii* **(C)** Voltage dependence of mean fractional blockades for levels *i* and *ii*. Level *ii* is nearly independent of voltage, whereas level *i* decreases with voltage. The blue curve is a fit using the two-state model described by Eq. 2, where the *M-* and *I*-state are interconverting at the mouth of the pore, prior to a transition to the *U*-state and threading into the pore. Fit parameters are given in Table 1. **(D-F)** Distribution of the level *i* lifetime with a single-exponential fit for *τ*_*i*_ (blue curve), distribution of a the level *ii* lifetime with a double-exponential fit for *τ*_*ii*1_ and *τ*_*ii*2_ (black curve), and voltage dependence of time constants *τ*_*i*_, *τ*_*ii1*_ and *τ*_*ii*2_, respectively.

#### Threading of the unfolded protein after a transition at the mouth of the 2.0 nm diameter pore

As the pore size is further reduced, we observe increases in the magnitude of the translocation barrier. In Figs. 2B - 2F we present results for a 2.0 nm diameter pore. A representative distribution of fractional blockades obtained for the 2.0 nm pore is shown in Fig. 2B. Since we observed two characteristic levels (*i* and *ii*) produced by the protein, which often occur in succession (*i → ii* for the same molecule, see inset trace), we decoupled the two states by using a custom script to independently quantify the durations and blockade levels for each of these states (see SI Sec. 1). Based on these observations we assigned level *i* to protein docking at the pore mouth where it attempts (but fails) to squeeze through the pore via the partially unfolded *M*-state, which retains its full *α*-helical content. Instead of squeezing, the *M*-state begins to undergo transitions with another unfolding intermediate (*I*-state, where the shortest of the three cyt c *α*- helices is unfolded) at the mouth of the pore. Further *α*-helical unfolding results in the *U*-state, which either escapes back to the *cis* chamber or threads deeper into the pore leading to blockage level *ii*. Similar event shapes (*i → ii*) were observed for a 1.5 nm diameter pore at −500 mV (see SI: Sec. *i*, Figs. S5A, S5B).

**Table 1:**
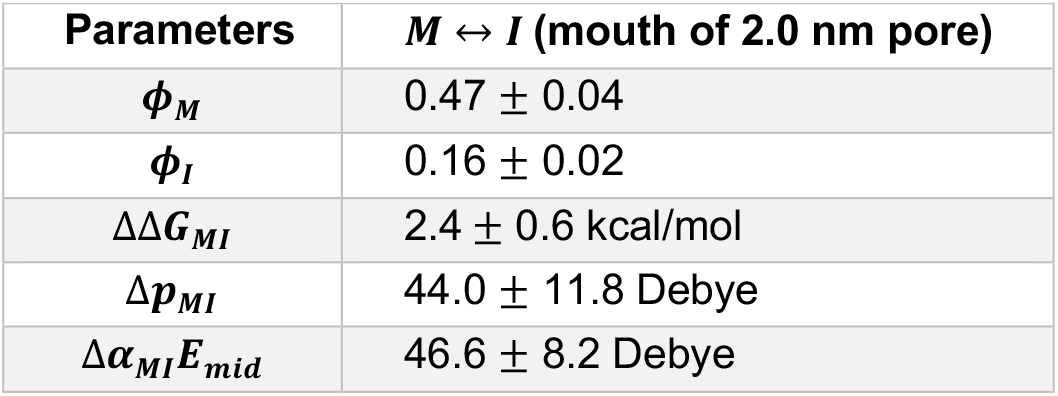
Fit parameters from the experimental data (d_pore_ = 2.0 nm, L = 4.1 nm) in Fig. 2C using Eq. 2 with either Δ*p*_*MI*_ or Δ*α*_*MI*_ as the dipole interaction term.

In additional studies, we did not observe a substantial number of events with level *ii* at voltages weaker than −250 mV, indicating that there is a significant voltage threshold necessary to fully unfold cyt c so that it is able to enter the 2.0 nm pore constriction. As higher voltages were applied, the number of level *ii* events increased up to about −400 mV and then leveled off (see SI: Sec. 10, Fig. S6). Further, whereas the ΔI/I_*0*_ peak position of level *ii* (Fig. 2C) remains constant as a function of voltage in the range −250 mV < V < −650 mV, we find that the blockade level systematically decreases with stronger applied voltage. This behavior stands in contrast to DNA unzipping (26-28) and translocation experiments (30), where fractional blockades do not change with the voltage. We attribute this behavior (Fig. 2C blue curve) to electric field-induced transitions of cyt c between the partially unfolded metastable *M*-state and the more unfolded state, *I*, which leads to more ion permeation depending on the field dependent relative population of the *M*- and *I*-states. When the protein fully unfolds to *U*, the level *ii* blockade ratio is established because the protein is now finally able to enter into the 2.0 nm pore where it blocks the ion current.

We assign the formation of the cyt c *M*-state to electric field-induced breakage of the salt bridge involving E62, which subsequently releases residues 40-57 within an *Ω*-loop region (59). Because the applied field should also lead to dissociation of Met80 (34), the unfolding of its associated *Ω*-loop residues 70-87 is also assumed to take place. Thus, we suggest that the action of the electric field at the entrance to the nanopore can induce the lowest energy unfolding transitions of cyt c (32, 58). This produces the smaller and more compressible *M*-state, which is chiefly composed of the three primary *α*-helices of cyt c. Because these helices are energetically stabilized, they are more difficult to unfold (32, 58). Based on prior work (32, 58), the *I*-state is presumed to involve the unfolding of the short *α*-helix associated with residues 61-69.

The blue curve in Fig. 2C is a simple 2-state fit (see SI: Sec. 7) to the level *i* blockade of the 2.0 nm pore using:

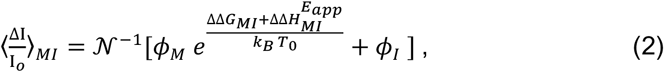

where the fractional blockades for states *M* and, are given by ϕ _*M*_ and ϕ_*I*_, respectively. The normalization is given by 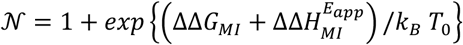 and *T*_*0*_ is room temperature. The results of the fits are summarized in Table 1. Simultaneous use of both terms in Eq. 1 introduces too many free parameters for the fits to converge. However, the two limiting cases result in similar values for the net dipole difference parameter (note that the polarizability difference, Δ*α*_*MI*_, is expressed in dipole units using the midpoint field in Fig. 2C, *E_mid_* = 116.6 MV/m). It should also be noted that, at the mouth of the pore, the fractional blockades are much smaller than when the protein transitions to the *U*-state and moves inside the pore, forming the blockade level *ii*.

To analyze the event durations for each level, we fit the event duration distribution for each state to exponentially decaying distributions. For level *i* (Fig 2D) we find a single time constant of *τ*_*i*_ = 23.2 ± 0.9 ms at −300 mV, which reflects the lifetime at the mouth of the 2.0 nm pore prior to threading via the *U*-state. Beyond −300mV there is a monotonic decrease of *τ*_*ii*2_ (Fig. 2F). In contrast, for the distribution of level *ii* events we observe two distinct time constants, a fast timescale *τ*_*ii*1_ and a slower timescale *τ*_*ii2*_ (Fig. 2E). Interestingly, the duration of *τ*_*ii1*_ (∼0.5 ms) does not appreciably change with voltage (Fig. 2F), and it appears to vanish at voltages beyond −500 mV. This suggests that *τ*_*ii1*_ can be due to transient threading of cyt c into the pore without successful translocation. In contrast, *τ*_*ii2*_ monotonically decreases with increasing voltage beyond −300 mV (Fig. 2F), consistent with the idea that level *ii* corresponds to successful threading and more rapid translocation of the *U*-state as the voltage is increased. The disappearance of *τ*_*ii1*_ at larger voltages suggests a threshold where threading with unsuccessful translocation is apparently eliminated. This observation is correlated with the small decrease in blockade level *ii* above ∼150 MV/m (Fig. 2C). This decrease is assigned to the loss of the transiently and unsuccessfully threaded protein conformation, which contributes to a slightly larger fractional blockade than for a successfully threaded protein.

#### Denaturant and electric field-induced unfolding during cyt c translocation through a 2.5 nm diameter pore

To experimentally verify electric-field induced protein unfolding, we investigated the fractional blockade distributions as a function of voltage for a 2.5 nm diameter pore for different concentrations of guanidinium hydrochloride (Gdm-Cl), a chaotropic denaturant. Notwithstanding previous studies (60, 61) of cyt c at very high (∼1mM) concentrations in 60% ethanol, we observe no evidence of guanidinium-induced aggregation and oligomerization since relatively low fractional blockades are observed, indicating that cyt c is in monomeric state at these very low (0.5-1.0 μM) concentrations. The plotted distributions (Fig. 3A) reveal a clear transition from the native state of the protein to metastable and unfolded states. Below a threshold voltage (<350 mV), we observe exclusively deep blockade ratios (0.8-0.9), attributed to interactions of the native state (N) of cyt c and conformationally excited states *M*. In contrast, for voltages above 350 mV we observe a gradual transition to shallower fractional blockades, indicating further conformational excitation to stretched and unfolded states. We also observed similar behavior in the blockade distributions with respect to voltage for a 3.0 nm pore (SI: Sec.12, Fig. S9), where the transition occurred at much lower voltage (V ∼ −250 mV) as compared to the 2.5 nm pore.

**Figure 3.**
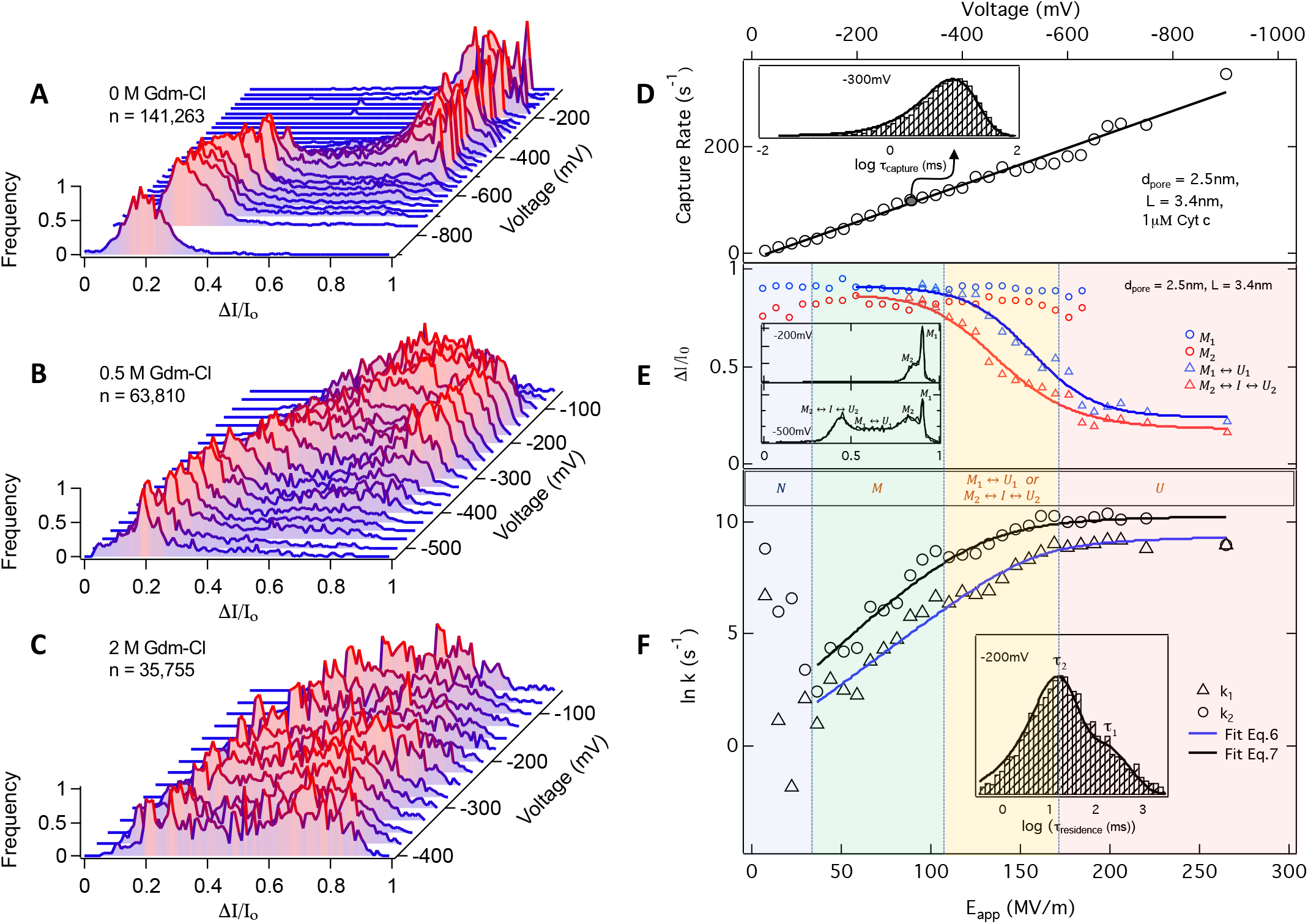
Electric field-induced unfolding of cyt c: Distributions of fractional current blockades as a function of applied voltage measured for pores with d_pore_ = 2.5 nm, L=3.4 nm. **(A)** In absence of Gdm-Cl ([cyt c] = 1 *μ*M), distributions show a clear transition from metastable (higher Δ*I/I*_*0*_) to unfolded (lower AI/I_0_) states. **(B)** In presence of 0.5 M Gdm-Cl ([cyt c] = 0.5 +M), distributions show transitions at a lower voltage, and **(C)** in presence of 2M Gdm-Cl solution (([cyt c] = 0.5 +M), broad distributions without any transition and no well-defined peaks indicate a disordered ensemble of protein conformations. The total number of events collected for each experiment, n, is indicated on each panel. **(D)** capture rates as a function of applied voltage and electric field, when 1 *μ*M cyt c was placed in the cis compartment. The inset shows a typical distribution of log(*τ*_*capture*_) measured at −300 mV and the black solid line represents a single exponential fit to the distribution. **(E)** The mean value of fractional change in current as a function of electric field and applied voltage for different assigned metastable states *(M*_1_, *M*_2_) and dynamically unfolding states *(M*_1_ ↔ *U*_1_, *M*_2_ ↔ *U*_2_). The inset shows typical distributions of fractional change in current, measured at −200 mV and −500 mV, along with the fits using the multicomponent gaussian function. The blue curve is fit with the two-state (*M*_1_ ↔ *U*_1_) model and the red curve is fit with the three-state model *(M*_2_ ↔ *I* ↔ *U*_2_). These fits yield the values of ΔΔ*G*_*MU*_ and Δ*p*_*MU*_ (or Δ*a*_*MU*_, see Table 2 and SI: Sec. 15). **(F)** The rate of protein translocation as a function of applied voltage and electric field. The inset shows a typical distribution of *τ*_*residence*_ measured at −200 mV and the solid curve represents its fit with a bi-modal distribution, yielding two time constants (*τ*_1_ and *τ*_2_) and rates (*k*_1_ and *k*_2_). See SI: Sec.16, Fig. S17 for *τ*_*residence*_ distributions at higher fields. Along with the fractional blockade distributions shown in panel E, this strongly indicates that there are two distinct metastable state configurations (*M*_1_ and *M*_2_) that can squeeze into the pore. Upon increasing the electric field to ∼30 MV/m, the protein is successfully trapped and partially unfolded to form the squeezed *M*-states. As the field increases, transitions take place between the metastable and fully unfolded states (*M*_1_ ↔ *U*_1_ and *M*_2_ ↔ *I* ↔ *U*_2_). The higher field leads to a higher probability of *U*-state formation and much faster translocation. Saturation of fully unfolded protein occurs above ∼200 MV/m.

Next, to confirm that our observations can be attributed to protein shape, we performed experiments in the presence of different concentrations of Gdm-Cl (in 1M KCl, 10mM HEPES, pH7.5) using a 2.5 nm pore (SI: Sec.13, Table S3, Fig. S10-S15). In the presence of 0.5 M Gdm-Cl (Fig. 3B), a transition to unfolded state (low ΔI/I_*0*_) occurs more gradually and begins at lower voltages than in the absence of Gdm-Cl. In contrast, in 2 M Gdm-Cl (Fig. 3C) we observe no clear population and no transition, indicating that random portions of the protein are more likely to enter the pore due to its disordered state (data for 1 M Gdm-Cl are presented in SI: Fig. S12). Further, in 3 M Gdm-Cl (SI: Sec.13, Fig. S13, S14) we find a clear peak (around 0.3) in the fractional blockade distribution, attributed to fully unfolded cyt c conformations.

**Table 2:**
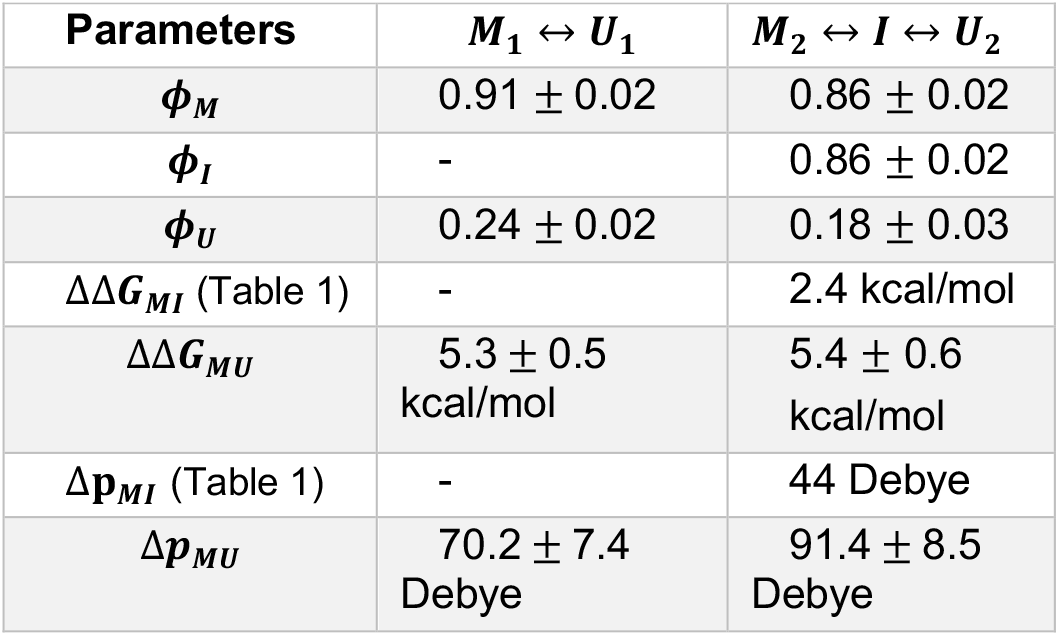
Fit parameters from the experimental data (d_pore_ = 2.5 nm, L = 3.4 nm) in Fig. 3E using two-state and three state models (SI: Sec. 15, Eqs. S8 and S7, respectively).

#### Squeezing and electric field-induced unfolding of distinct metastable states during cyt c translocation

Next, we examine the detailed process of cyt c passage through a 2.5 nm pore as a function of voltage, or electric field (*E*_*app*_). In Fig. 3D we plot the mean capture rates as a function of *E*_*app*_. Our finding of a linear increase in capture rate is consistent with drift-limited capture theory (24). Further, this observation, in addition to long measured residence times (relative to our 10 μs measurement time resolution), rule out the possibility of any time resolution artifacts in the entire voltage range of our experiments.

Analysis of the fractional blockade distributions reveals a total of four distinct populations, and they were fit using a four-component Gaussian where the centroids identify various states of cyt c in the pore, as plotted in Fig. 3E for each *E*_*app*_ (see also SI: Sec.14, Fig. S16). We observe two peaks centered around 0.82 (red markers) and 0.9 (blue markers), which vanish for *E*_*app*_ *≳* 200 MV/m. Between *E*_*app*_ ∼0-30 MV/m there is a gradual transition from initial native states N to squeezed metastable states, designated as *M*_*1*_ and *M*_*2*_, which may differ from each other by how these partially unfolded configurations are able to squeeze into the 2.5 nm pore. As *E*_*app*_ is increased into the range ∼110-170 MV/m, large subsets of the two populations undergo a major dynamic unfolding transition as evidenced by a significant reduction in the fractional blockades (triangles in Fig. 3E). These transitions are designated as *M*_*2*_ ↔ *U*_*2*_ (blue triangles) and *M*_*1*_ ↔ *U*_*1*_ (red triangles). Finally, as *E*_*app*_ approaches ∼250 MV/m, the protein transitions to a completely unfolded state, which is marked by very low fractional blockades. We envision that the unfolded protein has two types of blockade states (denoted *U*_1_ and *U*_2_) within the pore depending upon the initial squeezing configurations and the ensuing unfolding pathway (*M*_*1*_ ↔ *U*_*1*_ or *M*_*2*_ ↔ *I* ↔ *U*_*2*_).

#### Energetics of cyt c unfolding

To quantitatively understand the observed electric field-induced unfolding transition and to evaluate the energy gaps and dipole interactions of the various states, we have considered both a simplified 2-state model involving conformational states (*M* and *U*), separated by an energy gap ΔΔ*G*_*MU*_ as well as a sequential 3-state unfolding model, which also includes the *I*-state. Both of these models lead to expressions that are analogous to Eq. 2 and they are discussed more completely in the SI: sec 15. Because the *M*_*1*_ ↔ *U*_*1*_ transition shows a much broader distribution compared to the *M*_*2*_ ↔ *U*_*2*_ transition (inset Figure 3E and SI: Sec.14, Fig. S16), we assume that the *M*_*1*_ ↔ *U*_*1*_ distribution of blockade states is related to a more direct transition from *M*_*1*_ to a random set of unfolded conformations *U*_*1*_, which translocate at differing rates centered around the mean value. This suggests a direct, non-sequential, unfolding pathway between *M*_*1*_ and *U*_*1*_ exists within the pore. For the *M*_*2*_ ↔ *U*_*2*_ transition, we assume a sequential pathway where one of the *α*-helices unfolds first, forming the *I*-state, followed by the other two *α*-helices to form the *U*-state (32, 58). Thus, the blue triangles, which are a measure of the mean of the *M*_*1*_ ↔ *U*_*1*_ blockade distribution, are fit using a simple two-state *M*_*1*_ ↔ *U*_*1*_ model (SI: Sec. 15, Eq. S8), which yields 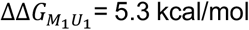, (Table 2). On the other hand, the mean of the blockade distribution for *M*_*2*_ ↔ *U*_*2*_ (red triangles Fig. 3E) is described using a three-state *M*_*2*_↔*I* ↔ *U*_*2*_ sequential unfolding model (SI: Sec.15, Eq. S7), which is constrained by the *M* ↔ *I* parameter values found previously (Table 1). This approach yields a similar transition energy between *M* and *U* (i.e., 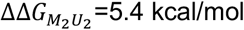, Table 2).

Thus, both the *M*_*1*_ ↔ *U*_*1*_ and *M*_*2*_ ↔ *U*_*2*_ free energy gaps are found to be in excellent agreement with the value of the total free energy, 5.4 kcal/mol, previously found (32) for unfolding the three major *α*-helical regions of cyt c. It is also worthwhile to mention that a threshold voltage required to rupture a protein-DNA complex using a solid-state nanopore was previously found to correlate well with the equilibrium free energy of the complex formation (62). Finally, we note that the initial breakage of the salt bridge, involving E62, and the partial unfolding that leads to formation of the squeezed *M*-state, should leave the free energy gap between *M* and *U* unaffected. This is verified by independent studies using a E62G mutation to remove the salt bridge, which demonstrated that the ΔΔ*G*_*MU*_ energy gap remains unchanged (32, 58).

#### Kinetics of cyt c translocation

In order to discern the kinetics of translocation of metastable and unfolded states, we analyzed the rate *k*_*tr*_ = 1/⟨τ_*residence*_⟩ of protein translocation as a function of voltage and electric field (for *E*_*app*_ *≳* 30 MV/m) where we again make use of the fast *α*-helical folding/unfolding transitions (51-56) relative to the experimental (∼ms) translocation times. Thus, the thermodynamically averaged translocation rates for the two and three state models can be expressed as:

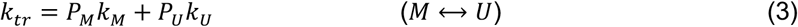

or

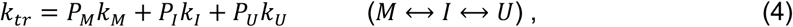

where *P*_*i*_ is the *E*_*app*_ dependent probability of finding cyt c in the *i^th^* state, and *k*_*i*_ is the *E*_*app*_ dependent translocation rate of the *i*^*th*^ state.

For the *U*- and *I*-states, we assume the field dependent translocation rates are given by:

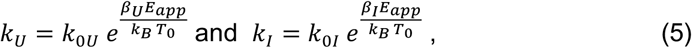

with 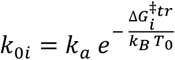. The Arrhenius prefactor, *k*_*a*_, is estimated below and, for simplicity, we assume that *k*_*M*_ ≅ 0 for the 2.5 nm pore. The unknown parameters, *β*_*U*_ and *β*_*I*_, account for the translocation barrier reduction due to the electrophoretic forces. This parameter can be combined with the state dependent dipole difference values (Eq. 1), associated with *p*_*i*_, and obtained from the fits to the blockade ratio analysis (Table 2). Thus, upon combining, we can define 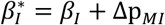 and 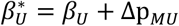, which become the final fitting parameters for analysis of the translocation rates. When the zero-field free energy gaps are constrained by the values found in Table 2, the expression for the translocation rate in the two-state model depends only on *k*_o*U*_ and 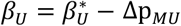 (see SI: section 17):

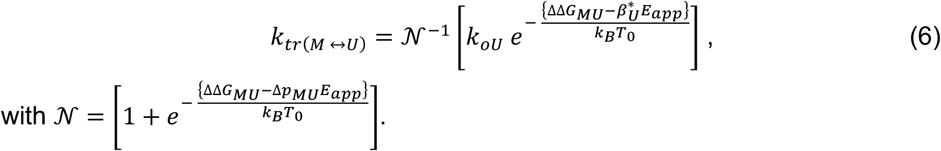

For the three-state model we have:

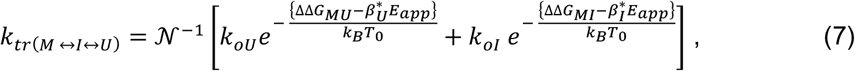

with

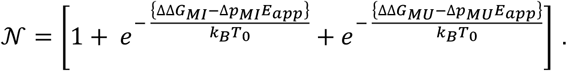

As shown in the plot of *ln* [*k*_*tr*_] (Fig. 3F), after trapping and conversion to the squeezed *M*-state, there is an increase in translocation rate as a function of the electric field above ∼30 MV/m. Additionally, at each voltage, we observed two distinct rate distributions (inset Fig. 3F and SI: Sec.16, Fig. S17), which we attribute to the translocation of two distinct conformationally excited states as noted in the caption of Fig. 3F.

Below 30 MV/m, *1*/⟨*τ*_*residence*_⟩ decreases with voltage and we attribute this behavior to the increased time spent by the N-state of cyt c as it is trapped at the pore mouth. At these low electric fields, the energy of the excited states on the unfolding pathway of cyt c are not lowered sufficiently to allow thermal excitation from) to *M*. However, in the regime between 30-100 MV/m, *ln*[*k*_*tr*_] starts to increase with the electric field, manifesting the conformational excitation to the *M*-state and its rapid thermodynamic exchange with the *I*- and *U*-states which also have a reduced free energy gaps due to the electric field interaction. Consistent with the blockade ratio results presented above, we assigned two distinct rates in this regime to the two metastable states, *M*_*1*_ and *M*_*2*_, differing by a small amount, presumably due to different trapping configurations of the *M*-state within the pore. Between 100-170 MV/m, the increase of *ln*[*k*_*tr*_] begins to diminish, while above 170 MV/m the rate levels off. This is attributable to the saturation of the *U*-state population (and its associated distribution of blockade ratios and translocation times) as the electric field is increased.

The fitting parameters for the solid dark lines in Fig. 3F, found using Eqs. 6 and 7, are listed in Table 3. It can be seen that the *U*-state has the dominant rate for translocation compared to the *I*-state (and the *M*-state rate was assumed to be small enough that it was neglected in the field-dependent kinetic analysis). Moreover, *β*_*U*_, which governs the field dependence of the *U*-state translocation rate, is seen to be negligible compared to the dipole difference, suggesting that the electrophoretic work done by the field, which lowers the *U*-state translocation barrier, is relatively small for the 2.5 nm pore. However, it should be noted that *β*_*1*_= 47 Debye is much larger than *β*_*U*_ = 2-3 Debye, suggesting there is much more resistance to the electrophoretic forces acting to pull the *I*-state protein through the 2.5 nm pore. This is also evidenced by its much slower zero-field rate, *k*_*01*_=72 s^-1^, compared to *k*_0*U*_ =10712 s^-1^. Additional analysis of the 2.0 nm pore data (e.g., Fig. 2E) in the SI (Sec. 18, Fig. S19) yields a low field slope indicating that *β*_*U*_ for the 2.0 nm pore is about an order of magnitude larger than found for the 2.5 nm pore, while the intercept at zero field suggests *k*_0*U*_ ∼5 s^-1^ for 2.0 nm pore.

**Table 3:**
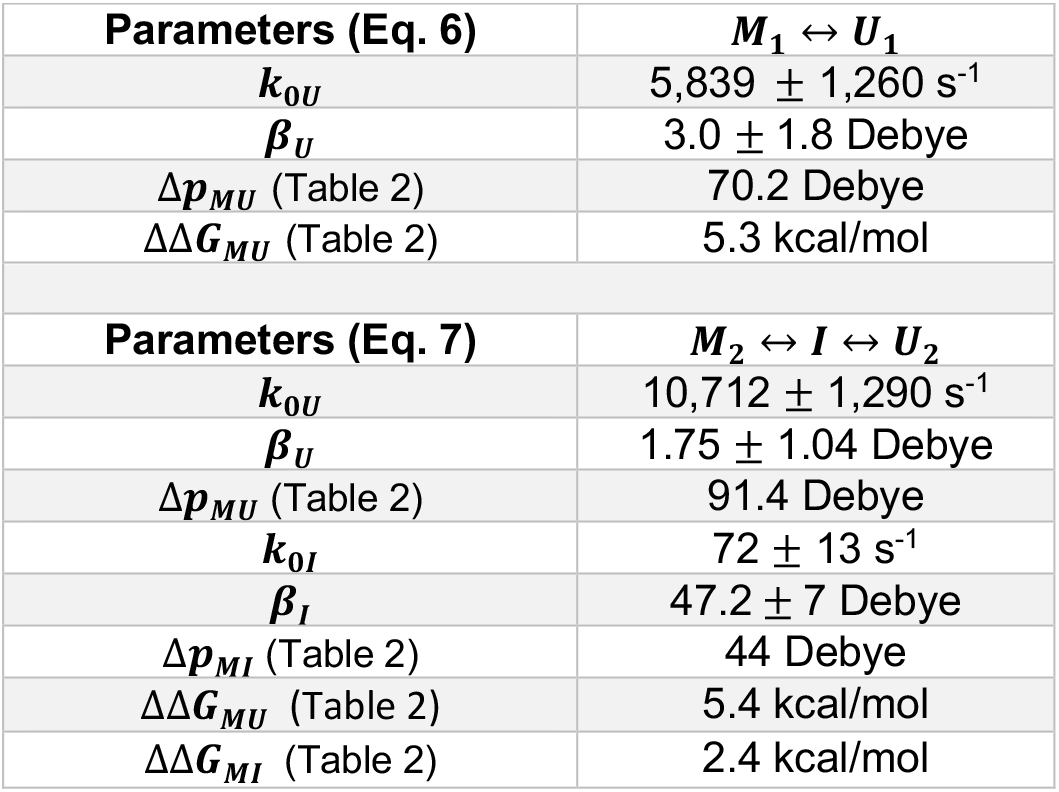
Parameters from the fit of kinetic data (Fig. 3F) using Eq. 6 for k_1_ (triangles) and Eq. 7 for k_2_ (circles).

As can be seen in Table 3, there is a dominant *k*_0*U*_ translocation rate extracted for the two *U*-state configurations. These separate rates derive from the different configurations of their parent states, *M*_*1*_ and *M*_*2*_. To evaluate the translocation barriers for 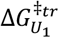 and 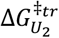, we can use a simple Arrhenius equation for the rate in zero field:

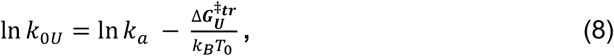

where *k*_*a*_ is the rate of translocation at zero electric field without any activation barrier (i.e., no pore). This can be calculated from the rate at which cyt c moves a distance comparable to the length (L) of the pore. Thus, assuming Stokes-Einstein diffusion of cyt c in bulk solution,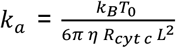 where η is the viscosity of water (1 centipoise) and R_*cyt c*_ is the hydrodynamic radius of cyt c (∼1.5 nm). Using these values and L =3.4 nm, we find *k*_*a*_ = 1.26×10^7^ s^-1^. The values for *k*_0*U*_ in Table 3 for the 2.5 nm pore, along with Eq. 8 then lead to: 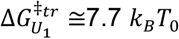 and 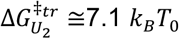. Using the *I*-state rate, *k*_0*I*_ =72 s^-1^, we can similarly deduce that 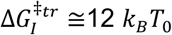. Finally, if we extrapolate the triangle data points in Fig. 3F to zero electric field and attribute this to *M*_*1*_ translocation, we can estimate a zero-field rate 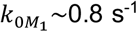. Similarly, by extrapolating the circle data points in Fig. 3F to zero-field, we can estimate 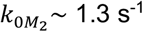. Thus, by using Eq. 8, we can deduce 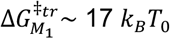 and 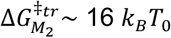.

#### MD simulations

To confirm that electric field alone can indeed produce unidirectional translocation of cyt c protein through a solid-state nanopore and to relate protein unfolding to nanopore diameter and changes in the ionic current blockade levels, we performed all-atom MD simulations of six nanopore systems, varying in diameter from 5.5 to 1.5 nm, Fig. 4A, SI: Sec.11, Table S2, and Fig. S7. Each simulation system contained one Si_3_N_4_ nanopore, one cyt c protein placed in front of the nanopore and 1 M KCl solution. Following equilibration, each system was simulated using the G-SMD protocol (63), where the open-pore electrostatic potential map, amplified by a scaling factor, was used to drive the translocation of the cyt c protein with an accelerated rate. To eliminate the uncertainty associated with various paths the protein could take to enter the nanopore, the protein was additionally restrained to have its center of mass located along the pore axis. The simulations were carried out at effective biases of 1, 2 and 3V, which allowed us to observe complete permeation events for the majority of the nanopore systems within a 100 ns time scale. Such a dramatic acceleration of the translocation process is expected as the rate of a forced barrier crossing exponentially depends on the magnitude of the applied force (64). Fig. 4B and C; Fig. 4D—F summarize the simulation results.

**Figure 4.**
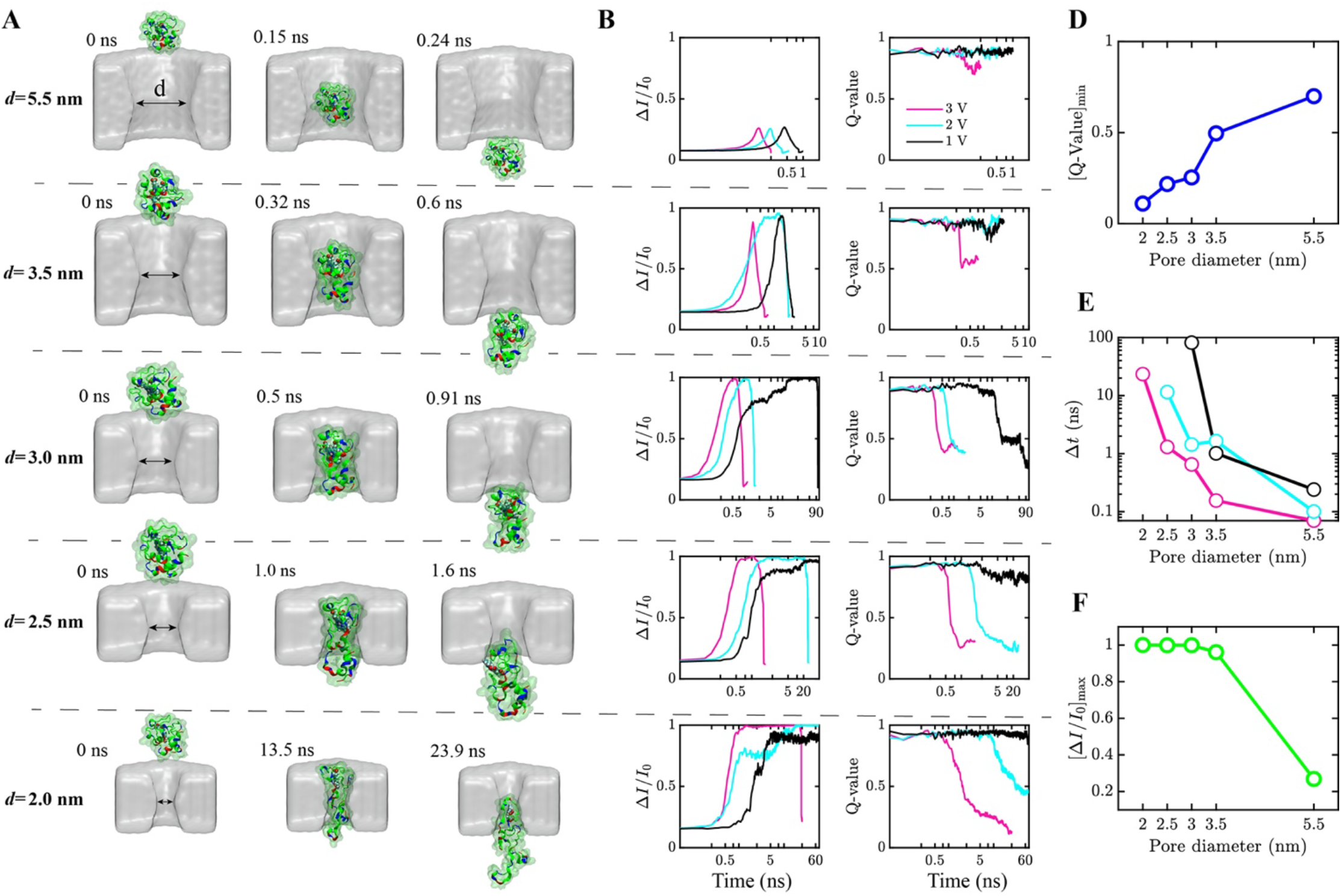
Molecular dynamics simulations of cyt c passage through nanopores. **(A)** Snapshots illustrating five MD trajectories where a single cyt c protein (shown as a carton enclosed by a semi-transparent surface) was forced to pass through a nanopore (gray) of specified diameter *d* using the G-SMD protocol (SI: Sec. 11(iii)) under a 3V effective bias. **(B, C)** Ionic current blockade (B) and the fraction of native protein contacts, Q-value (C), versus simulation time for three G-SMD simulations carried out at the effective biases of 1 (black), 2 (cyan) and 3 (pink) V. The pore diameter *d* was, from top to bottom, 5.5, 3.5, 3.0, 2.5, and 2.0 nm. The ionic currents were computed using the SEM approach (SI: Sec. 11 (iv)) Note the logarithmic scale of the time axis. **(D-F)** Minimum Q-value (D), nanopore translocation time (E, logarithmic scale) and maximum blockade current amplitude (F) observed during the MD simulations as a function of the pore diameter.

We found the pore diameter to have a pronounced effect on the time scale and the character of cyt c translocation. When the pore was slightly larger (*d* = 5.5 nm) than the protein, the translocation proceeded very fast (< 1ns) and produced well-defined shallow blockades of the nanopore current, Fig. 4A-B. The protein structure, which we characterize here as the fraction of native contacts remaining, i.e., the Q-value (65), remained largely unperturbed by the nanopore passage, Fig. 4C. The translocation through a nanopore that was slightly smaller (*d* = 3.5 nm) than the protein proceeded via reversible squeezing of the protein that largely preserved the secondary structure. The protein was found to block nearly 90% of the ionic current when confined to the pore constriction. A qualitatively different behavior was observed for 3.0, 2.5 and 2.0 nm diameter pore: the protein was found to partially unfold during the translocation, with the average degree of unfolding, Fig. 4D, decreasing with larger pore diameters. The average translocation time was found to exponentially depend on the pore diameter, Fig. 4E, whereas the maximum current blockade ratio (computed using SEM approach (66)) during the translocation, Fig. 4F, reached nearly 100% of the open pore current. Interestingly, in our simulations we did not observe cyt c passage through the 1.5 nm diameter nanopore, unless the protein was mechanically unfolded by the application of external force to a protein terminus (SI: Sec.11, Fig. S8).

### Discussion

One striking feature of our analysis is that for a 2.5 nm pore the values of 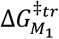 *and* 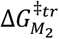 (estimated by use of Eq. 8 and straight line extrapolation of the rates to zero electric field as in SI: Sec.19, Table S7) are less than that of the unfolding barrier (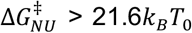 with *k*_*B*_*T*_*0*_ = 0.59 kcal/mol) reported in an earlier study (32). Thus, squeezing of cyt c via metastable conformations *M*_*1*_ and *M*_*2*_ into the pore is an energetically favorable path compared to the requirement that the protein completely unfolds and translocates via threading. The value of 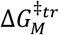 is also clearly greater than 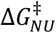 in the experiments with the 1.5 nm and 2 nm pores (SI: section 18, Fig. S19). Thus, for the smaller pores, excitation from *M* to a higher energy conformational state, *I*, at the pore mouth, followed by unfolding mediated threading of the *U*-state is the favorable path for translocation.

Another remarkable point is that the values of ΔΔ*G_MI_* = 2.4 kcal/mol (Table 1), 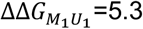, and 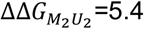 (Table 2) obtained from the analysis, are in very close agreement with the cyt c *α*-helix unfolding transitions reported in previous work (32, 58). In the prior studies, a 2.6 kcal/mol free energy gap was found between a partially unfolded state near 7.4 kcal/mol (which we identify as the *M*-state with the three main *α*-helices still folded), and the next higher state on the unfolding pathway (which we identify as the, *I*-state) where the *α*-helix associated with residues 61-68 has unfolded. Moreover, the free energy gap from *M* to the fully unfolded state *U* was found in prior work (32) to be 5.4 kcal/mol, almost exactly the value found for both of the configurations (*M*_*1*_ and *M*_*2*_) within the pore.

In summary, we have demonstrated protein translocation through a pore barrier (d_pore_ < d_protein_) without any requirement of chemical denaturants, an enzymatic motor, or an oligo tag. We achieved this by electric field manipulation of the conformational energy levels of a protein in the vicinity of the pore. We find that cyt c is able to squeeze into the 2.5 nm pore where it can still undergo rapid conformational transitions between folded and unfolded *α*-helical states. The protein then translocates through the pore with a rate that depends upon the thermodynamic probability to completely unfold the residual *α*-helices. In contrast, for 2 nm and 1.5 nm pores, cyt c undergoes conformational excitations at the mouth of the pore first from state *M*, with all three *α*-helices intact (32, 58, 59), to state *I*, with only a residual bi-helical segment (32, 58, 59) and finally to the fully unfolded state so that entry and translocation occurs only via threading through the pore. These results demonstrate that the necessity of complete unfolding for a protein to enter and translocate depends upon the diameter of the pore. The relatively large field strengths (∼10^7^-10^8^ V/m), needed for translocation in these experiments, owe to the more rigid nature of the SiN pore as compared to biological membrane pores, where typical electric field values that are used in translocation measurements are lower (<10^7^ V/m, or 50 mV/5 nm).

The experiments reported here set the stage for new studies where electrical manipulations sculpt the conformational landscape of proteins. Detection of protein signatures and folding patterns as they pass in unique ways through a nanopore can also serve as the basis of their identification, and this approach requires only a localized electric field without any additional requirements such as enzymes, oligo tags, or chemical denaturants. Future studies will examine the behavior of other proteins using similar experiments and comparisons among protein structural variants and proteins with and without post-translational modifications.

### Materials and Methods

We assembled the solid-state nanopore chips in a flow cell that separates two compartments *cis* and *trans* each containing 1 M KCl, 10 mM HEPES (pH 7.5) buffer solution (see Fig. 1A). To observe cyt c interactions with a pore, we placed 0.5 or 1.0 *μ*M of cyt c (1 M in absence of Gdm-Cl and 0.5 *μ*M in presence of Gdm-Cl) in the *cis* compartment, applied a negative potential (−25mV to −900 mV) to the *trans* compartment while keeping *cis* grounded. This generates a strong pore-localized electric field (∼10^8^ V/m) oriented in the *cis* to *trans* direction, which results in ion current that is subject to modulation when cyt c is trapped or enters the pore.

We used previously reported wafer fabrication methods (30, 50, 67, 68) to drill ultrathin solid-state nanopores in the diameter range of 1.5 - 5.5 nm in high-stress negatively-charged SiN membranes (SI: Sec .3, Fig. S2). Based on the measured conductance across the pores (SI: Sec. 4 Table S1) we used a standard procedure (67-69) to obtain the effective pore length L, as indicated in the figure captions. Additional chemical materials, experimental methods are described in SI: Sec. 1 and Sec. 2 and MD simulation methods are described in SI: Sec. 11.

## Acknowledgments

PT would like to thank Mehrnaz Mojtabavi for assistance with the gel electrophoresis experiments and providing access to the lab computer during COVID-19 outbreak. This work was supported by the National Science Foundation through grants DMR-1710211 (MW), DMR-1827346 (AA) and CHE-1764221 (PMC). Computer time was provided through the Extreme Science and Engineering Discovery Environment (XSEDE) allocation MCA05S028.

## Supplementary Information

## 1. Materials and methods (Experimental)

We used ultrathin high-stress SiN (250 MPa) membranes supported by a Si chip as substrates for nanopore fabrication(1-3). Nanopores were cleaned in hot piranha (2:1 H_2_SO_4_ / H_2_O_2_) for 30 minutes, followed by hot deionized water, before each experiment. After cleaning, nanopore chips were assembled in a custom flow cell equipped with Ag/AgCl electrodes, and a quick-curing silicone elastomer was applied between the chip and the cell to seal the device and thereby reduce the noise by minimizing the chip capacitance.

The purity of equine heart cyt c (Sigma Aldrich C2506) was confirmed by SDS PAGE gel electrophoresis (Fig. S1(ii)). All experiments were carried out at ambient temperature. The apparent electric field *(E*_*app*_) inside the pore were obtained by ratio of applied electric potential in *trans* chamber and length of the pore and was used to interpret the data.

### SDS-PAGE Gel Electrophoresis

Preparation of Gel: We added 4.9 ml of deionized water to a 50 ml cylindrical tube, then added 6 ml of 30% acrylamide mix, 3.8 ml of 1.5M tris (pH 8.8), 0.15 ml of 10% SDS, 0.15 ml of 10% ammonium persulfate, 0.006 ml of TEMED. The resultant solution was shaken and added to the gel-glass-plate. A comb was inserted to generate the well structures, and a 30 minute waiting time allowed polymerization of the gel.

10X Running Buffer: We dissolved 30.0 g of tris base, 144.0 g of glycine, and 10 g of SDS in 1000 ml of H_2_O. The pH of the buffer was observed to be 8.3 without any adjustment. The resultant solution was stored at room temperature and diluted to 1X before use for gel electrophoresis experiments.

Running the Gel: We loaded the gel-glass-plate in a vertical electrophoresis cell, and then added 1X running buffer. We heated the cyt c solution (1 mg/ml) at 95°C for 5 minutes in heat block and then 10 *μ*L of cyt c was added to one well, alongside another well that was loaded a molecular weight marker. Voltage was set to a fixed value of 150 V, and the gel was allowed to run for 45 minutes.

### Electrical detection and data acquisition

The ionic current through nanopores was measured using an Axopatch 200B amplifier (Molecular Devices) and low-pass filtered to indicated bandwidth using the internal Bessel filter of the Axopatch. Data points were digitized and sampled at 250 kHz sample rates on a National Instruments DAQ card using custom LabVIEW software. For the 5.5 nm pore we have performed high-bandwidth measurements of ionic current using a Chimera instruments VC100 amplifier (4). Data were processed and events were detected using Pythion (https://github.com/rhenley/Pyth-Ion/) and multilevel events were detected using a custom algorithmic procedure in Igor Pro software (as described below).

### Algorithms for multilevel detection

The algorithms developed here for multilevel detection in a typical nanopore trace fall under the umbrella of change point detection techniques in a time series. Here, we are mainly concerned with changes of the mean of the distribution; and below we briefly describe two algorithms that that have been used throughout this paper:

#### (a) Two sliding windows (TSW) algorithm

This method uses two sliding consecutive windows each with *n* datapoints all having the same y value corresponding to the mean of the trace within the window; which is denoted *w*_1_ for the first window and *w*_2_ for the second. The user chooses 2 datapoints on a continuous segment of baseline bn_1_ and bn_2_. These two data points are used by the program to calculate the mean and the standard deviation of the baseline denoted as *baseline* and *σ*, respectively. We then define a threshold, *th*, for change point detection from baseline to event and vice versa. *th* is defined as: *th* = *n*_*th*_*σ*; with *n*_*th*_ typically varies from 4 to 10. In most cases biomolecules rotate and change conformation at the mouth of or within the pore. These rotational and conformational motions superimpose with the intrinsic noise of the baseline and increase the apparent noise of the levels of an event. To deal with this situation we define a level threshold, *th*_*l*_, that is typically larger than *th*.

The two windows slide and scan the trace with steps of n datapoints using a for loop and at each step we check for the following conditions:

i. If (*baseline—w*_1_ < *th*)*and*(*baseline* — w_2_ > *th*): this indicates the end of baseline, start of an event and start of the first level of the event. The segment of the baseline that has ended is fitted with the mean of its datapoints.
ii. Elseif (*baseline — w*_1_ > *th*)*and*(*baseline — w*_*2*_ > *th*)*and*(*abs*(*w) — w*_*2*_) > *th*_*l*_): this indicates the end of a level and the start of another within an event. The segment of the level that has ended is fitted with the mean of its datapoints.
iii. Elseif (*baseline —w*_1_ > *th*)*and*(*baseline — w*_*2*_ < *th*): this indicates the end of an event and its last level and the start of the baseline. The segment of the last level that has ended is fitted with the mean of its datapoints.

The levels are indexed as *(i,j)*, where *i* is the index of the event and *j* is the index of the level within the event.

#### (b) Extending window and sliding window (EWSW) algorithm

This method uses two consecutive windows, an extending window (w_1_) which represents the cumulative mean of all the datapoint after the last change point and a sliding window (*w*_*2*_) with *n* datapoints all having the same y value corresponding to the mean of the trace within the window. The steps of this algorithm are the same as the TSW algorithm with the only exception is the change of the meaning of *w*_1_.

To speed up the calculation we do not calculate the cumulative mean each time by taking the average of the previous datapoint and we use the following trick instead: at each step k the cumulative sum *cusum* and cumulative number of points *cuN* are updated by calculating them recursively using the following formulas:

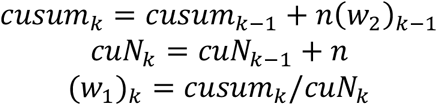

Both techniques fit multilevel traces well; however, in some cases the EWSW algorithm performs better than TSW. Therefore, all the traces in this paper are fit using EWSW algorithm. More details about the performance of these algorithms will be discussed in a separate paper.

## 2. Sequence and gel electrophoresis of cyt c used in our experiments

**Figure S1(i).**
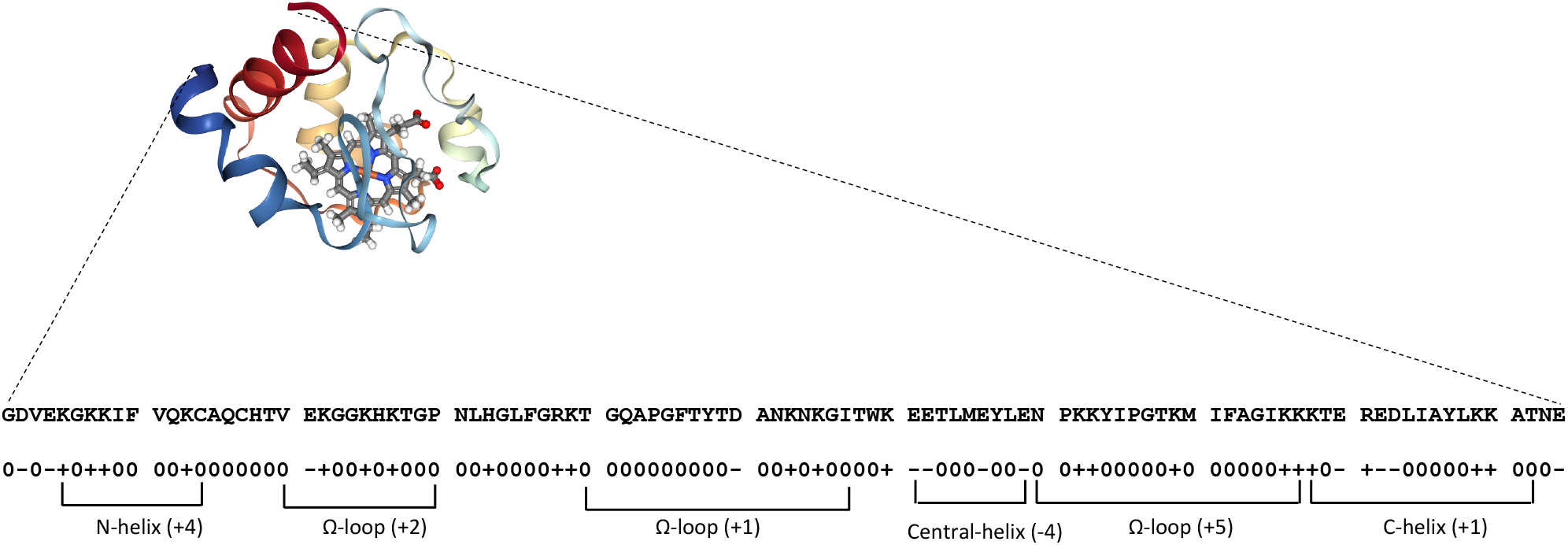
Sequence of equine heart cyt c used in the experiments and a segmental representation of the charge distributions.

**Figure S1(ii).**
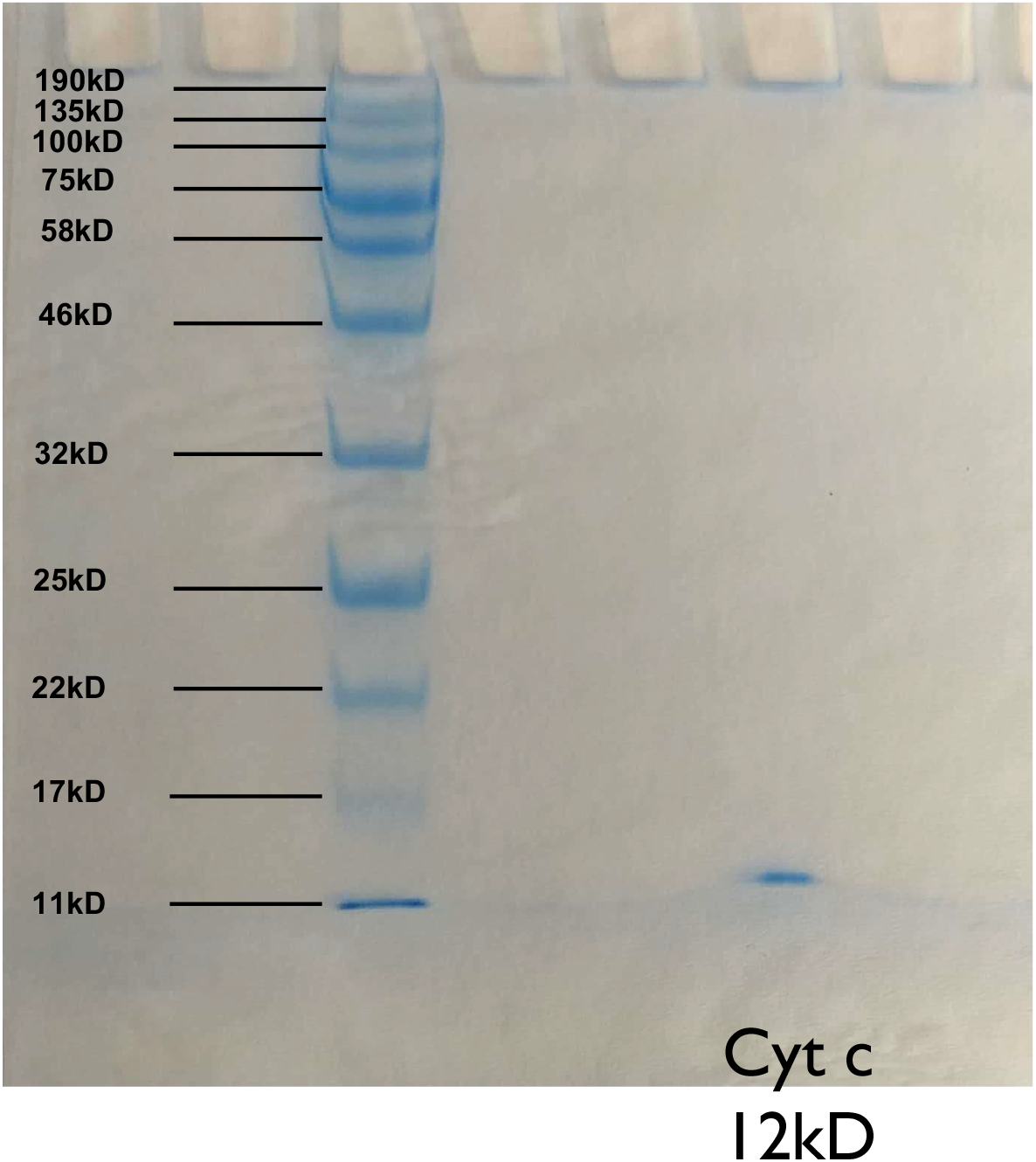
Gel electrophoresis of cyt c (equine heart) used in the experiments. A clear single band near 12 kD indicates the purity of the sample, and confirms that no covalent (e.g., disulfide-bridged) dimers are formed.

## 3. Surface charge measurements of a high-stress SiN nanopore

**Figure S2.**
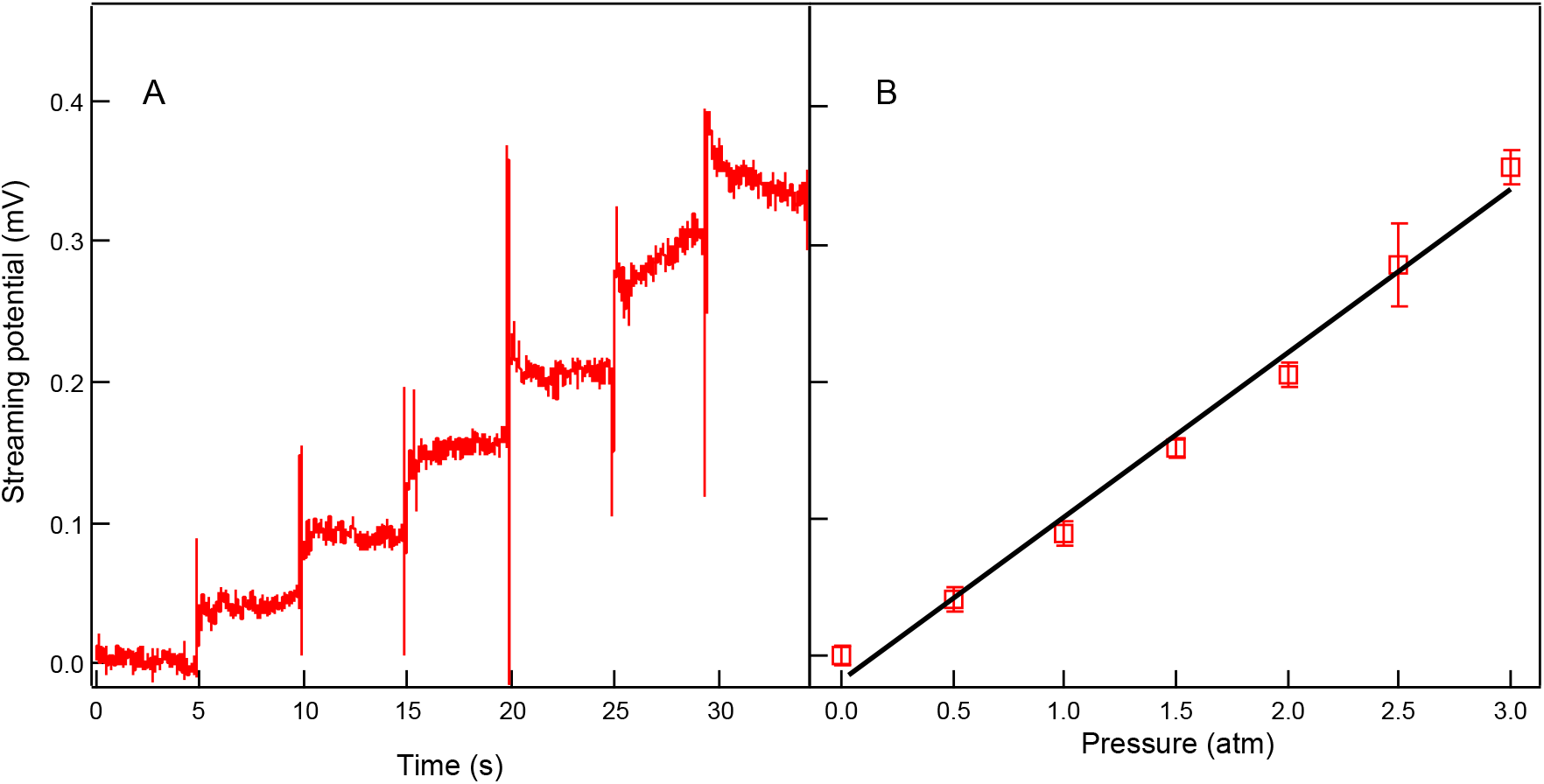
Example surface charge measurement of a high-stress (250 MPa) SiN pore. A) Streaming potential current trace of 6.1 nm pore in 0.4 M KCl, 10 mM Tris, 1 mM EDTA pH 7.8. Using automated pneumatic pressure controller, streaming potential was measured by applying 0.5 atm increments every 5 seconds. B) Streaming potential *vs*. pressure. Surface charge of SiN pore (−7.24 mC/m^2^) was calculated from the obtained ζ-potential value (−5.01 mV), which is derived from the linear fit to the streaming potential *vs*. pressure data. Error bars in panel B represent the ± standard deviation from the mean streaming potential data.

Fig. S2(A) shows 30-sec streaming potential trace of 6.1 nm diameter SiN pore with increment of 0.5 atm in 0.4 M KCl, 10 mM Tris, 1 mM EDTA, pH 7.8. The air pressure is applying on cis chamber using the automated pneumatic pressure scanner. The average streaming potential as a function of applied pressure is plotted in Fig. S2(B). As presented in our previous work (5, 6), the zeta potential is expressed as the following equation;

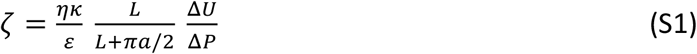

where *η, K, ε, L, a, U, P* are viscosity of solvent, solvent conductivity, solvent dielectric constant, thickness of pore, radius of pore, streaming potential, and applied pressure, respectively. The value of ΔU/ΔP obtained from the slope of linear fit in Fig. S2(B) yields *ζ* = −5.01±0.25 mV. When the electrical potential 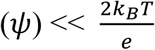, the surface charge (*σ*) is;

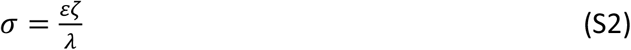

where *λ* is Debye length, which is calculated by 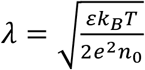. Using *ζ*-potential value, the 2e surface charge of silicon nitride is −7.24 ± 0.36 mC/m^2^. The extraction of zeta potential in the limit of dU/dP as P→0 by a quadratic fit of the data yields *ζ* = −3.22 ± 0.26 mV and surface charge of silicon nitride is −4.65 ± 0.38 mC/m^2^. On the other hand, using linear fit of the first three data points yields *ζ* = −3.76 ± 0.19 mV and surface charge of silicon nitride is −5.43 ± 0.27 mC/m^2^. The different protocols for fitting the data shown in Fig. S2(B) lead to small variations in the surface charge that only weakly perturb the overall applied force on the molecules (given that transport is an electrokinetic phenomenon that has both electrophoretic and electroosmotic effects, and given the ultrathin nanopore shape). This has a negligible effect on the MD simulations because of the large driving forces that are applied. The fitting protocol has no significant effect on the analysis, interpretations and conclusions presented in the main manuscript.

## 4. Open pore conductance for different pore diameters

**Table S1:** Conductance values for different pore diameters: The conductance values were obtained from the slope of current vs voltage data (see Fig. S21 in Sec. 20) and were used to estimate the effective pore length L as described in previous reports (1, 2, 7). Error values in the conductance represents the errors obtained in the slopes of the current vs voltage data when fitted with a straight line (Fig. S21). The nanopores were fabricated and their diameters measured using TEM (transmission electron microscopy).

| Pore Diameter (nm) | Conductance (nS) | Pore Length (nm) |
| --- | --- | --- |
| 5.5 | $23.02 \pm 0.07$ | 7.1 |
| 3.5 | $12.55 \pm 0.02$ | 6.0 |
| 3 | $11.78 \pm 0.05$ | 4.3 |
| 2.5 | $10.08 \pm 0.02$ | 3.4 |
| 2 | $6.13 \pm 0.04$ | 4.1 |
| 1.5 | $5.28 \pm 0.04$ | 2.5 |

## 5. Electric field and dipole orientations

If the minimization of the potential energy (*-p*.*E*) due to alignment of the electric dipole (*p*) of cyt c along the electric field (*E*) is greater than the thermal energy 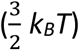, then cyt c would begin to trap in an orientation along the electric field. For 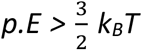, the requirement is *E* > *1*.*5 k*_*B*_*T/p*. Putting in literature values (8) of *p* =320 × 0.0208 e-nm and *k*_*B*_*T* = 25.7 meV, suggests that in an electric field *E* > 5.8 mV/nm, cyt c will begin to orient itself along the electric field.

## 6. Representative current traces for cyt c using a 2.5 nm pore at different voltages

**Figure S3(i).**
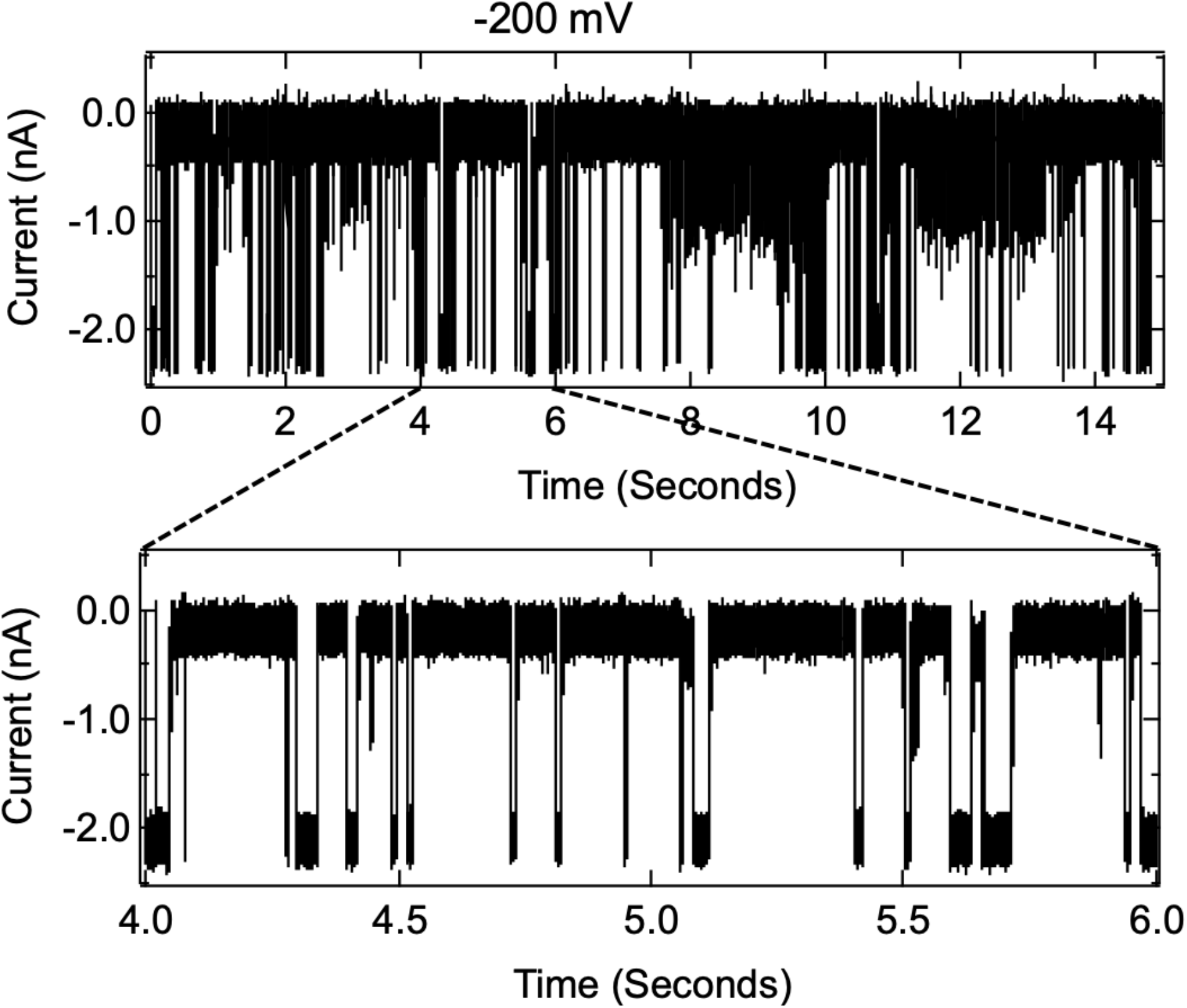
Ionic current traces for cyt c using a 2.5 nm pore at −200 mV applied voltage. The measurements were carried out in 1M, KCl, 10mM HEPES, pH 7.5, sampling rate of 250 kHz, and filtered using a low-pass Bessel filter of 100 kHz. Open pore current value is ≅ −2 nA.

**Figure S3(ii).**
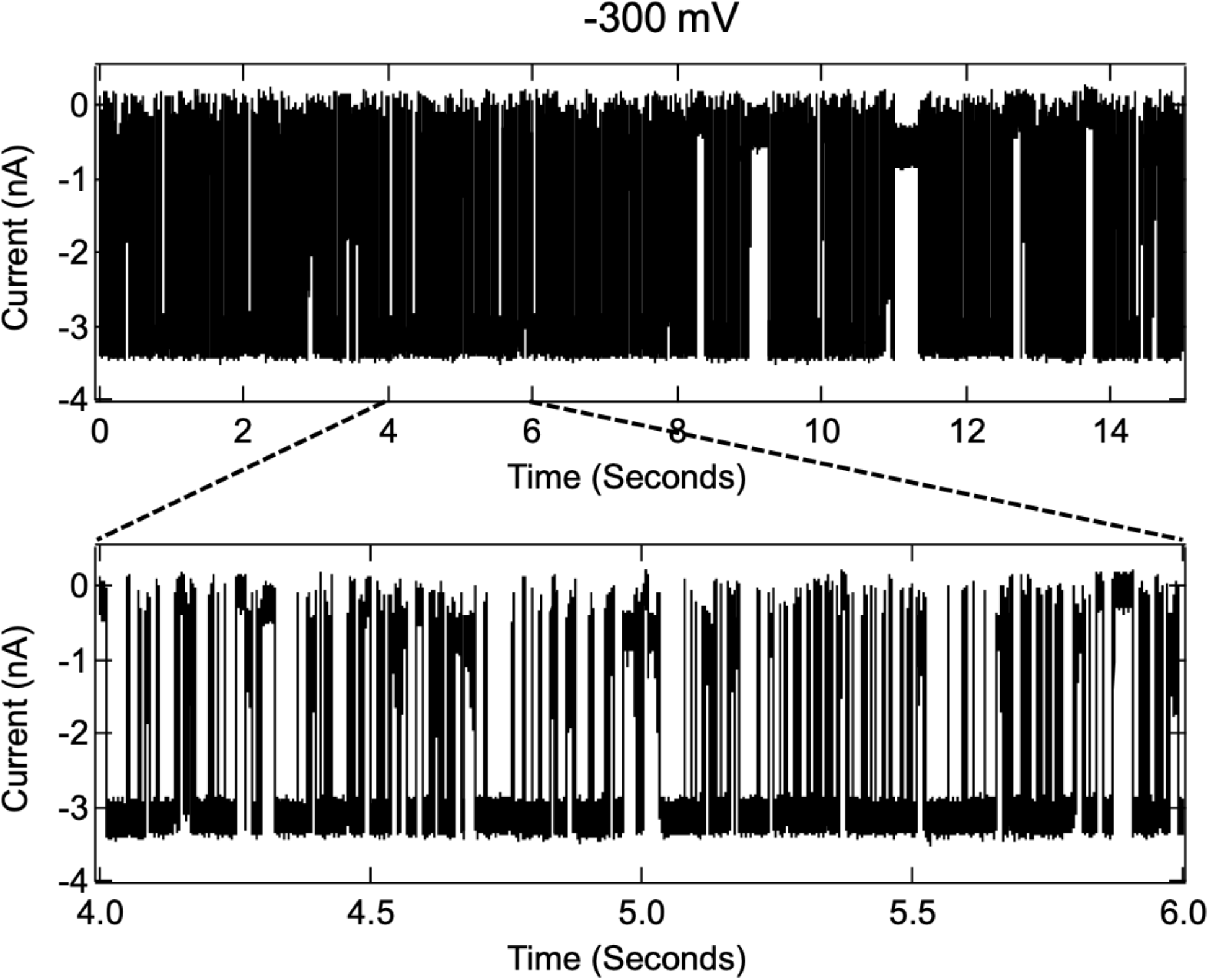
Ionic current traces for cyt c using a 2.5 nm pore at −300 mV applied voltage. The measurements were carried out in 1M, KCl, 10mM HEPES, pH 7.5, sampling rate of 250 kHz, and filtered using a low-pass Bessel filter of 100 kHz.

**Figure S3(iii).**
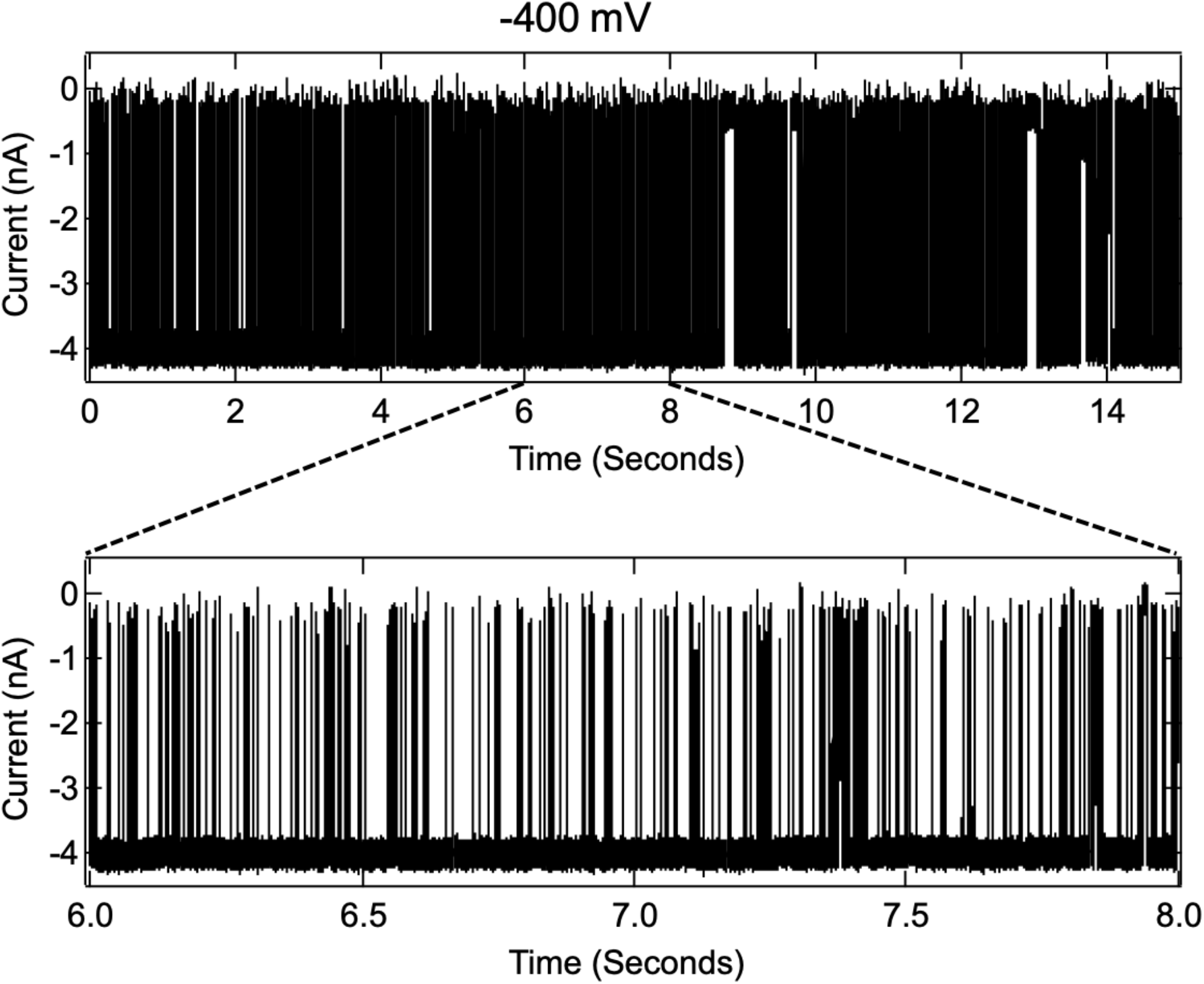
Ionic current traces for cyt c using a 2.5 nm pore at −400 mV applied voltage. The measurements were carried out in 1M, KCl, 10mM HEPES, pH 7.5, sampling rate of 250 kHz, and filtered using a low-pass Bessel filter of 100 kHz.

**Figure S3(iv).**
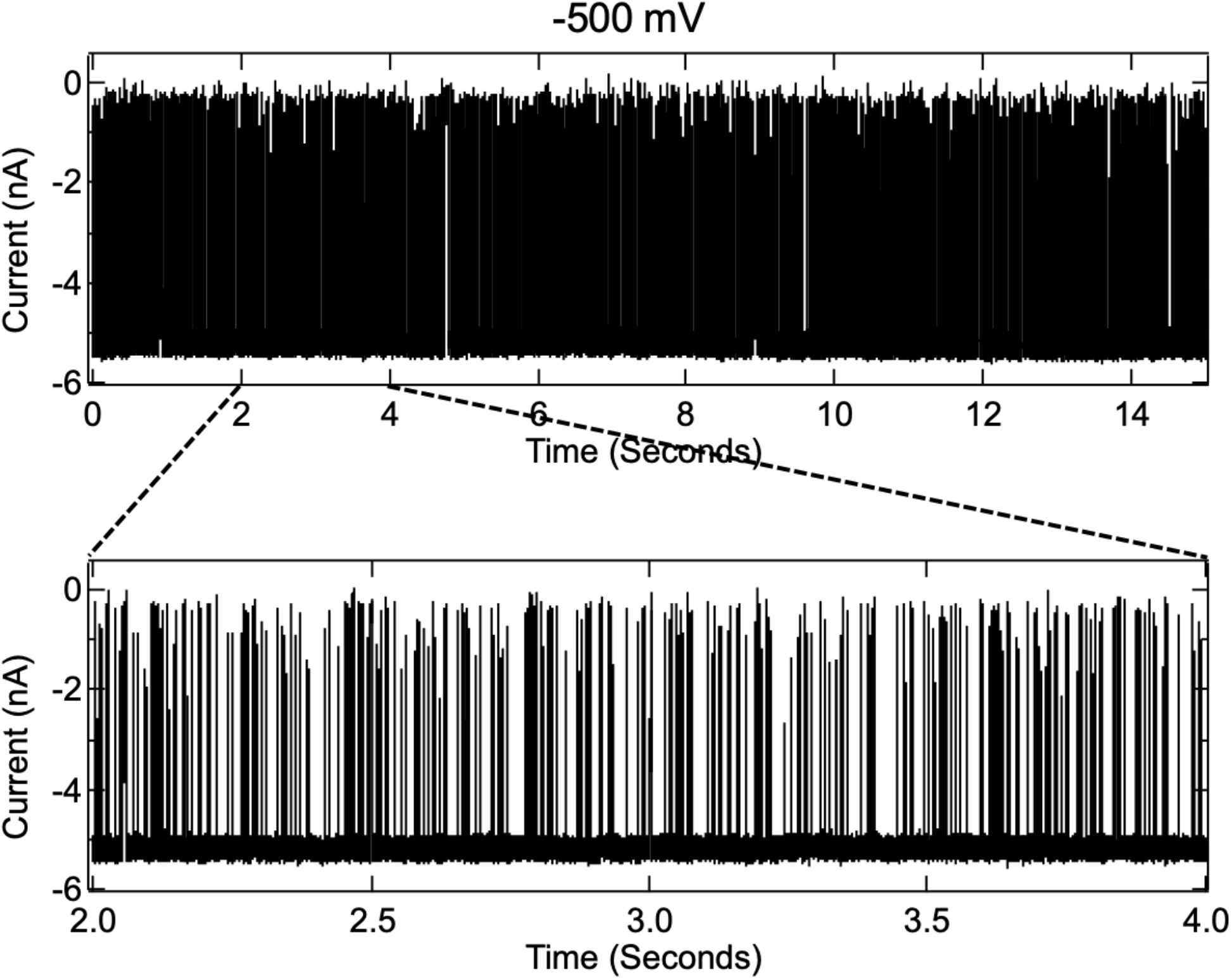
Ionic current traces for cyt c using a 2.5 nm pore at −500 mV applied voltage. The measurements were carried out in 1M, KCl, 10mM HEPES, pH 7.5, sampling rate of 250 kHz, and filtered using a low-pass Bessel filter of 100 kHz.

**Figure S3(v).**
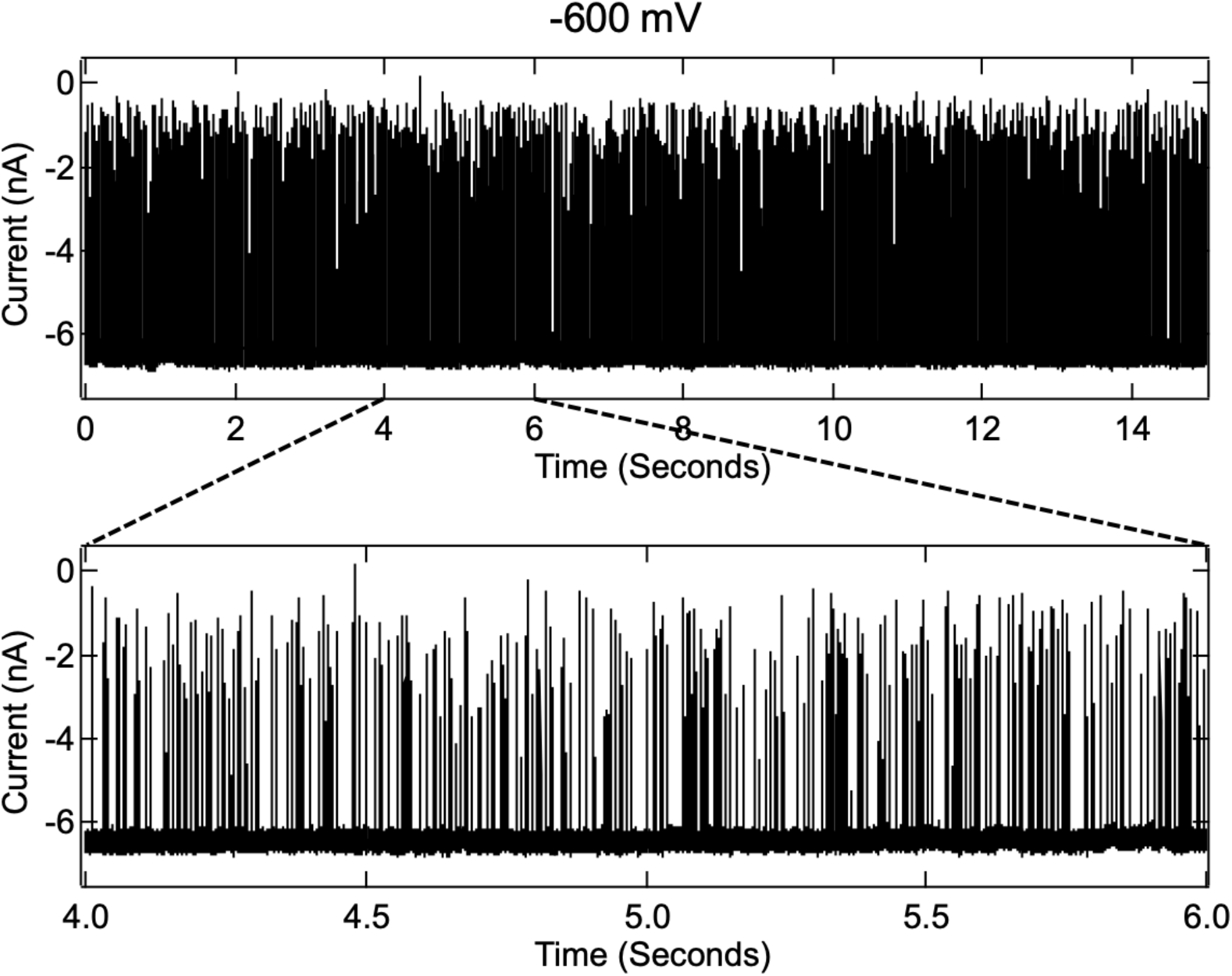
Ionic current traces for cyt c using a 2.5 nm pore at −600 mV applied voltage. The measurements were carried out in 1M, KCl, 10mM HEPES, pH 7.5, sampling rate of 250 kHz, and filtered using a low-pass Bessel filter of 100 kHz.

**Figure S3(vi).**
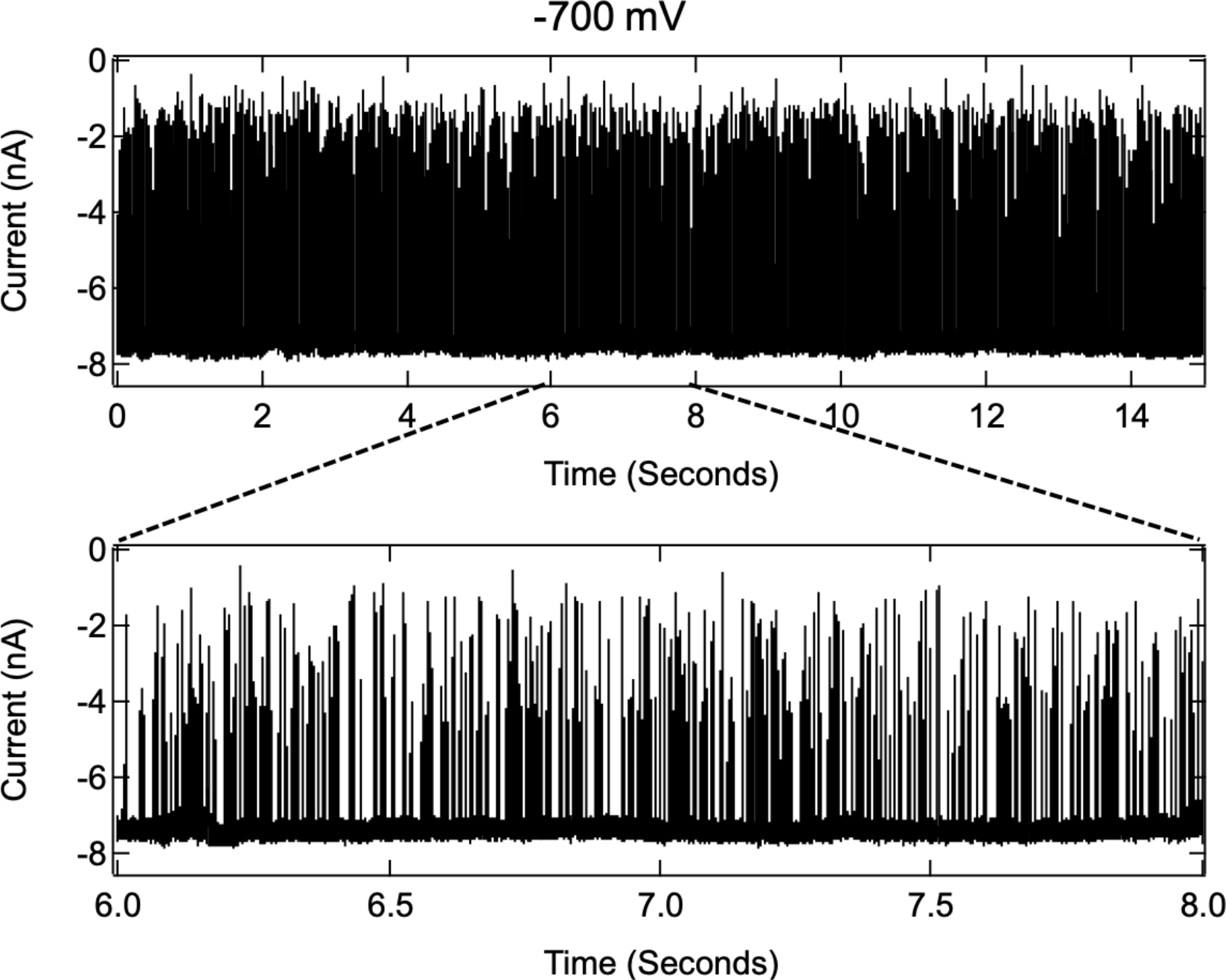
Ionic current traces for cyt c using a 2.5 nm pore at −700 mV applied voltage. The measurements were carried out in 1M, KCl, 10mM HEPES, pH 7.5, sampling rate of 250 kHz, and filtered using a low-pass Bessel filter of 100 kHz.

**Figure S3(vii).**
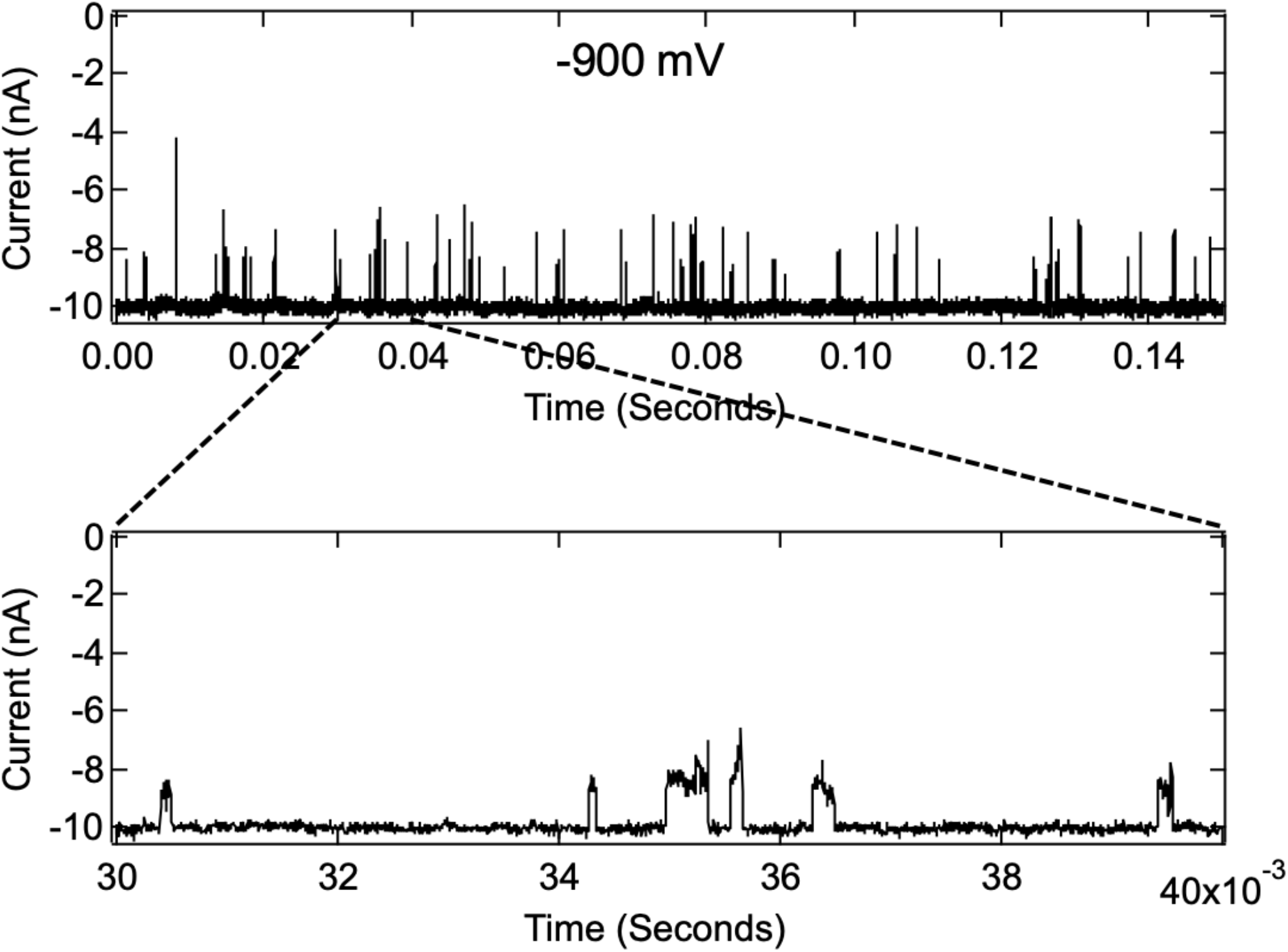
Ionic current traces for cyt c using a 2.5 nm pore at −900 mV applied voltage. The measurements were carried out in 1M, KCl, 10mM HEPES, pH 7.5, sampling rate of 250 kHz, and filtered using a low-pass Bessel filter of 100 kHz.

## 7. Two-state model for ΔI/I_*o*_ vs. E_app_ in the dynamical unfolding limit (for 2nm pore)

An example of the two-state model for a field-dependent fractional blockade is given by Eq. 2 in the text:

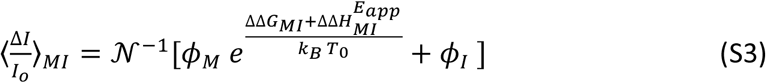

where 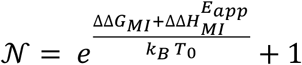. Boltzmann’s constant is denoted by *K*_*B*_ and *T*_0_ is room temperature. This equation was applied to analyze the interconversion of two conformational states on the unfolding pathway (*M* and *I*, where *G*_*M*_ < *G*_*I*_ in zero applied field). A single cyt c molecule is suggested to be trapped and retained by an electric field at the mouth of the 2 nm pore, giving rise to the level *i* blockade changes as the electric field is increased. The fractional blockades for the *I*-state and *M*-state are denoted in Eq. S3 by *ϕ*_*I*_ and *ϕ*_*M*_, respectively. The energies and the difference dipoles are referenced to the native state so that: **ΔΔ** *G*_*MI*_ = **Δ** *G*_*NI*_− **Δ** *G*_*MN*_. For the limit where the permanent dipole term in Eq. 1 of the text is dominant, we have:

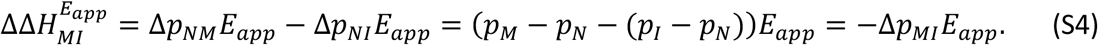

For the limit where the induced dipole term in Eq. 1 of the text is dominant, we have:

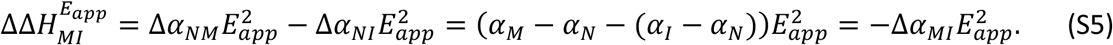

Thus, as the field increases, the more energetic *I*-state is lowered relative to the *M*-state and their respective fractional blockades at the mouth of the 2 nm pore can be found by fitting the level *i* blockade data shown in Fig. 2C of the main text. It should be emphasized that the fractional blockade for a given state is strongly dependent upon both the pore size and whether or not the given conformation is interconverting at the mouth of the pore or is trapped (i.e., squeezed) more deeply inside the pore.

## 8. Representative current traces for cyt c using different pore diameters at −100 mV

**Figure S4.**
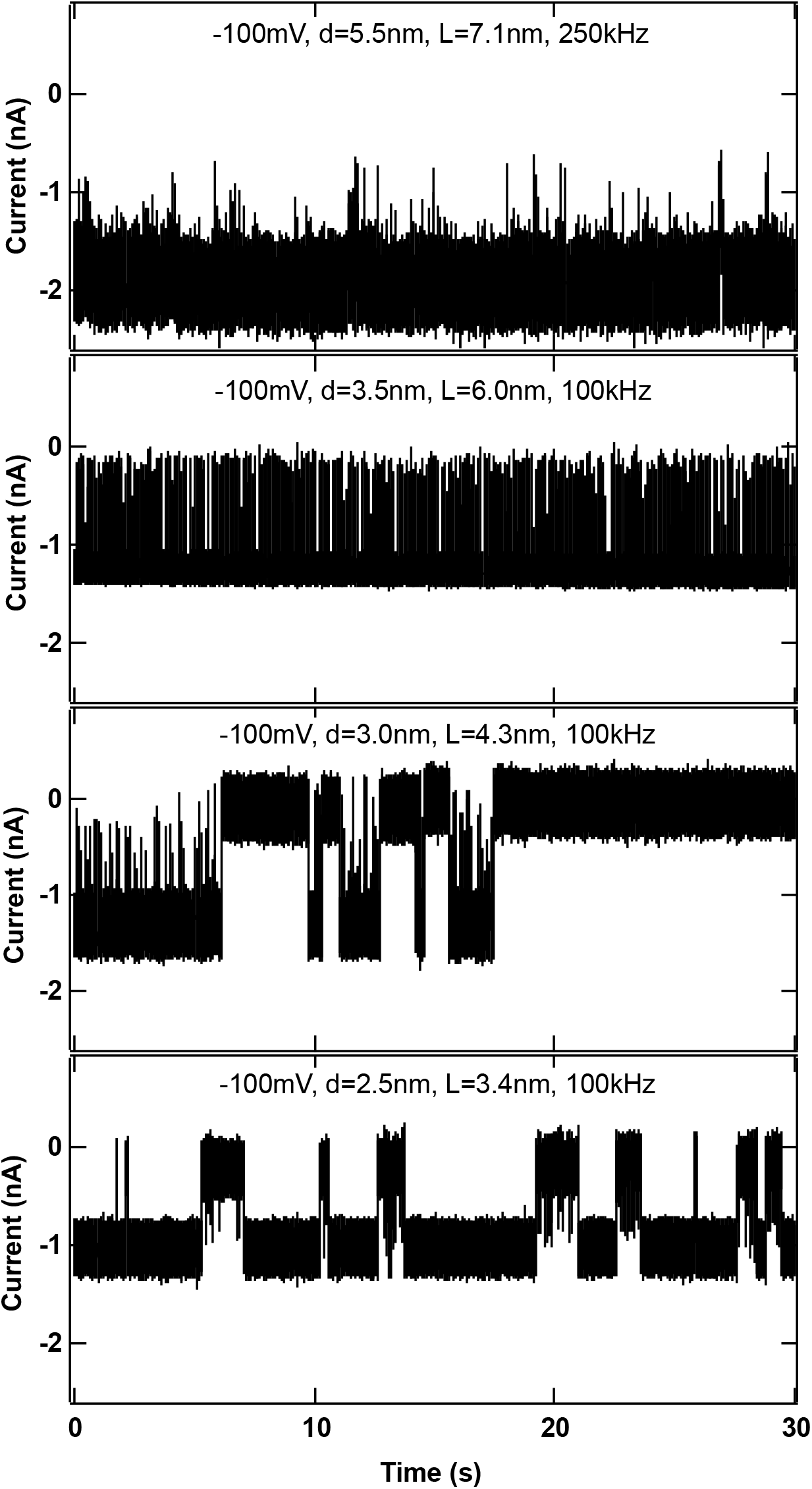
Ionic current traces for cyt c using pores in the diameter range of 2.5 nm < d_pore_ < 5.5 nm at −100 mV applied voltage. The measurements were carried out in 1M, KCl, 10mM HEPES, pH 7.5. The 5.5 nm pore data was measured using Chimera VC100 Instruments, and filtered at low pass Bessel filter of 250 kHz whereas all other data were measured using Axopatch and filtered at 100 kHz.

## 9. Representative current traces for cyt c using 2 nm and 1.5 nm diameter pores

**Figure S5.**
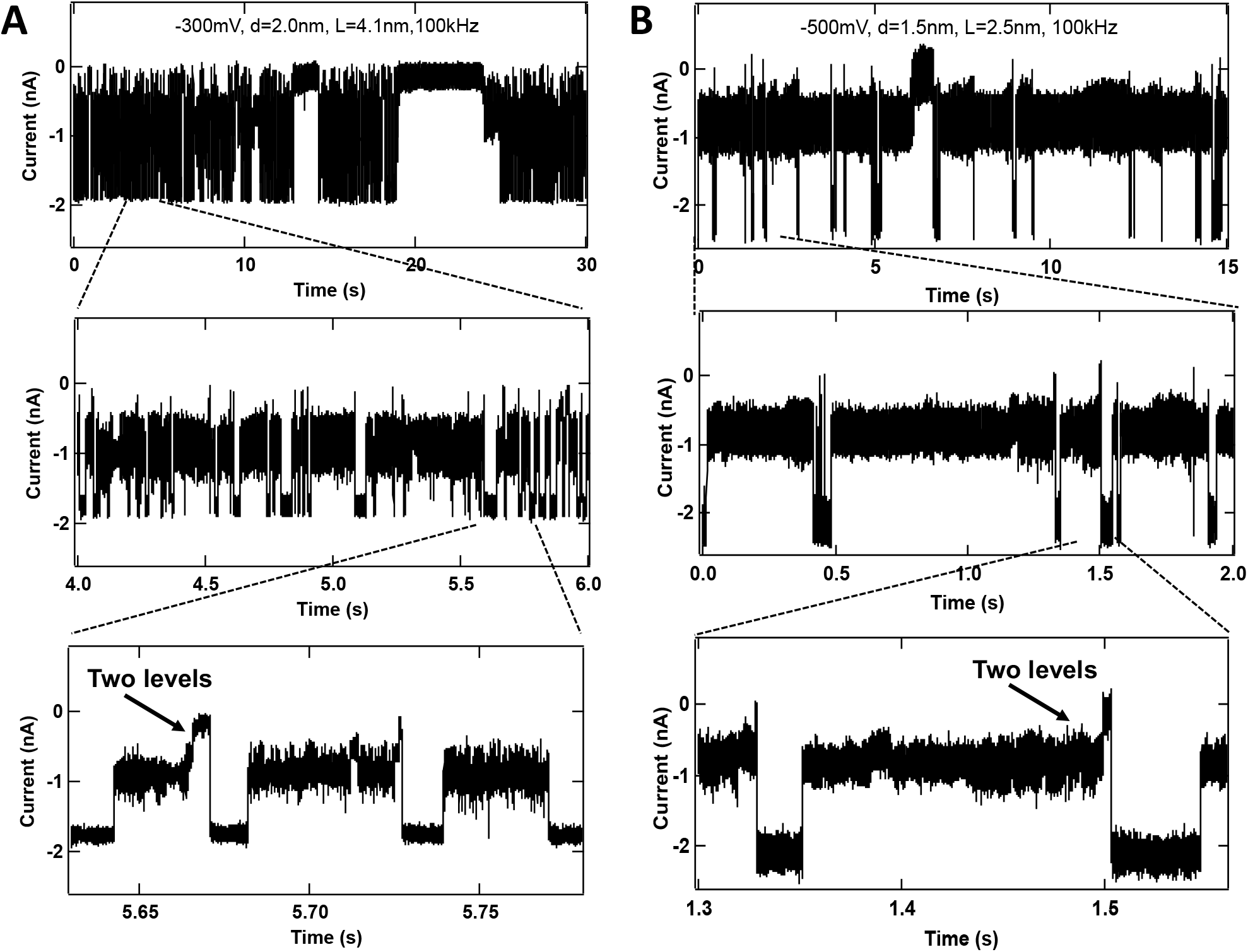
Ionic current traces for cyt c using **(A)** a 2 nm diameter pore at −300 mV, and **(B)** a 1.5 nm diameter pore at −500 mV. The measurements were carried out in 1M, KCl, 10mM HEPES, pH 7.5. The data were filtered at 100 kHz and recorded at a sampling rate of 250 kHz. Two-level events were rarely observed below a threshold voltage strength −250 mV for the 2nm pore and –500 mV for the 1.5 nm pore.

## 10. Representative ΔI/I_*o*_ histograms and fraction of events with level *ii* for cyt c using a 2 nm pore

**Figure S6.**
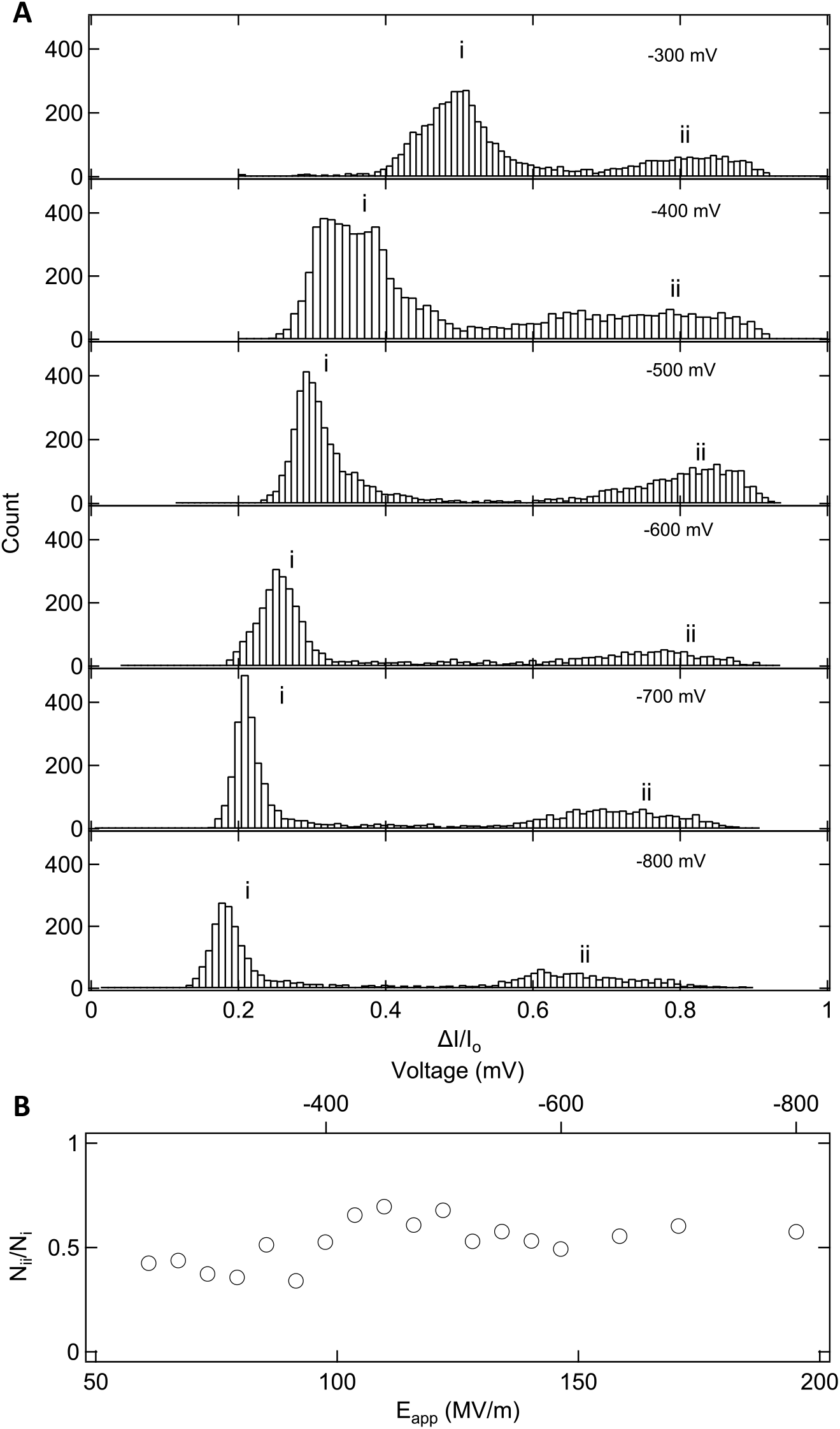
**(A)** Representative histograms of **Δ***I***/***I*_*o*_ measured for a 2nm pore (L = 4.1 nm) at various voltages and **(B)** fraction of level *ii* events. The histograms show two clear populations (level *i* and level *ii*) with the peak position of level *i* decreasing with voltage.

## 11. MD simulation methods and data for 1.5 nm pore

### i: Molecular models and general systems preparation

The hexagonal patches of Si_3_N_4_ membranes were generated using the Inorganic Builder plugin of VMD (9). The membranes were aligned with the x-y plane of the coordinate system. By the removal of atoms from the membrane, a double-cone pore was created in each membrane. The axis of the cone was aligned with the z-axis. The minimum diameter of the cone was set to be at the middle of the membrane (which henceforth is call pore diameter); the cone angle was 15° degrees (with respect to the z axis). The charge of the atoms comprising the Si_3_N_4_ membrane was adjusted by a small (< 0.1%) amount to make the Si_3_N_4_ membrane neutral. Table S2 summarizes the geometry of the simulated systems. Initial atomic coordinates of cytochrome c were taken from the crystal structure reported by Bushnell et al (PDB code: 1HRC) (10). Missing hydrogen atoms were added to the protein using the PSFGen plugin of VMD. The minimum distance between the protein and membrane was set to be at least 15Å away from the membrane surface to ensure that, at the beginning of the simulation, there were no contact forces between the membrane and the protein. Next, a pre-equilibrated volume of TIP3 water was added to the system using the Solvate Plugin of VMD (11). Following that, potassium and chloride ions were added using the Autoionize VMD plugin to produce 1 M KCl solution.

**Table S2:** Geometry of the simulated systems

| Pore diameter<br>(nm) | Membrane<br>thickness<br>(nm) | Simulation box size*<br>(nm × nm) | Number of atoms | |
| --- | --- | --- | --- | --- |
|  |  |  | Open pore | Protein + pore |
| 1.5 | 2.5 | $3.8 \times 11.4$ | 53619 | 53148 |
| 2.0 | 4.1 | $3.8 \times 13.2$ | 61669 | 61129 |
| 2.5 | 3.4 | $3.8 \times 12.3$ | 57664 | 57124 |
| 3.0 | 4.3 | $3.8 \times 13.4$ | 63061 | 62484 |
| 3.5 | 6.0 | $4.2 \times 14.9$ | 85037 | 84457 |
| 5.5 | 7.1 | $5.7 \times 16.1$ | 169241 | 168741 |

\*The box size is specified as the side edge length of the hexagonal prism and the height of the simulation box.

### ii: Molecular dynamics simulation

All of the simulations were performed using the molecular dynamics program NAMD2 (12). To describe the atomic interactions in the simulations, CHARMM36 (13) force field parameters for the protein and ions, TIP3P model for water and the custom force field describing crystalline Si_3_N_4_ (14) were used. For van der Waals and short-range electrostatic interactions, a smooth cutoff of 12 Å with a switching function starting at 10 Å was used. Long range electrostatic interactions were evaluated with particle mesh Ewald (PME) (15) over a 1 Å-spaced grid. Periodic boundary conditions were employed in all simulations. Each system was minimized using the conjugate gradient method for 5000 steps followed by an NPT equilibration run of 10 ns at 295 K and 1 atm. The constant pressure was realized using a Nose-Hoover Langevin piston (16) and temperature was maintained at a constant value by coupling the system to a Langevin thermostat (17).

### iii: Simulation Protocols

To simulate electric field-driven transport of cyt c through the Si_3_N_4_ nanopores, grid-steered molecular dynamics (G-SMD) method (18) was employed. In this method, nanopore transport of biomolecules is accelerated by subjecting the solute’s atoms to a grid-based potential that accurately reproduces the distribution of the electrostatic potential in the nanopore system. The accelerated transport of the solute is obtained when the force from such a grid-based potential is amplified along the direction of the nanopore transport and selectively applied to the atoms of the solute. In our case, the distribution of the electrostatic potential was determined by simulating each nanopore system for 10 ns in pure 1M KCl solution (without the cyt c protein) under a 1V transmembrane voltage. The voltage *V* was generated by applying a constant electric field *E* = -V/*L*_Z_ along the z axis, where *L*_Z_ is the length of the simulated system along the direction of the applied electric field (19). As the resulting distribution of the electrostatic potential was highly non-homogeneous (especially within the pore) and depended on the pore geometry, the open pore simulations were performed for each pore geometry. The average distributions of the electrostatic potential was then calculated and averaged over the respective MD trajectory using the PMEpot Plugin of VMD (19), producing a 1 Å-spaced grid potential. In the G-SMD simulations of the nanopore transport, the potential was applied to all atoms of the cyt c protein with the scaling factor of 1, 2 or 3 for the z component of the electric field and 0 for the x and y components. The force applies to each atom of the cyt c molecule by the extrernal grid potential was scaled by the partial charge of that atom. In addition to G-SMD, a custom colvar script was used to harmonically restrain the protein’s center of mass to the nanopore axis; the spring constant was 10 kcal/(mol Å^2^).

### iv. Analysis of the simulation results

VMD was used to analyze and post process the simulation trajectories. The steric exclusion model (SEM) (20) of nanopore conductance was used to measure the fractional blockade current during the protein permeation. As aforementioned in the previous section, the membrane (as well as the simulation box) was a hexagonal prism, however, the SEM is developed for a cubic simulation box. Therefore, SEM was only applied to the largest inscribed rectangular in the hexagon of the membrane (See Fig. S7). To quantify the deformation/unfolding state of the protein during the permeation, the fraction of native contacts, Q-value (21), was calculated for the structured parts (sheets and helices) of the protein in each simulation. For calculating the Q-value, the reference structure of the protein was the initial crystal structure of the protein.

**Figure S7.**
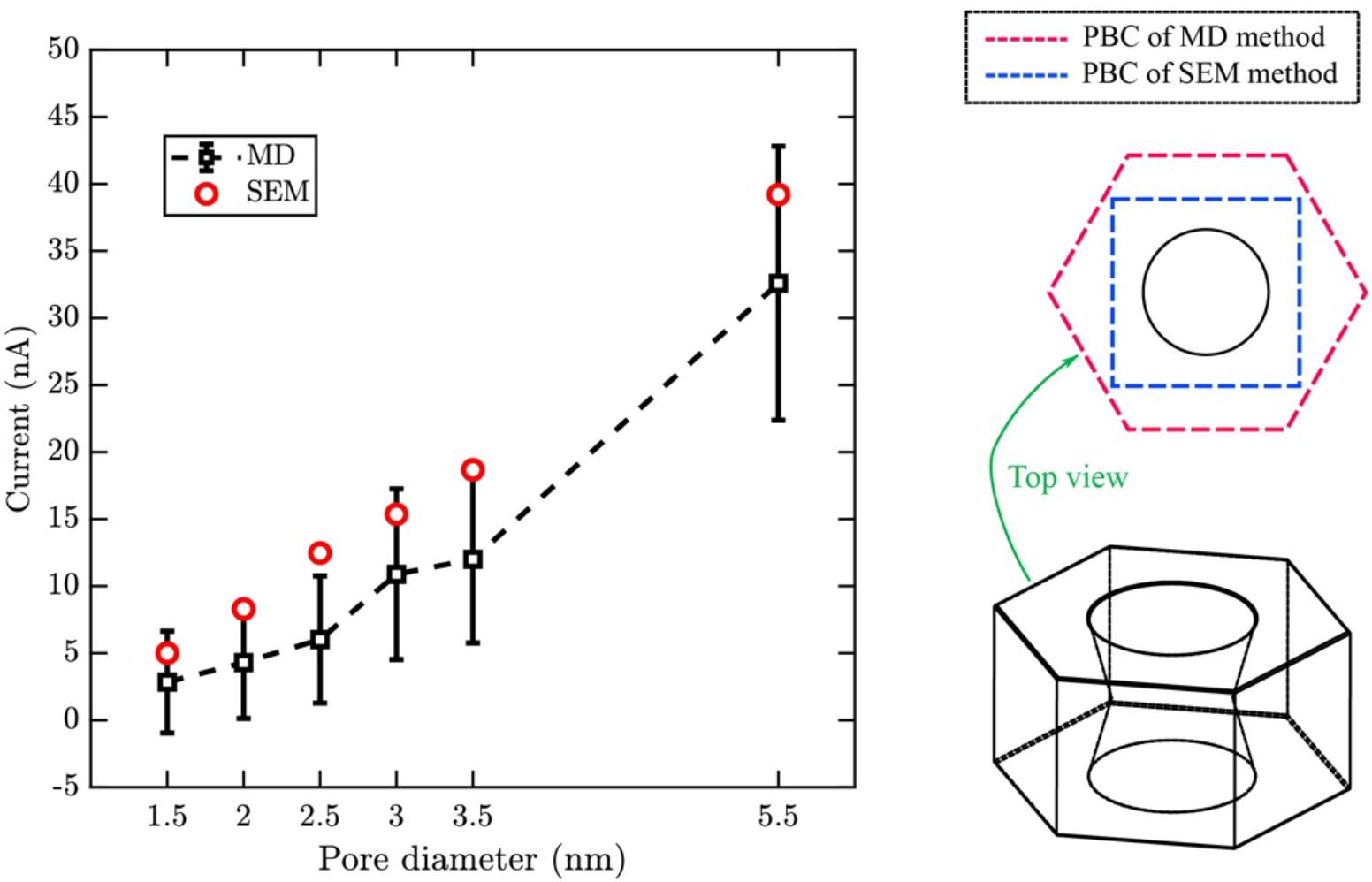
Open-pore currents obtained directly from all-atom simulations (black) and from SEM calculations (red). Each MD current value was obtained by averaging instantaneous ion displacements over a 10 ns MD trajectory at 1V bias. The error bars show the standard deviations of the 5 ps-sampled current traces; the dashed line is the guide to the eye. The bottom right image shows a schematic 3D representation of the simulated Si_3_N_4_ membrane whereas the top right image shows how the system’s cross section is represented in SEM.

**Figure S8.**
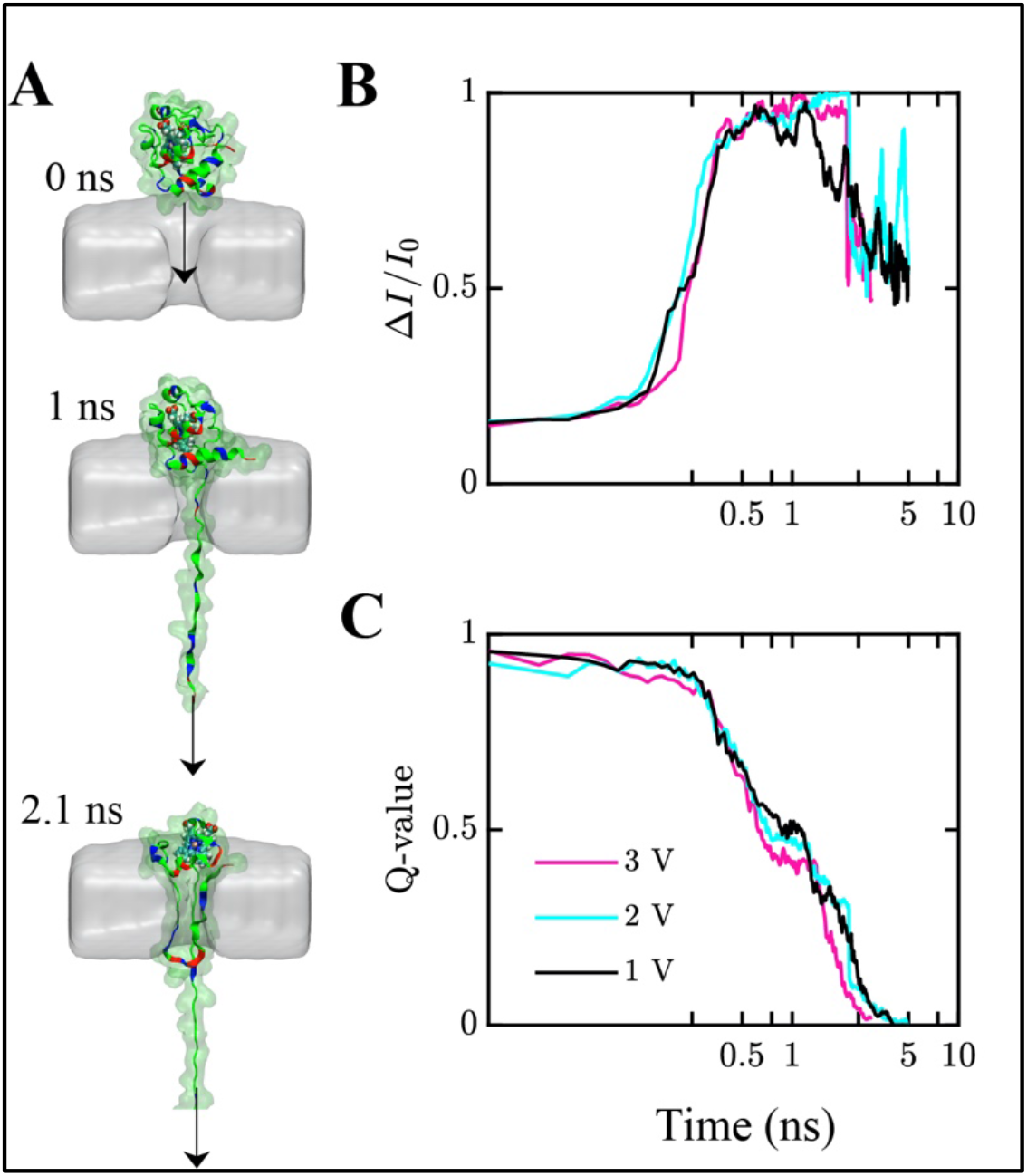
MD simulation of cyt c translocation through a 1.5 nm pore. **(A)** Snapshots representing an MD trajectory where a single cyt c protein was forced to pass through a 1.5 nm nanopore (gray) using the G-SMD protocol under a 3V effective bias and the constant velocity SMD pulling (illustrated by the black arrow). **(B, C)** Ionic current blockade (B) and cyt c Q-value (C) versus simulation time for three simulations carried out at the specified effective biases. In addition to G-SMD, the N-terminus of the protein was pulled along the pore axis with 0.1 Å/ps velocity using the SMD protocol, the SMD spring constat was 5.5 kcal/(mol Å^2^). The SEM approach was used to calculate the ionic current blockades. Note the logarithmic scale of the horizontal axis.

## 12. Distributions of ΔI/I_*O*_ as a function of voltage for cyt c using a 3 nm pore

**Figure S9.**
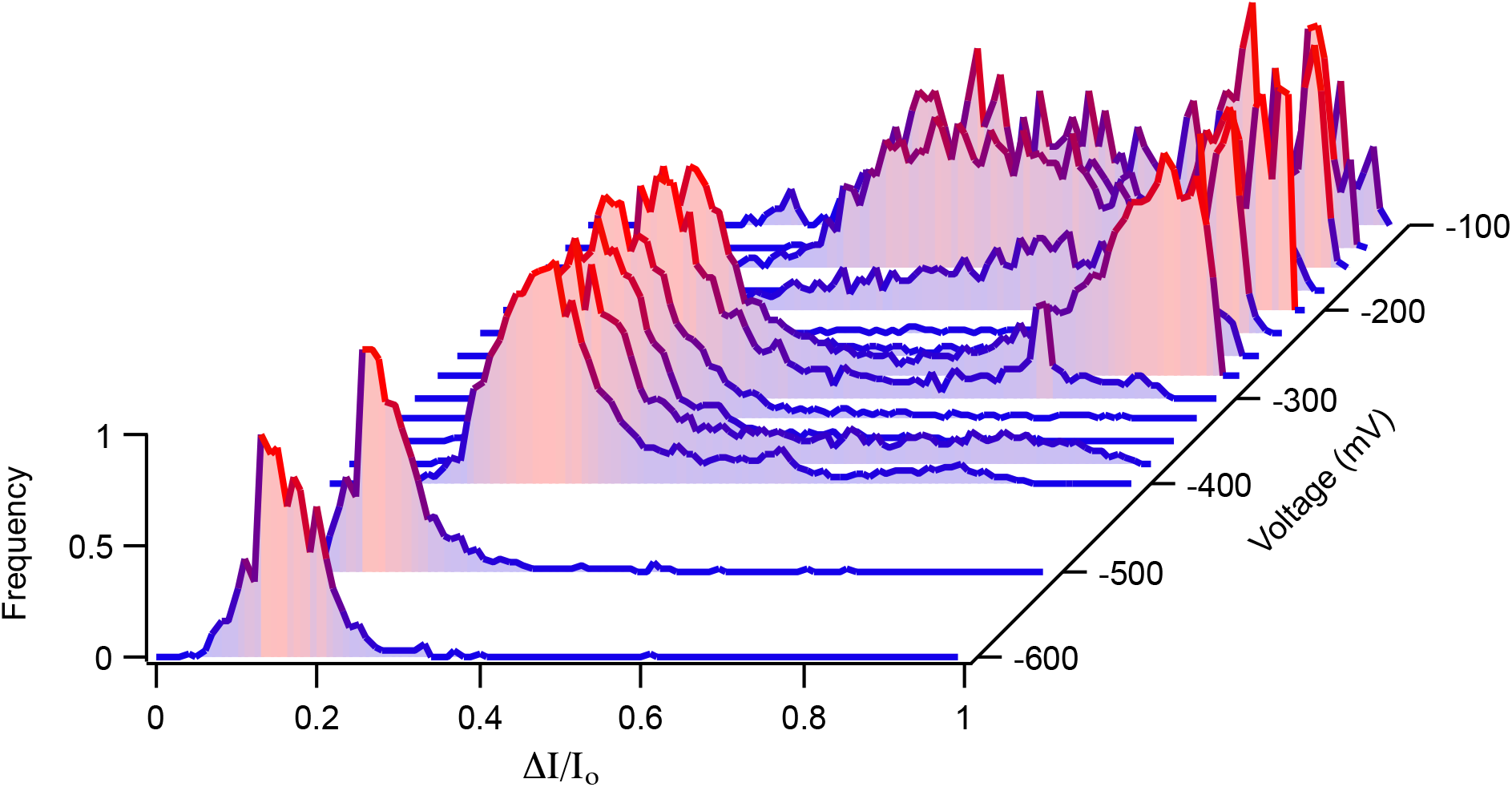
Distributions of fractional change in current as a function of applied voltage measured for cyt c using a 3 nm diameter pore, L =4.3 nm (1M KCl, 10mM HEPES, pH7.5).

## 13. Experiments in presence of Gdm-Cl (Denaturation data)

**Table S3:** Conductivity of bulk solution and conductance of nanopore (d_pore_= 2.5 nm). The conductance was measured as described in section 3.

| Electrolyte solution<br>(10 mM HEPES, pH 7.5) | Conductivity in<br>bulk solution<br>(mS/cm) | Conductance in<br>Nanopore (nS) |
| --- | --- | --- |
| 0.5M Gdm-Cl, 1M KCl | 130 mS/cm | 11 nS |
| 1M Gdm-Cl, 1M KCl | 158 mS/cm | 13 nS |
| 2M Gdm-Cl, 1M KCl | 207 mS/cm | 14.5 nS |
| 3M Gdm-Cl, 1M KCl | 240 mS/cm | 16.6 nS |

**Figure S10.**
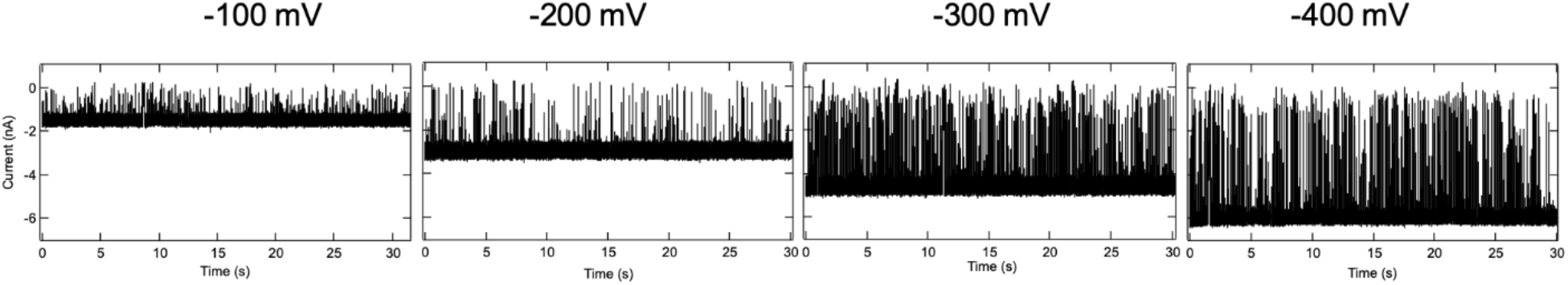
Ionic current traces for cyt c (0.5 μM) translocation in 2M Gdm-Cl, 1M KCl, 10mM HEPES, pH7.5 solution at different voltages for a 2.5 nm pore (L=3.4 nm).

**Figure S11.**
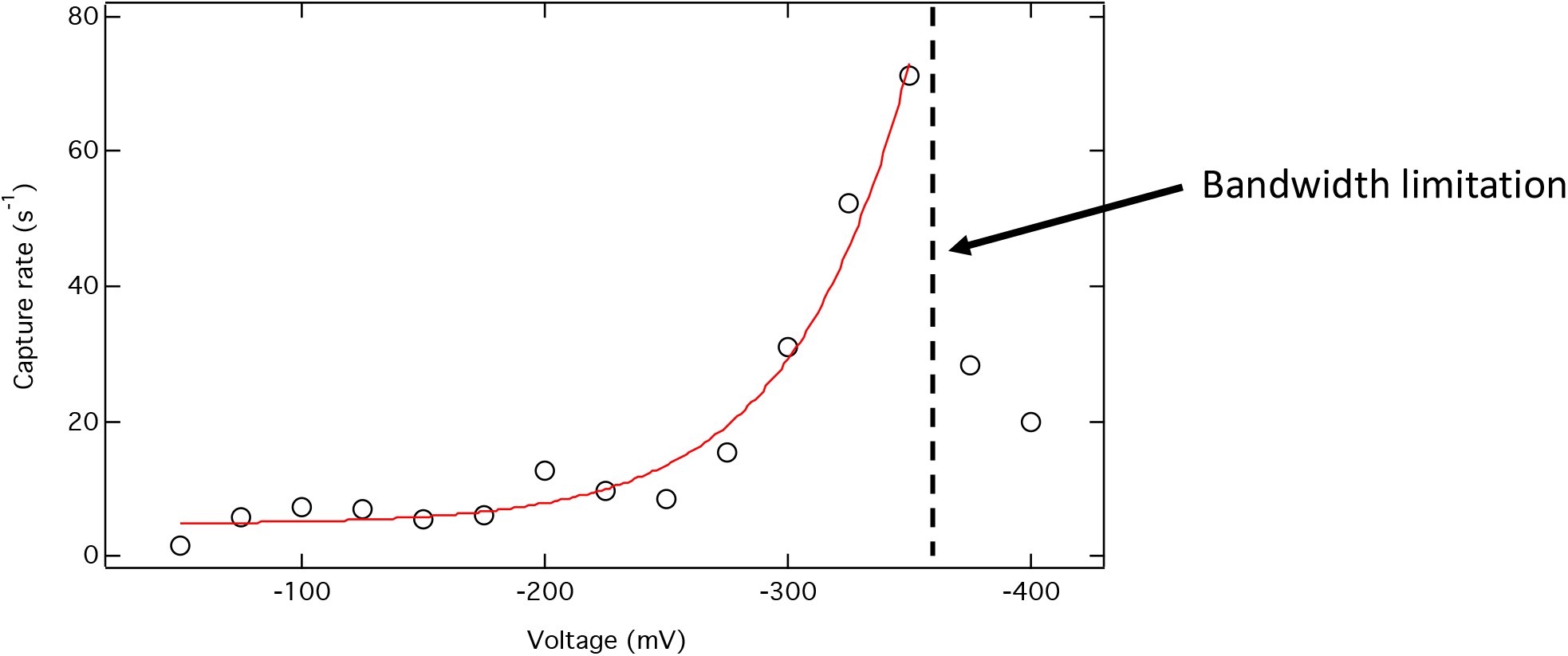
Capture rate as a function voltage for cyt c (0.5 µm) using a 2.5 nm pore under 2M Gdm-Cl, 1M KCl, 10mM HEPES, pH 7.5. The red curve represents a fit to the function 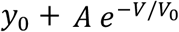, with *y*_0_ = 4.70, *A* = 0.05, and *V*_0_ = 48.6. The observed exponential dependence reflects an entropy barrier for capture, consistent with a blob-like polymer behavior of an unfolded protein. Above −350 mV capture rates were observed to drastically decrease, due to extremely fast translocation pulses for unfolded cyt c, which our limited bandwidth of measurement cannot resolve.

**Figure S12.**
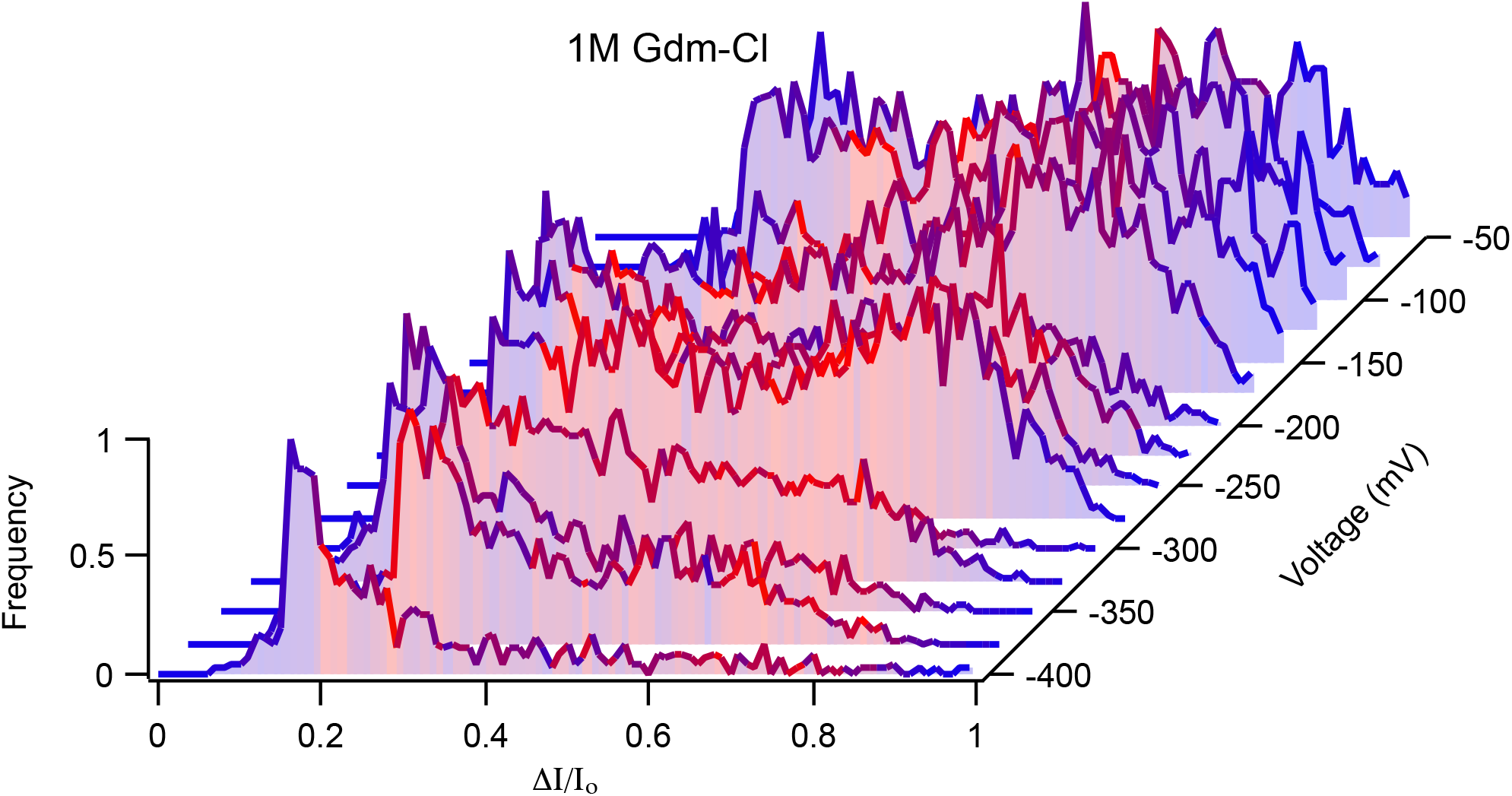
Distributions of fractional blockade as a function of applied voltage measured for cyt c using a 2.5 nm pore. Cyt c (0.5 μM) was incubated in 1 Gdm-Cl, 1M KCl, 10mM HEPES, pH7.5 solution.

**Figure S13.**
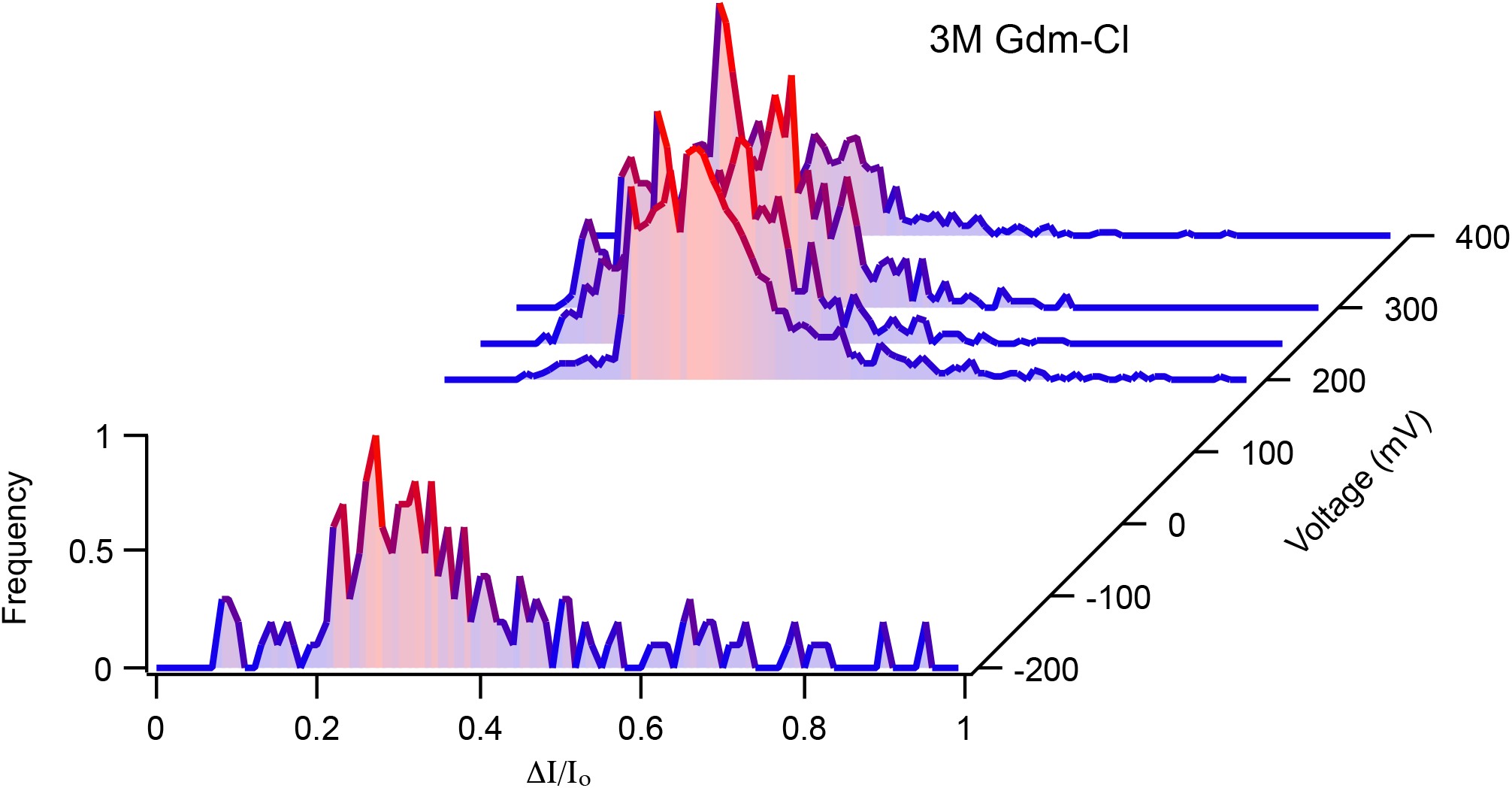
Distributions of fractional change in current as a function of applied voltage measured for a 2.5 nm pore, when 0.5 μM cyt c were incubated in 3 Gdm-Cl, 1M KCl, 10 mM HEPES, pH7.5 solution.

**Figure S14.**
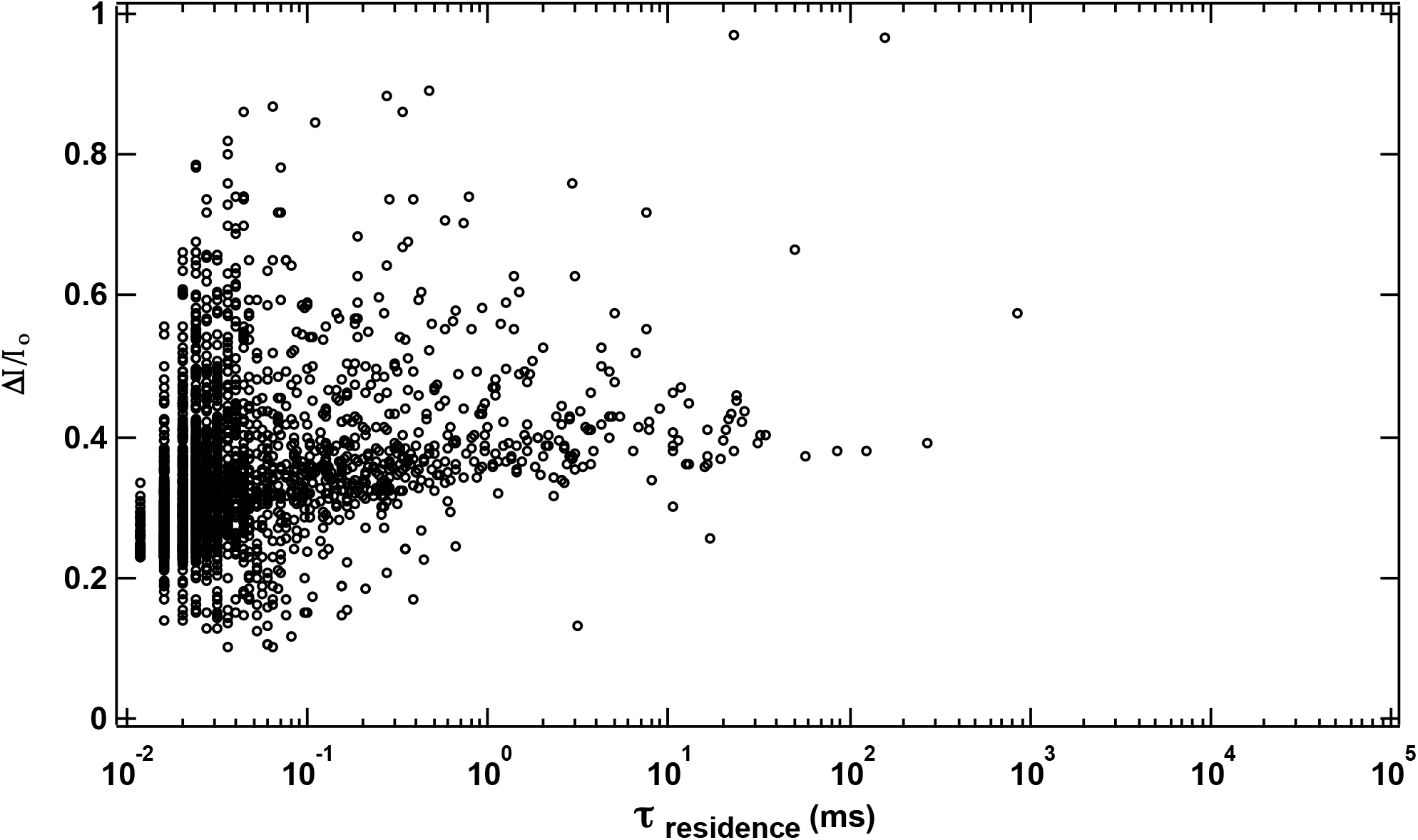
Scatter plots of fractional current blockade and residence time measured at 200 mV and in 3M Gdm-Cl, 1M KCl, 10mM HEPES, pH7.5 solution (d_pore_ = 2.5 nm, L = 3.4 nm). The concentration of cyt c used was 0.5 μM.

**Figure S15.**
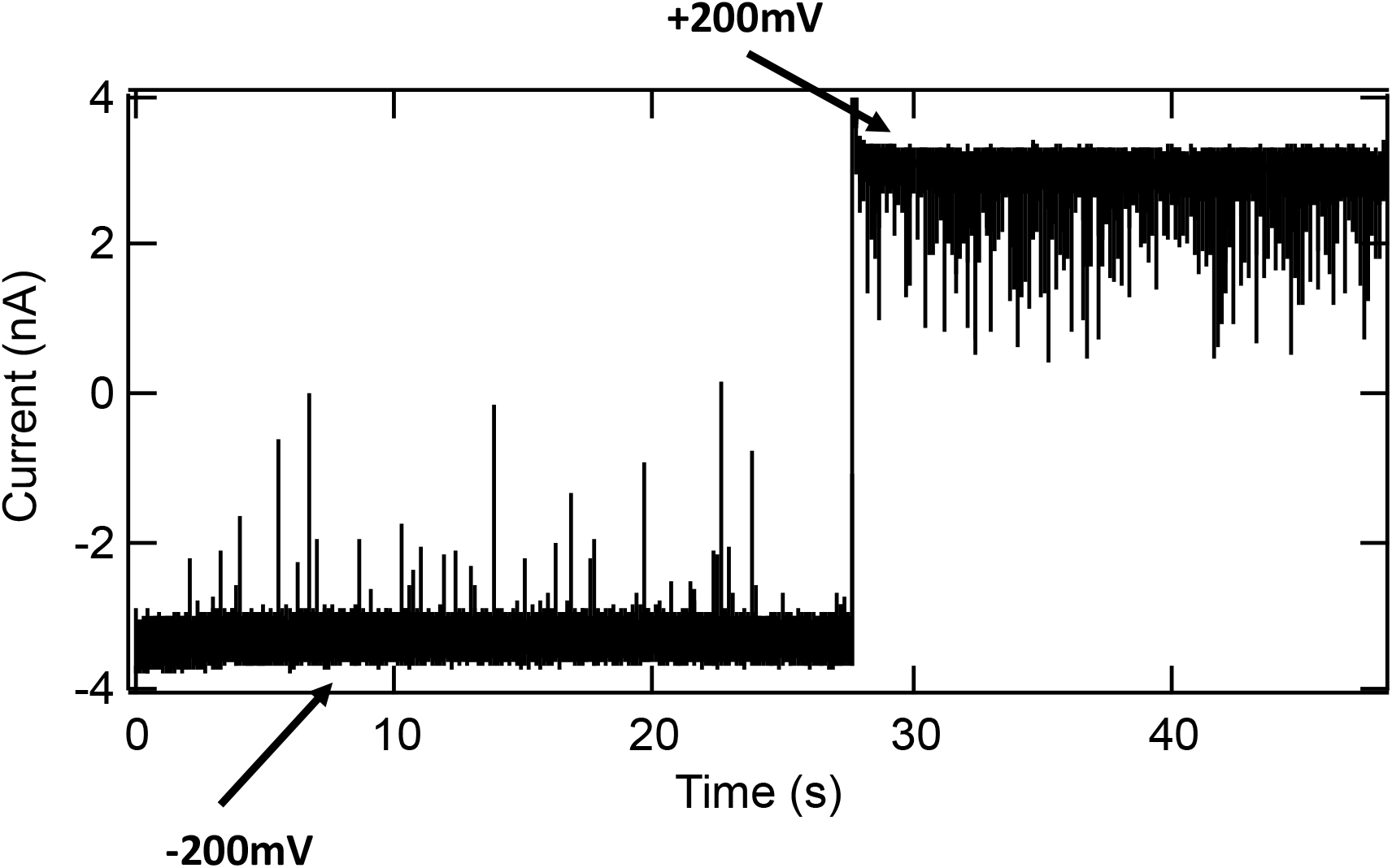
Ionic current trace for cyt c (0.5 μM) in 3M Gdm-Cl, 1M KCl, 10mM HEPES, pH7.5 (d_pore_ = 2.5 nm, L = 3.4 nm). We observed translocation events more frequently at +200 mV compared to −200 mV. We did not observe events at positive voltage in other experiments. While we do not understand why this occurs, it could be due to other effects such as charge-reversal of the pore at this high Gdm-Cl concentration, which can influence the mechanism of protein capture at the pore. A similar effect has been observed for poly-ethylene glycol (PEG), where 4 M KCl was used to drive the translocation of the otherwise neutral polymer (22).

## 14. Multicomponent Gaussian fits to the Δ1/1_0_ distributions of cyt c using a 2.5 nm pore

**Figure S16.**
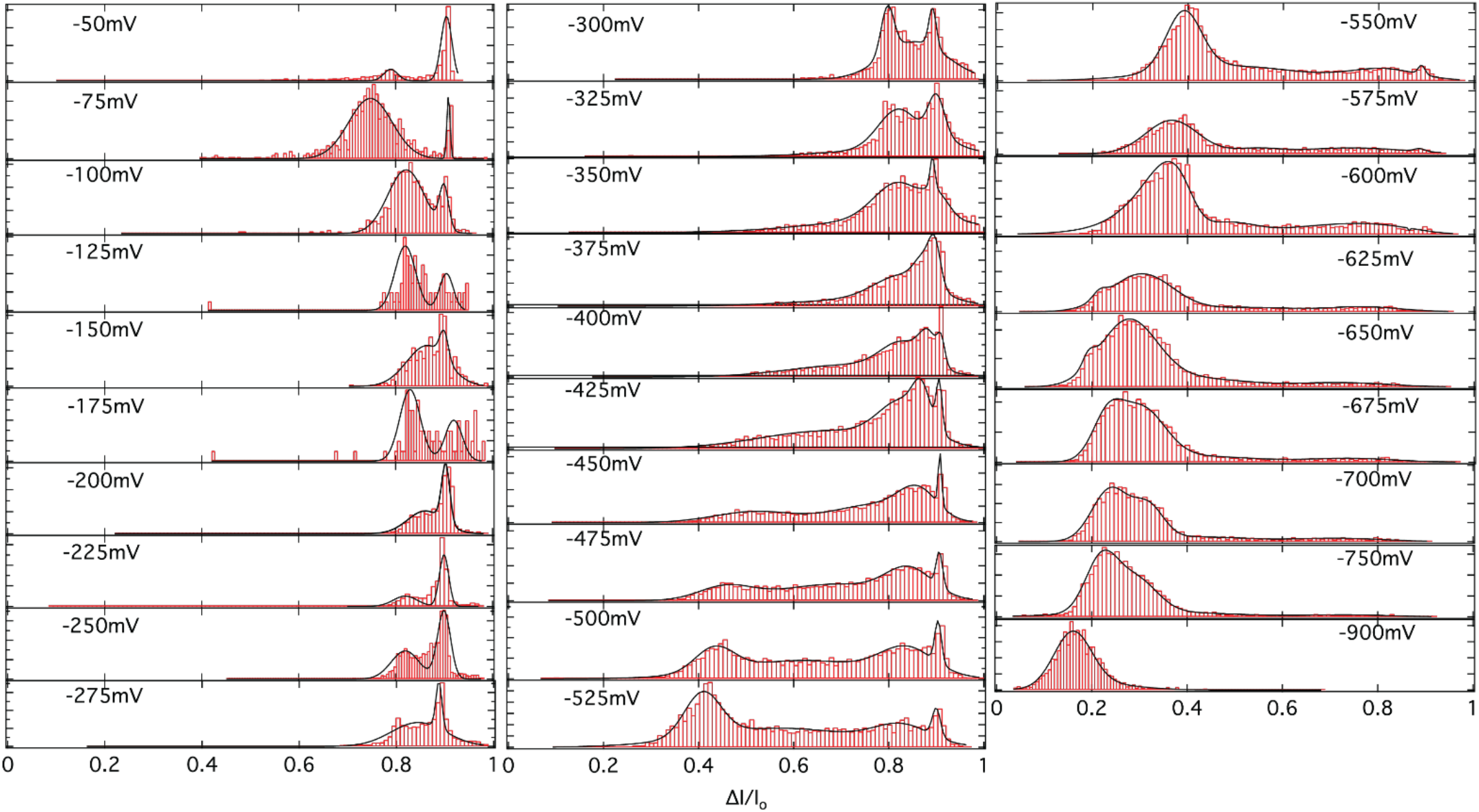
Distributions of Δ*I*/*I*_*0*_ and their fits to a multi-component Gaussian function at different voltages for the pore size d_pore_ =2.5nm, L =3.4nm (1M KCl, 10mM HEPES, pH7.5)

## 15. Two-state and three-state model for ΔI/I_*0*_ vs. Eapp in the dynamical unfolding limit (analysis for 2.5 nm pore)

We consider the case where state & squeezes into the 2.5 nm pore under the action of an applied electric field and, as the field is increased, dynamic unfolding and refolding of the three cyt c *α*-helices can take place. This can be visualized using either a two-state model between *M* and *U* or within a three-state scheme where an intermediate *I*-state, with one of the cyt c *α*-helices unfolded, is populated. The blockade data for the 2.5 nm pore are shown in Fig. 3E of the text. The sub-states *M*_1_ and *M*_2_ are thought to involve two different, non-interconverting, squeezed pore configurations of cyt c where the stabilizing salt bridge, involving residue E62, has broken and there is a loss of the associated short 2 stranded beta sheet (23). This is followed by Ω-loop unfolding of residues 40-57 and 60-87 (including Met80 dissociation). The two *M*-states are differently squeezed and configured within the pore and do not appear to interconvert with each other so the configuration notation (1 or 2) can be attached in an *ad hoc* manner.

The 3-state equilibrium average is given by:

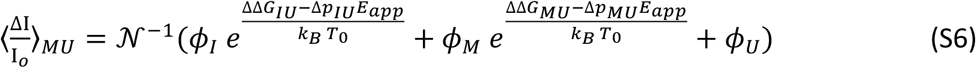

with 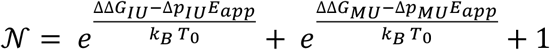 Where, for simplicity, we have used only the simple permanent dipole difference term as a parameter to describe the action of the electric field on the conformational state free energy. The energy gap parameters can be simplified by using the 2 nm pore results for *M-I* equilibration so that: ΔΔ*G*_*MU*_ = ΔΔ*G*_*MI*_ + ΔΔ*G*_*IU*_= *4*6*K*_*B*_*T*_0_*+*ΔΔ*G*_*IU*_,where the value of ΔΔ*G*_*MI*_ in Table 1 of the text has been used. Similarly, for a simple oriented dipole with an effective net charge separation (*d*_*i*_) in each state given by *d*_*U*_ > *d*_*l*_ > *d*_*M*_, we can deduce that: Δ*p*_*MU*_ =Δ*p*_*MI* +_Δ*p*_*IU*_=44*Debye*_+_Δ*p*_*IU*_, again using the information from Table 1. This results in a 3-state fitting function for the dynamic transitions in the 2.5 nm pore that can be written as:

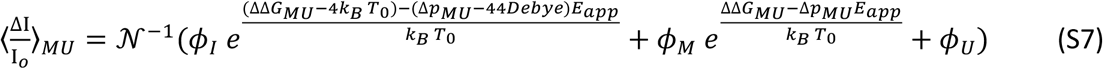

with 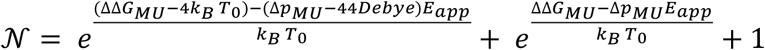 Equation S7 has one additional free parameter compared to a two-state model.Because the much broader distribution of blockade current ratios for the *M*_1_ ↔ *U*_1_ transition suggests more direct access to the unfolded conformations, we used a 2-state model to fit its blockade ratio in the 2.5 nm pore. In this case, we assume a direct interconversion between *M*_1_ and a broad set of unfolded states*U*_1_, rather than sequentially passing through the *I*-state. The two-state model used for the dynamic unfolding was:

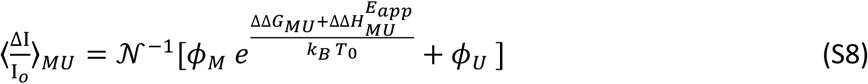

where 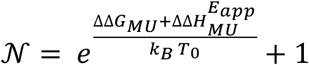.

We were able to successfully fit the 2.5 nm pore blockage ratio data for both *M*_1_ and *M*_2_using the two-state model (Eq. S8) as seen in Table S4. Nearly indistinguishable fits were found whether we take 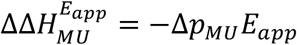 or 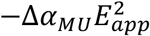. The value of the polarizability difference, Δ*α*_*MU*_, is presented in Table S4 as an induced dipole(Δ*α*_*MU*_*E*_*mid*_), where we used the midpoint fields in Fig. 3E, *E*_*mid*_= 140.3 MV/m and 158.9 MV/m for *M*_2_ ↔ *U*_2_ and *M*_1_ ↔ *U*_1_ respectively.

**Table S4:** Two-state fitting (4 free parameters) for the experimental data (d_pore_ = 2.5 nm, L = 3.4nm) in Fig. 3E using Eq. S8 with either Δ*p*_*MU*_ or Δ*α*_*MU*_ set to zero.

| Parameters | $M_1 \leftrightarrow U_1$ | $M_2 \leftrightarrow U_2$ |
| --- | --- | --- |
| $\phi_M$ | 0.91 | 0.82 |
| $\phi_U$ | 0.24 | 0.20 |
| $\Delta\Delta G_{MU}$ | $8.9 k_B T_0$ (5.3 kcal/mol) | $8.2 k_B T_0$ (4.9 kcal/mol) |
| $\Delta p_{MU}$ | 70.2 Debye | 71 Debye |
| $\Delta\alpha_{MU} E_{mid}$ | 73 Debye | 81.5 Debye |

We also fit the data with the 3-state model, Eq. S7, which assumes that intermediate *I* between *M* and *U* is also accessed. In order for the fits to converge, we found that the blockade ratio for the *I*-state must be similar to that of the &-state, consistent with unfolding of only one of the three *α*-helices. In this scenario for the 2.5 nm pore, both *M* and *I* lead to high blockade ratios because they still contain significant *α*-helical content that evidently leads to ion current blockage. The fitting results for the dynamics of the 3-state system are given in Table S5, where we have constrained *ϕ* _*I*_ by using *ϕ* _*I*_ = *ϕ*_*M*_ so that only 4 free parameters are needed for the fit.

**Table S5:** 3-state fitting (4 free parameters) for the experimental data (d_pore_ = 2.5 nm, L = 3.4nm) in Fig. 3E using Eq. S7.

| Parameters | $M_1 \leftrightarrow I \leftrightarrow U_1$ | $M_2 \leftrightarrow I \leftrightarrow U_2$ |
| --- | --- | --- |
| $\phi_M$ | 0.91 | 0.86 |
| $\phi_I$ | 0.91 | 0.86 |
| $\phi_U$ | 0.23 | 0.18 |
| $\Delta\Delta G_{MI}$ | $4.0 k_B T_0$ (2.4 kcal/mol) | $4.0 k_B T_0$ (2.4 kcal/mol) |
| $\Delta\Delta G_{MU}$ | $11.2 k_B T_0$ (6.6 kcal/mol) | $9.1 k_B T_0$ (5.4 kcal/mol) |
| $\Delta p_{MI}$ | 44 Debye | 44 Debye |
| $\Delta p_{MU}$ | 101.5 Debye | 91.4 Debye |

## 16. Representative histograms of *τ*_*residence*_ for a 2.5 nm pore (L = 3.4 nm) at higher electric fields

**Figure S17.**
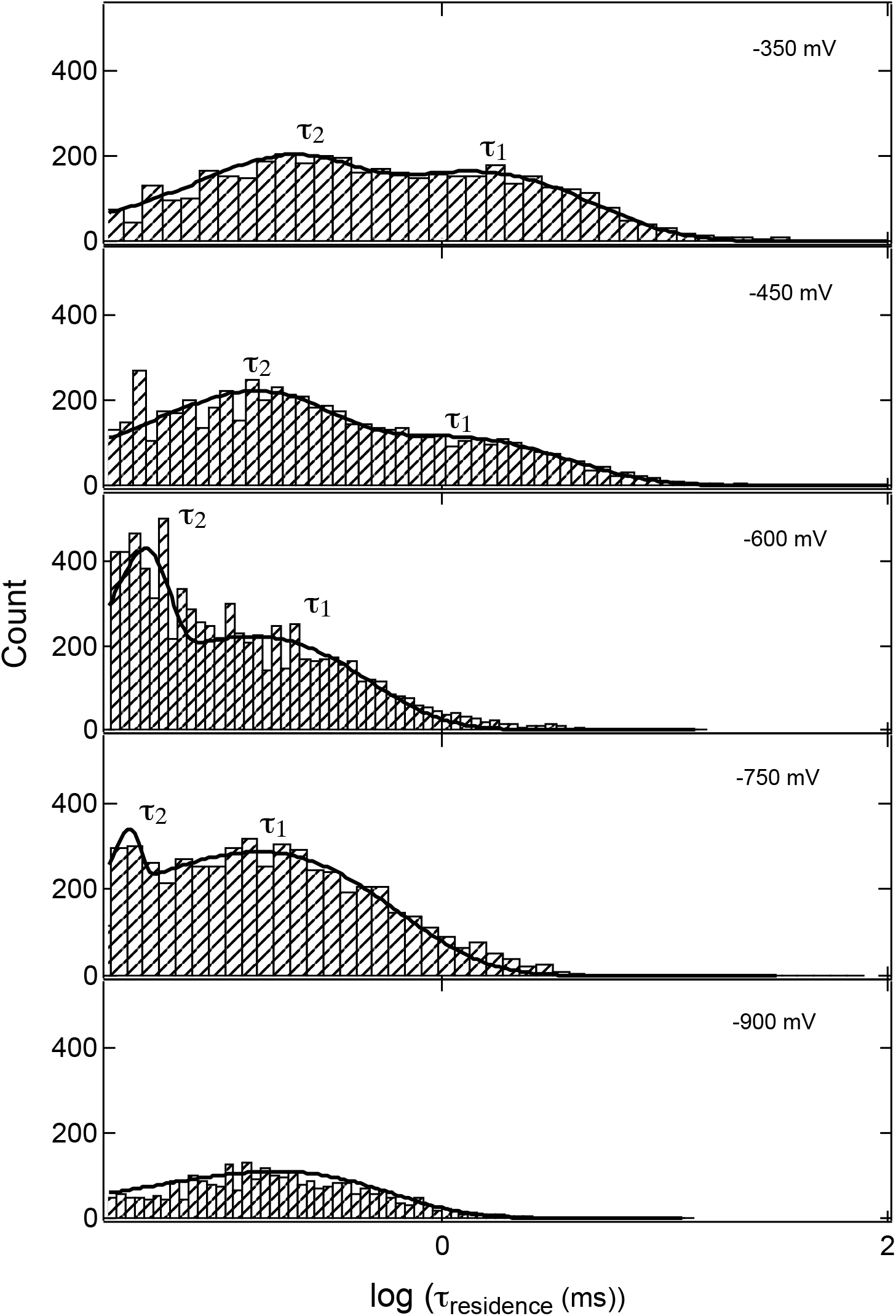
Typical distributions of *τ*_*residence*_ measured at −350 mV, −450 mV (*M ↔ U* regime) and at −600 mV, −750 mV, −900 mV (*U* regime). The solid curve represents its fit with a bi-modal distribution, yielding two-time constants (*τ*_1_ and *τ*_2_) and rates (*k*_1_ and *k*_2_). At −900 mV only a single time constant was observed and attributed to complete linearization of cyt c.

## 17. Translocation kinetics and fast exchange of folding/unfolding *α*-helices (2.5 nm pore)

Here we identify the states *M, I*, and *U* with the folding of the three main *α*-helices of cyt c. We consider them to be in rapid exchange as the electric field reduces the free energy gap between the folded and unfolded states of these short (∼3 turn) *α*-helices that form an important part of the secondary structure of cyt c. As these states undergo thermodynamic exchange on the sub-microsecond timescale(24-26), the main channel for the translocation within the 2.5 nm pore involves the unfolded *U*-state. We hypothesize that the folded *α*-helices block ion current as well as act to retard passage of the protein through the pore, so that the *U*-state is the primary form of the protein during its passage. Once formed, the various configurations and conformations of the *U*-state within the pore are pulled by electrophoretic forces through the pore on a much slower timescales than the folding/unfolding transitions of the α-helical segments. These slower translocation timescales, which are associated with the various *U*-state configurations within the pore, give rise to the distribution of residence times observed on the ms timescale in Fig. S17 and Fig. 3F (insert). In order to enter the pore, a partially unfolded *M*-state must be formed at the mouth of the pore where the E62 salt bridge has broken and the omega loops have loosened or unfolded, allowing squeezing of the protein into the pore in two separate configurations, which we have labeled as *M*_1_ and *M*_2_. We have considered two different routes between the *M*- and *U*-states. One involves a two-state model (a direct transition from *M* where all 3 helices unfold to form *U*) and the other is a three-state model which is sequential(23, 27, 28) and involves formation of an intermediate *I*-state where one of the α-helices unfolds at lower energies, followed by the other two at higher energy, in a stepwise pathway to the *U*-state. In order to better visualize the time-scale separation and the role of the electric field in modifying the state energies, the simpler kinetic scheme for the 2-state model is depicted below. In this example, there are two conformational states (*M*_1_ and *U*_2_) considered in the limit where only the *U*_1_-state is able to effectively translocate through the pore (i.e., where 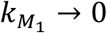).

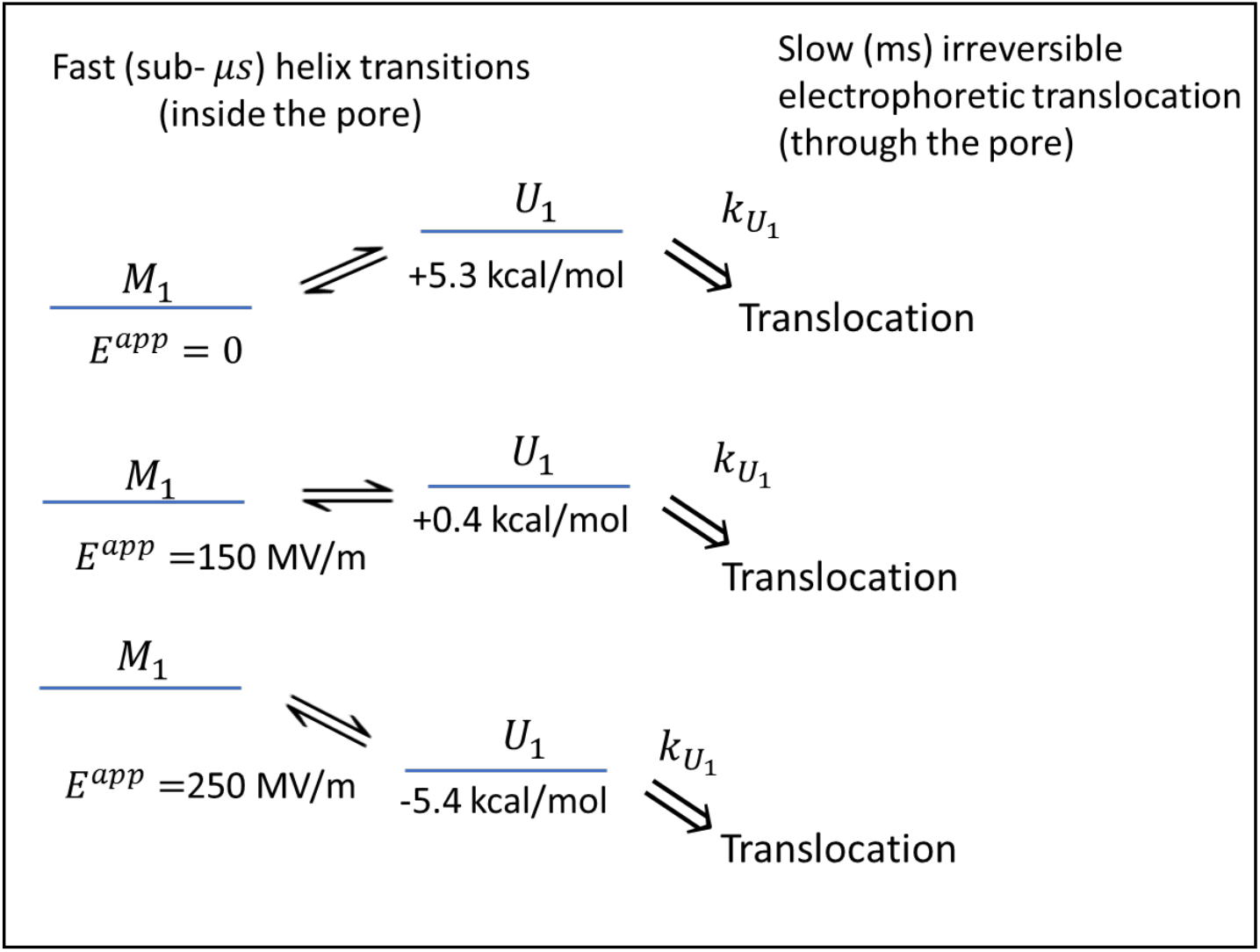

### Two-state Model

When two-states undergo fast interconversion (*M*_1_ *⇌ U*_1_) the overall kinetic translocation rate can be written as:

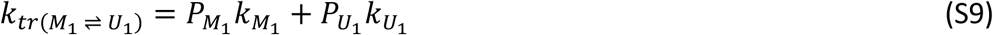

where we take 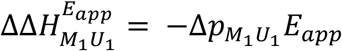, for simplicity and write the probability of finding cyt c in state *U*_1_ as

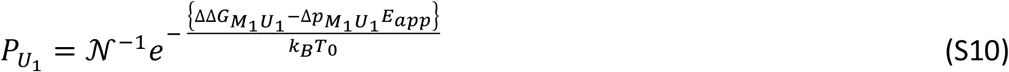

with 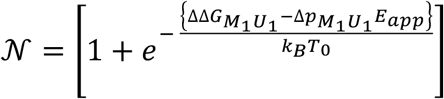

We let 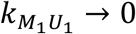 because of its significant α-helical content and then take:

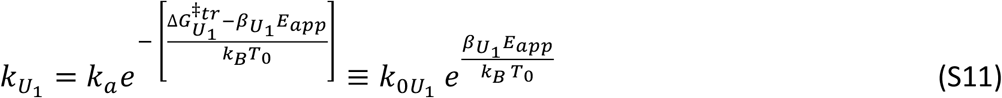

where 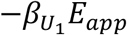 represents the reduction in the translocation kinetic barrier due to the electrophoretic force (29) that pulls the unfolded protein through the pore. Thus, the translocation rate can be written as:

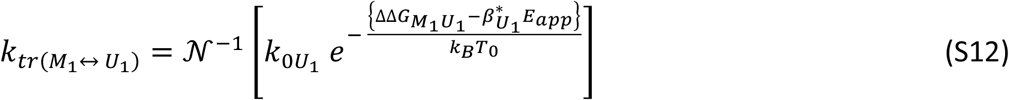

with 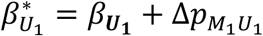, and

### Three-state model

For three-state fast exchange translocation kinetics 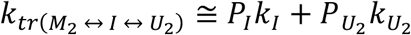, so that:

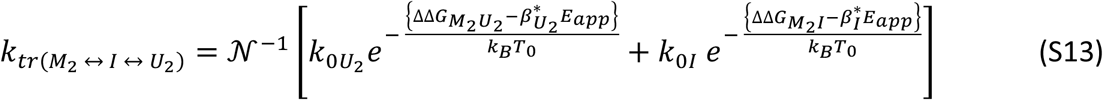

with 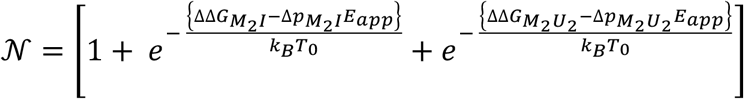 and 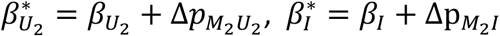.

Where, just as for the two-state model, we have defined:

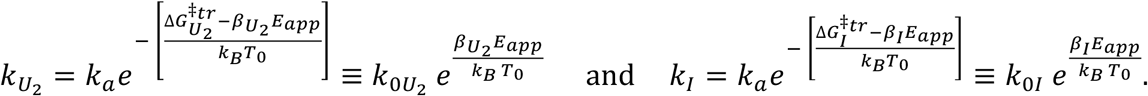

### Fitting parameters

Using the value of 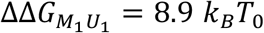 and 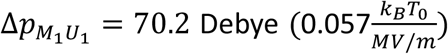 (Table 2 main text), we have 4 unknown parameters 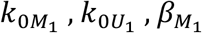 and 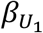 to fit the two-state translocation kinetics. Using the extrapolated translocation rates in Fig. 3F of the main text (Sec. 19, Fig. S20, Table S7), we have 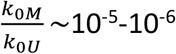. Thus, as noted above, we take 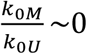 which leads directly to Eq. S12.

In Fig. S18, we fit the *M*_1_ data with Eq. S12 (two unknown parameters 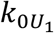 and 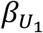), which yields 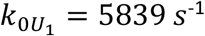 and 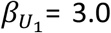 Debye. The value of ln 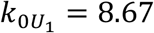 also agrees very well with extrapolated translocation rate of state *U*_1_ in Fig. 3F (see Sec. 19, Fig. S20, Table S7). In Fig. S18, we fit the *M*_1_ data with Eq. S13 and use 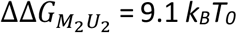 and 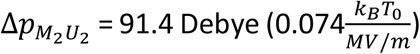 as well as 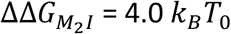 and 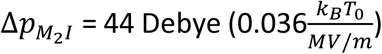from Table 2 of main text. Thus, we have 4 unknown parameters 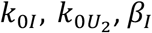 and 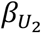. The extrapolated translocation rates from Fig. 3F (see Sec. 19, Fig. S20, Table S7) of the main text again justify setting 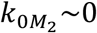. We fit the black circle data using Eq. S13 which yields 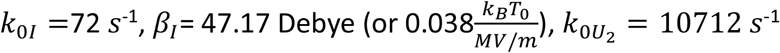, and 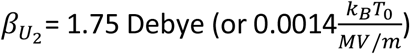. The values of 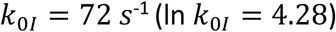 is in very close agreement with the translocation rates (extrapolated to zero field) using data in the *M*_2_ ↔ *I* ↔ *U*_2_ region of Fig. 3F. Further the value of β_1_, which is a measure of the field dependence of the electrophoretic force, is in very close agreement with the slope of translocation rates in *M U* regime. The fit did not converge when we set *K*_*I*_=0 and tried to fit the data with only 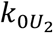 and 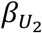as adjustable parameters.

**Figure S18.**
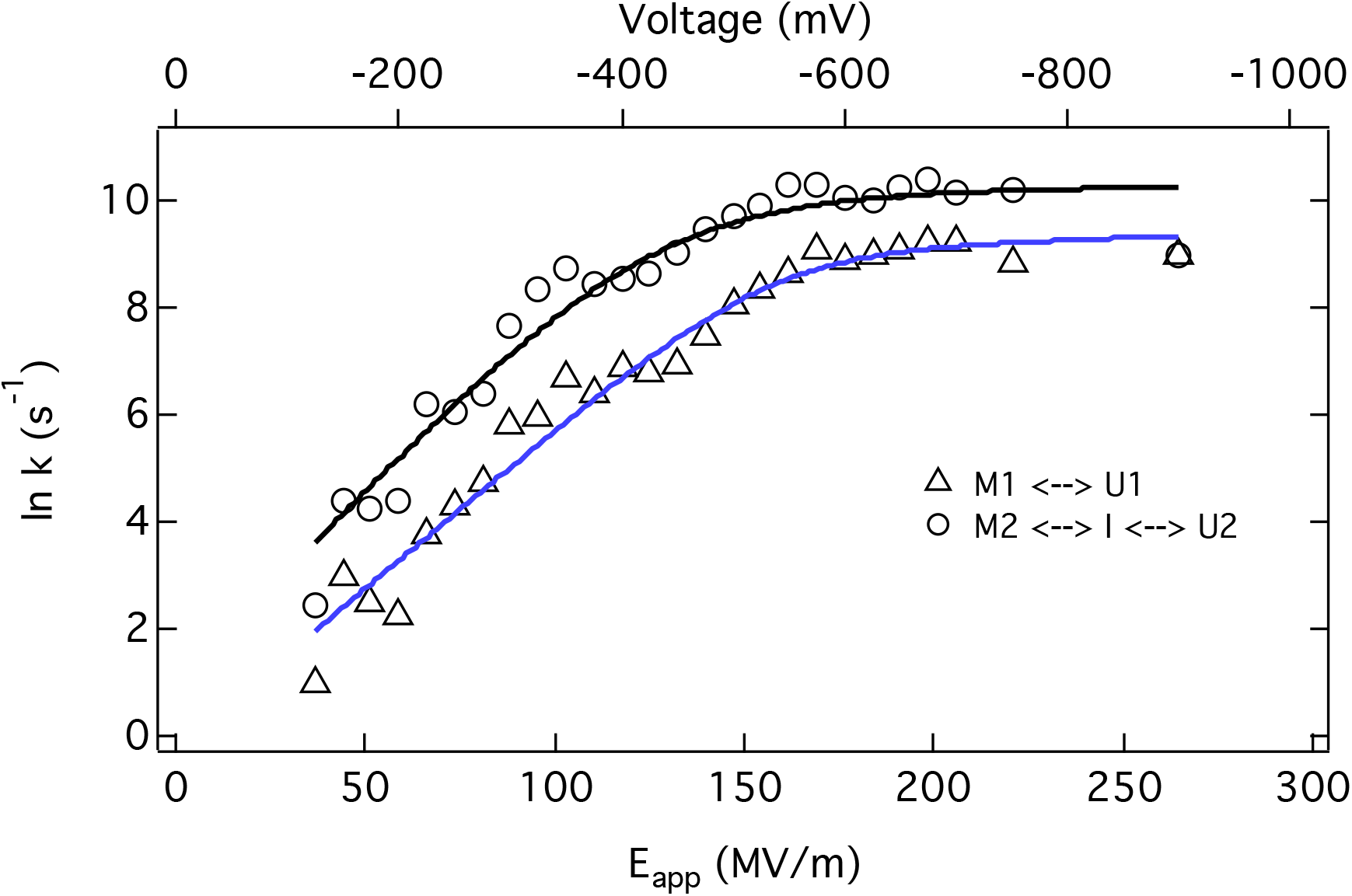
Fit of translocation rates (d_pore_ =2.5 nm, L=3.4nm) with Eq. S12 (blue curve) for black triangles and fit with Eq. S13 (black curve) for black circle. The parameters obtained from the fit are given Table S6.

**Table S6:** Extracted parameters from the fit of kinetic data (Fig. S18) from Eq. S12 (triangles) and Eq. S13 (circles).

| Parameters | | $M_1 \leftrightarrow U_1$ |
| --- | --- | --- |
| $k_{0U_1}$ | | $5839 \text{ s}^{-1}$ |
| $\beta_{U_1}$ | | 3.02 Debye |

| Parameters | | $M_2 \leftrightarrow I \leftrightarrow U_2$ |
| --- | --- | --- |
| $k_{0I}$ | | $72 \text{ s}^{-1}$ |
| $\beta_I$ | | 47.2 Debye |
| $k_{0U_2}$ | | $10712 \text{ s}^{-1}$ |
| $\beta_{U_2}$ | | 1.75 Debye |

## 18. Extraction of zero-field translocation barrier of *U* state for a 2 nm pore (L =4.1 nm)

To extract the translocation barrier of *U*-state we used time constant *τ*_*ii*2_ of life-time histograms of level *ii* (left inset Fig. 2B and Fig. 2E). This is because only the time constant of level *ii* decreases with voltage (Fig. 2F) which is indicative of successful translocation unlike *τ*_*ii*1_ which does not change with voltage in this region. The value of 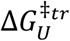 for the 2nm pore (Fig. S19) is about 2 *k*_*B*_*T*_*0*_ greater than the value 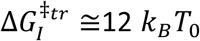 found for the 2.5 nm pore and at least 2 times greater than 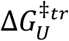 for 2.5 nm pore (see discussion following Eq. 8 of main text). This, along with the kinetic analysis of translocation rates, strongly suggests that the value of 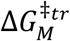 for 2nm is likely to be much greater than the unfolding energy barriers 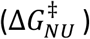 and that squeezing of the metastable *M*-state into the 2 nm pore is kinetically unlikely.

**Figure S19.**
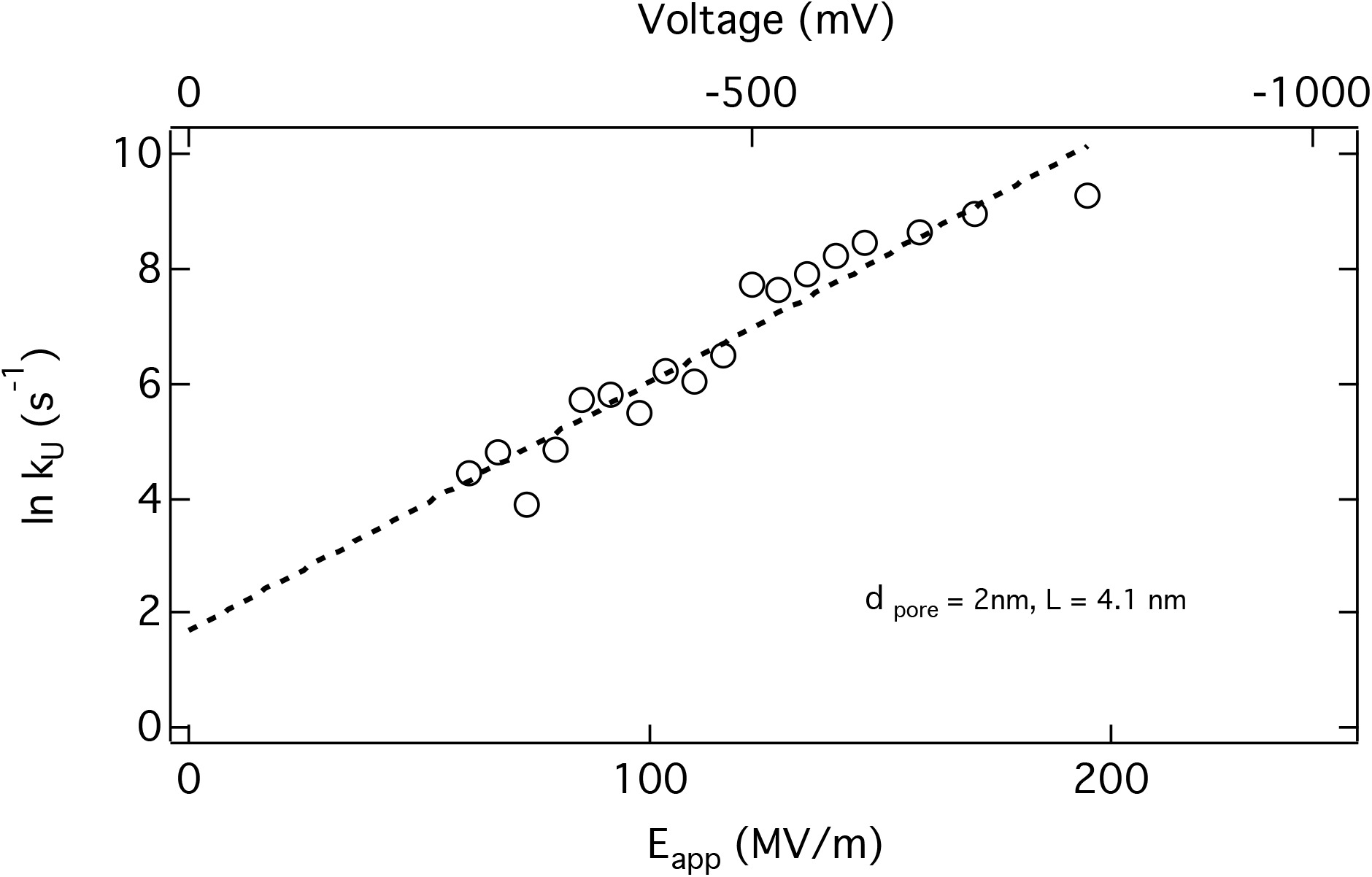
The rate of *U*-state translocation as a function of applied voltage and electric field. The dashed line is linear fit of the data 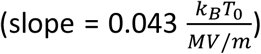. The extrapolation of rate at zero electric field yields ln *ko* = 1.68. Using *Eq. 8* of main text of the paper and L = 4.1 nm pore, the zero-field translocation barrier 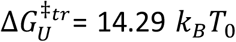. Here *k*_*U*_ = 1/*τ*_*ii*2_, the rate for the unfolded and successfully threaded conformation in the 2 nm pore.

## 19. Extraction of zero-field translocation barriers for a 2.5 nm pore (L =3.4 nm)

Here, we assigned two distinct rates in the regime 30-100 MV/m to the two metastable states, *M*_1_ and *M*_2_, which must cross their respective zero field translocation free energy barriers 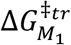 and 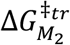, differing by a small amount due to the specifics of the trapping configurations of the squeezed M-state within the pore. Between 100-170 MV/m, *In* [*k*] increases with a smaller slope. This behavior is attributed to the additional flexibility associated with the dynamic unfolding of cytochrome c associated with interconversions between states *M, I*, and *U* (*M*_1_ ↔ *U*_1_ and M_2_ ↔ *I ↔ U*_2_). In the regime above 170 MV/m, *In* [*k*] has a still weaker dependence on the electric field. We attribute this to translocation that primarily involves only the unfolded states associated with U_1_ and U_2_. As an alternative to using the fitting parameters, *k*_0*I*_, noted in Table S6 to evaluate the values of 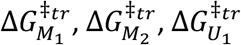, and 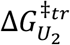, we can directly extrapolate the translocation rates to zero electric field (*k*_0_) and use Eq. 8 of the main text.

**Figure S20.**
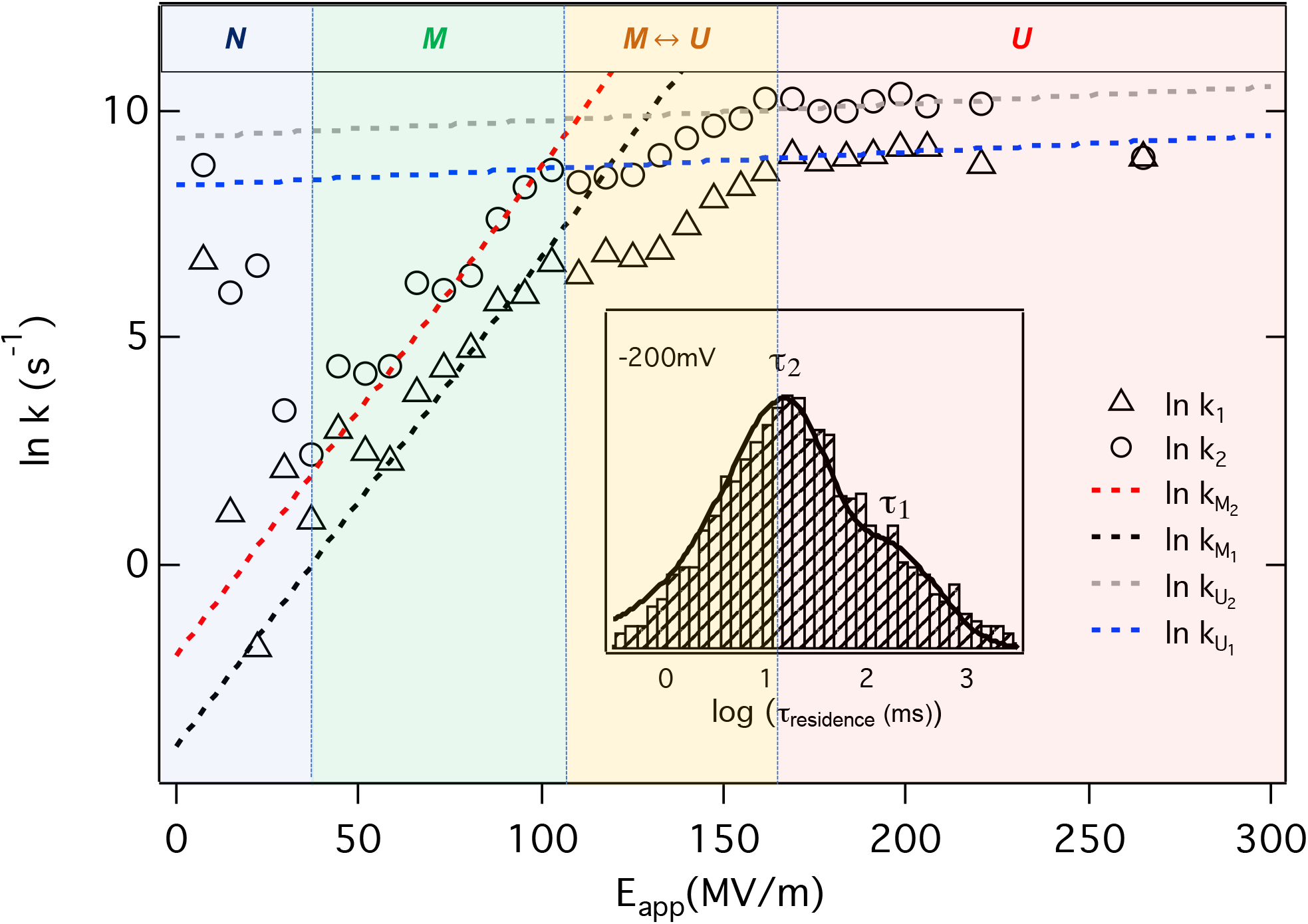
Extrapolation of translocation rates at zero electric-field (2.5 nm pore, L=3.4 nm). By extrapolating the rates to zero field with a linear fit of the data (dashed lines), we have evaluated the zero field Δ*G*^*‡*tr^ for each state (see Table S7). Slopes of the dashed lines (which represents electrophoretic force constants or *β*-values) are 0.11, and 0.004 for the *M*, and *U* regimes, respectively (in units of 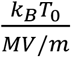). In the intermediate *M* ↔ *U* regime, the slope is 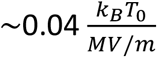.

**Table S7:** Extrapolated translocation rate constants (From Fig. S20) of conformationally excited states (*M*_1_, *M*_2_, *U*_1_, and *U*_2_) of cyt c at zero electric field and the corresponding values of electrophoretic force constant or *β*-value and 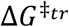 (using Eq. 8 of the main text).

| States | $\ln [k_0] \text{ (s}^{-1}\text{)}$ | $\beta \text{ (Debye)}$ | $\Delta G^{\ddagger tr} (k_B T_0)$ |
| --- | --- | --- | --- |
| $M_1$ | -4 | 135.9 | 20.35 |
| $M_2$ | -1.9 | 135.9 | 18.25 |
| $U_1$ | 8.35 | 4.9 | 8 |
| $U_2$ | 9.4 | 4.9 | 6.95 |

## 20. Open pore current vs voltage measurments for different pore diameters

**Figure S21.**
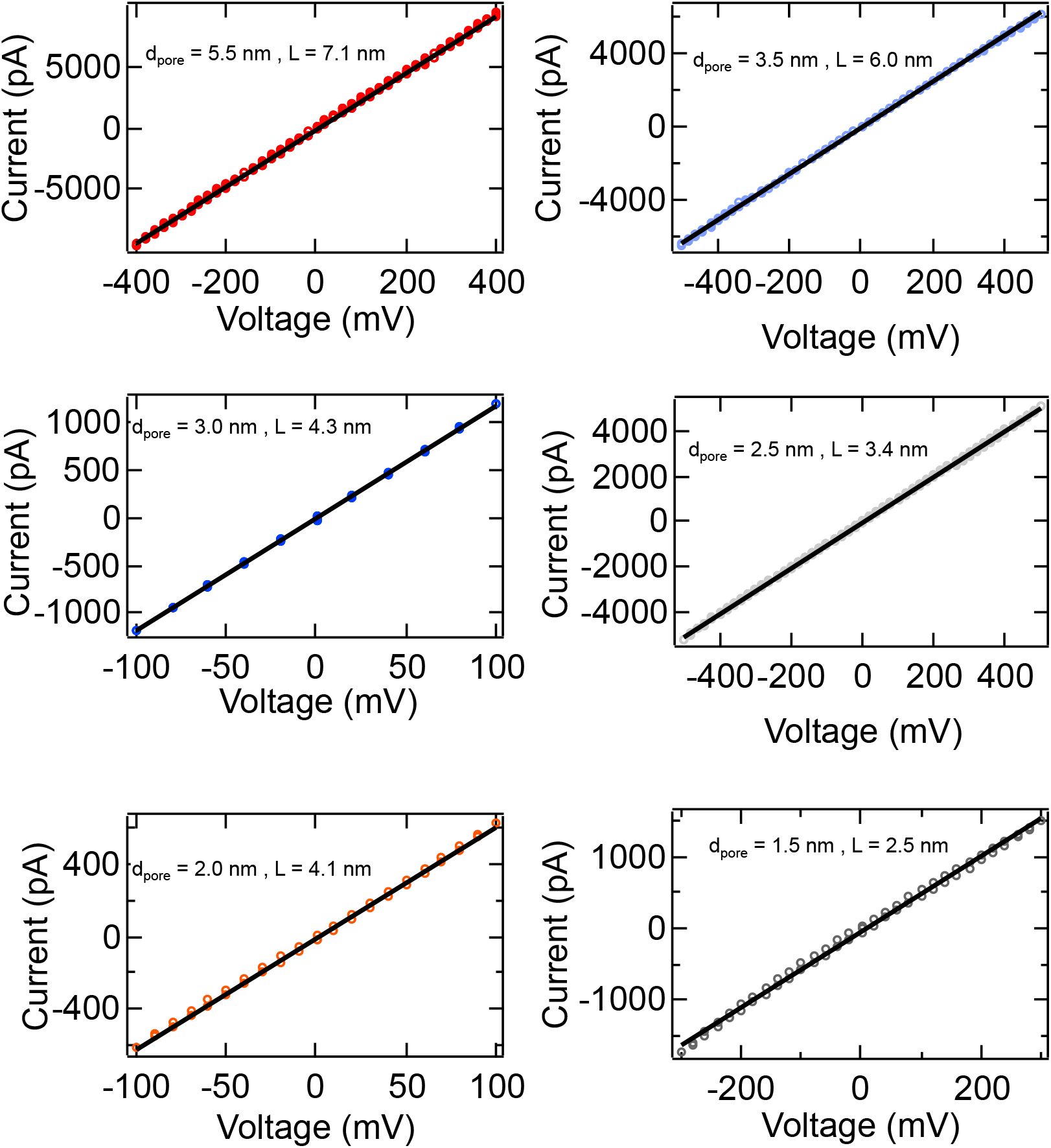
Open-pore (I_o_) current vs voltage measurments for different pore diameters employed in this study. The colored circles are the measured open pore currents at a given voltages and dark black lines are the linear fit of the data.

## Notes

**Competing Interest Statement**: No competing interests.

### Competing Interest Statement

The authors have declared no competing interest.

## References

1. B. Alberts, Molecular biology of the cell (Garland Science, New York, 2008).

2. W. Wickner, R. Schekman, Protein translocation across biological membranes. Science 310, 1452–1456 (2005).

3. S. Huang, K. S. Ratliff, A. Matouschek, Protein unfolding by the mitochondrial membrane potential. Nat. Struct. Biol. 9, 301–307 (2002).

4. R. J. Collier, Membrane translocation by anthrax toxin. Mol. Aspects. Med. 30, 413–422 (2009).

5. A. Martin, T. A. Baker, R. T. Sauer, Pore loops of the AAA+ ClpX machine grip substrates to drive translocation and unfolding. Nat. Struct. Mol. Biol. 15, 1147–1151 (2008).

6. S. Orrenius, B. Zhivotovsky, Cardiolipin oxidation sets cytochrome c free. Nat. Chem. Biol. 1, 188–189 (2005).

7. G. Oukhaled et al., Unfolding of proteins and long transient conformations detected by single nanopore recording. Phys. Rev. Lett. 98, 158101 (2007).

8. B. Cressiot et al., Protein Transport through a Narrow Solid-State Nanopore at High Voltage: Experiments and Theory. ACS Nano 6, 6236–6243 (2012).

9. A. Oukhaled et al., Dynamics of Completely Unfolded and Native Proteins through Solid-State Nanopores as a Function of Electric Driving Force. ACS Nano 5, 3628–3638 (2011).

10. M. Pastoriza-Gallego et al., Dynamics of Unfolded Protein Transport through an Aerolysin Pore. J. Am. Chem. Soc. 133, 2923–2931 (2011).

11. J. Nivala, D. B. Marks, M. Akeson, Unfoldase-mediated protein translocation through an alpha-hemolysin nanopore. Nat. Biotechnol. 31, 247–250 (2013).

12. D. Rodriguez-Larrea, H. Bayley, Multistep protein unfolding during nanopore translocation. Nat. Nanotechnol. 8, 288–295 (2013).

13. D. Rodriguez-Larrea, H. Bayley, Protein co-translocational unfolding depends on the direction of pulling. Nat. Commun. 5, 4841 (2014).

14. D. S. Talaga, J. Li, Single-molecule protein unfolding in solid state nanopores. J. Am. Chem. Soc. 131, 9287–9297 (2009).

15. D. J. Niedzwiecki, J. Grazul, L. Movileanu, Single-molecule observation of protein adsorption onto an inorganic surface. J. Am. Chem. Soc. 132, 10816–10822 (2010).

16. J. Houghtaling et al., Estimation of Shape, Volume, and Dipole Moment of Individual Proteins Freely Transiting a Synthetic Nanopore. ACS Nano 13, 5231–5242 (2019).

17. C. Plesa et al., Fast Translocation of Proteins through Solid State Nanopores. Nano Lett. 13, 658–663 (2013).

18. E. C. Yusko et al., Controlling protein translocation through nanopores with bio-inspired fluid walls. Nat. Nanotechnol. 6, 253–260 (2011).

19. K. J. Freedman, S. R. Haq, J. B. Edel, P. Jemth, M. J. Kim, Single molecule unfolding and stretching of protein domains inside a solid-state nanopore by electric field. Sci. Rep. 3, 1638 (2013).

20. J. Larkin, R. Y. Henley, M. Muthukumar, J. K. Rosenstein, M. Wanunu, High-bandwidth protein analysis using solid-state nanopores. Biophys. J. 106, 696–704 (2014).

21. P. Waduge et al., Nanopore-Based Measurements of Protein Size, Fluctuations, and Conformational Changes. ACS Nano 11, 5706–5716 (2017).

22. R. Hu et al., Differential Enzyme Flexibility Probed Using Solid-State Nanopores. ACS Nano 12, 4494–4502 (2018).

23. L. Movileanu, Squeezing a single polypeptide through a nanopore. Soft Matter 4, 925–931 (2008).

24. M. Muthukumar, POLYMER TRANSLOCATION (CRC PRESS, Boca Raton, FL), (2019).

25. I. Nir, D. Huttner, A. Meller, Direct Sensing and Discrimination among Ubiquitin and Ubiquitin Chains Using Solid-State Nanopores. Biophys. J. 108, 2340–2349 (2015).

26. O. K. Dudko, J. Mathe, A. Szabo, A. Meller, G. Hummer, Extracting kinetics from single-molecule force spectroscopy: nanopore unzipping of DNA hairpins. Biophys. J. 92, 4188–4195 (2007).

27. J. Mathe, H. Visram, V. Viasnoff, Y. Rabin, A. Meller, Nanopore unzipping of individual DNA hairpin molecules. Biophys. J. 87, 3205–3212 (2004).

28. J. Mathe, A. Aksimentiev, D. R. Nelson, K. Schulten, A. Meller, Orientation discrimination of single-stranded DNA inside the alpha-hemolysin membrane channel. Proc. Natl. Acad. Sci. U.S.A. 102, 12377–12382 (2005).

29. J. B. Heng et al., Stretching DNA using the electric field in a synthetic nanopore. Nano Lett. 5, 1883–1888 (2005).

30. S. Carson, J. Wilson, A. Aksimentiev, M. Wanunu, Smooth DNA transport through a narrowed pore geometry. Biophys. J. 107, 2381–2393 (2014).

31. Y. Bai, T. R. Sosnick, L. Mayne, S. W. Englander, Protein folding intermediates: native-state hydrogen exchange. Science 269, 192–197 (1995).

32. W. Hu, Z. Y. Kan, L. Mayne, S. W. Englander, Cytochrome c folds through foldon-dependent native-like intermediates in an ordered pathway. Proc. Natl. Acad. Sci. U.S.A. 113, 3809–3814 (2016).

33. H. Roder, G. A. Elove, S. W. Englander, Structural characterization of folding intermediates in cytochrome c by H-exchange labelling and proton NMR. Nature 335, 700–704 (1988).

34. P. M. De Biase et al., Molecular Basis for the Electric Field Modulation of Cytochrome c Structure and Function. J. Am. Chem. Soc. 131, 16248–16256 (2009).

35. M. Saito, S. J. Korsmeyer, P. H. Schlesinger, BAX-dependent transport of cytochrome c reconstituted in pure liposomes. Nat. Cell. Biol. 2, 553–555 (2000).

36. S. Shimizu, M. Narita, Y. Tsujimoto, Bcl-2 family proteins regulate the release of apoptogenic cytochrome c by the mitochondrial channel VDAC. Nature 399, 483–487 (1999).

37. Y. Chen et al., How can proteins enter the interior of a MOF? Investigation of cytochrome c translocation into a MOF consisting of mesoporous cages with microporous windows. J. Am. Chem. Soc. 134, 13188–13191 (2012).

38. G. Taler, A. Schejter, G. Navon, I. Vig, E. Margoliash, The nature of the thermal equilibrium affecting the iron coordination of ferric cytochrome c. Biochemistry 34, 14209–14212 (1995).

39. G. A. Elove, A. K. Bhuyan, H. Roder, Kinetic mechanism of cytochrome c folding: involvement of the heme and its ligands. Biochemistry 33, 6925–6935 (1994).

40. S. Berezhna, H. Wohlrab, P. M. Champion, Resonance Raman investigations of cytochrome c conformational change upon interaction with the membranes of intact and Ca2+-exposed mitochondria. Biochemistry 42, 6149–6158 (2003).

41. Y. Sun, V. Karunakaran, P. M. Champion, Investigations of the low-frequency spectral density of cytochrome c upon equilibrium unfolding. J. Phys. Chem. B 117, 9615–9625 (2013).

42. A. Benabbas, P. M. Champion, Adiabatic Ligand Binding in Heme Proteins: Ultrafast Kinetics of Methionine Rebinding in Ferrous Cytochrome c. J. Phys. Chem. B 122, 11431–11439 (2018).

43. L. C. Godoy et al., Disruption of the M80-Fe ligation stimulates the translocation of cytochrome c to the cytoplasm and nucleus in nonapoptotic cells. Proc. Natl. Acad. Sci. U.S.A 106, 2653–2658 (2009).

44. J. Hanske et al., Conformational properties of cardiolipin-bound cytochrome c. Proc. Natl. Acad. Sci. U.S.A 109, 125–130 (2012).

45. J. P. Kitt, D. A. Bryce, S. D. Minteer, J. M. Harris, Raman Spectroscopy Reveals Selective Interactions of Cytochrome c with Cardiolipin That Correlate with Membrane Permeability. J. Am. Chem. Soc. 139, 3851–3860 (2017).

46. L. J. McClelland, T. C. Mou, M. E. Jeakins-Cooley, S. R. Sprang, B. E. Bowler, Structure of a mitochondrial cytochrome c conformer competent for peroxidase activity. Proc. Natl. Acad. Sci. U.S.A 111, 6648–6653 (2014).

47. X. Kang, M. A. Alibakhshi, M. Wanunu, One-Pot Species Release and Nanopore Detection in a Voltage-Stable Lipid Bilayer Platform. Nano Lett. 19, 9145–9153 (2019).

48. C. Dekker, Solid-state nanopores. Nat. Nanotechnol. 2, 209–215 (2007).

49. M. Wanunu, Nanopores: A journey towards DNA sequencing. Phys. Life Rev. 9, 125–158 (2012).

50. M. Wanunu, W. Morrison, Y. Rabin, A. Y. Grosberg, A. Meller, Electrostatic focusing of unlabelled DNA into nanoscale pores using a salt gradient. Nat. Nanotechnol. 5, 160–165 (2010).

51. J. A. Ihalainen et al., α-Helix folding in the presence of structural constraints. Proc. Natl. Acad. Sci. U.S.A 105, 9588 (2008).

52. J. H. Werner, R. B. Dyer, R. M. Fesinmeyer, N. H. Andersen, Dynamics of the Primary Processes of Protein Folding:? Helix Nucleation. J. Phys. Chem. B 106, 487–494 (2002).

53. E. A. Gooding et al., The effects of individual amino acids on the fast folding dynamics of α-helical peptides. Chem. Comm. 10.1039/B511072F, 5985-5987 (2005).

54. K. Kuczera, G. S. Jas, R. Elber, Kinetics of Helix Unfolding: Molecular Dynamics Simulations with Milestoning. J. Phys. Chem. A 113, 7461–7473 (2009).

55. G. S. Jas, K. Kuczera, Helix–Coil Transition Courses Through Multiple Pathways and Intermediates: Fast Kinetic Measurements and Dimensionality Reduction. J. Phys. Chem. B 122, 10806–10816 (2018).

56. C. S. H. Jesus, P. F. Cruz, L. G. Arnaut, R. M. M. Brito, C. Serpa, One Peptide Reveals the Two Faces of α-Helix Unfolding–Folding Dynamics. J. Phys. Chem. B 122, 3790–3800 (2018).

57. W. Si, A. Aksimentiev, Nanopore Sensing of Protein Folding. ACS Nano 11, 7091–7100 (2017).

58. H. Maity, M. Maity, S. W. Englander, How cytochrome c folds, and why: submolecular foldon units and their stepwise sequential stabilization. J. Mol. Biol. 343, 223–233 (2004).

59. M. M. Krishna, Y. Lin, J. N. Rumbley, S. W. Englander, Cooperative omega loops in cytochrome c: role in folding and function. J. Mol. Biol. 331, 29–36 (2003).

60. P. P. Parui et al., Formation of Oligomeric Cytochrome c during Folding by Intermolecular Hydrophobic Interaction between N- and C-Terminal α-Helices. Biochemistry 52, 8732–8744 (2013).

61. S. Hirota et al., Cytochrome c polymerization by successive domain swapping at the C-terminal helix. Proc. Natl. Acad. Sci. U.S.A 107, 12854 (2010).

62. B. Dorvel et al., Analyzing the forces binding a restriction endonuclease to DNA using a synthetic nanopore. Nucleic Acids Res. 37, 4170–4179 (2009).

63. D. B. Wells, V. Abramkina, A. Aksimentiev, Exploring transmembrane transport through α-hemolysin with grid-steered molecular dynamics. J. Chem. Phys. 127, 125101 (2007).

64. E. Evans, Probing the Relation Between Force—Lifetime—and Chemistry in Single Molecular Bonds. Annu. Rev. Biophys. Biomol. Struct. 30, 105–128 (2001).

65. M. P. Eastwood, C. Hardin, Z. Luthey-Schulten, P. G. Wolynes, Evaluating protein structure-prediction schemes using energy landscape theory. IBM J. Res. Dev.45, 475–497 (2001).

66. J. Wilson, K. Sarthak, W. Si, L. Gao, A. Aksimentiev, Rapid and Accurate Determination of Nanopore Ionic Current Using a Steric Exclusion Model. ACS Sens. 4, 634–644 (2019).

67. M. Wanunu et al., Rapid electronic detection of probe-specific microRNAs using thin nanopore sensors. Nat. Nanotechnol. 5, 807–814 (2010).

68. M. J. Kim, M. Wanunu, D. C. Bell, A. Meller, Rapid Fabrication of Uniformly Sized Nanopores and Nanopore Arrays for Parallel DNA Analysis. Adv. Mater. 18, 3149–3153 (2006).

69. S. W. Kowalczyk, A. Y. Grosberg, Y. Rabin, C. Dekker, Modeling the conductance and DNA blockade of solid-state nanopores. Nanotechnology 22, 315101 (2011).

## References

1. M. Wanunu, W. Morrison, Y. Rabin, A. Y. Grosberg, A. Meller, Electrostatic focusing of unlabelled DNA into nanoscale pores using a salt gradient. Nat. Nanotechnol. 5, 160–165 (2010).

2. M. Wanunu et al., Rapid electronic detection of probe-specific microRNAs using thin nanopore sensors. Nat. Nanotechnol. 5, 807–814 (2010).

3. S. Carson, J. Wilson, A. Aksimentiev, M. Wanunu, Smooth DNA transport through a narrowed pore geometry. Biophys. J. 107, 2381–2393 (2014).

4. J. Larkin, R. Y. Henley, M. Muthukumar, J. K. Rosenstein, M. Wanunu, High-bandwidth protein analysis using solid-state nanopores. Biophys. J. 106, 696–704 (2014).

5. P. Waduge et al., Nanopore-Based Measurements of Protein Size, Fluctuations, and Conformational Changes. ACS Nano 11, 5706–5716 (2017).

6. H. Yamazaki et al., Label-Free Single-Molecule Thermoscopy Using a Laser-Heated Nanopore. Nano Lett. 17, 7067–7074 (2017).

7. S. W. Kowalczyk, A. Y. Grosberg, Y. Rabin, C. Dekker, Modeling the conductance and DNA blockade of solid-state nanopores. Nanotechnology 22, 315101 (2011).

8. W. H. Koppenol, J. D. Rush, J. D. Mills, E. Margoliash, The dipole moment of cytochrome c. Mol. Biol. and Evol. 8, 545–558 (1991).

9. W. Humphrey, A. Dalke, K. Schulten, VMD: Visual molecular dynamics. J. Mol. Graph.14, 33–38 (1996).

10. G. W. Bushnell, G. V. Louie, G. D. Brayer, High-resolution three-dimensional structure of horse heart cytochrome c. J. Mol. Biol. 214, 585–595 (1990).

11. W. L. Jorgensen, J. Chandrasekhar, J. D. Madura, R. W. Impey, M. L. Klein, Comparison of simple potential functions for simulating liquid water. J. Chem. Phys. 79, 926–935 (1983).

12. J. C. Phillips et al., Scalable molecular dynamics on CPU and GPU architectures with NAMD. J. Chem. Phys. 153, 044130 (2020).

13. A. D. MacKerell et al., All-atom empirical potential for molecular modeling and dynamics studies of proteins. J. Phys. Chem. B 102, 3586–3616 (1998).

14. J. Comer, V. Dimitrov, Q. Zhao, G. Timp, A. Aksimentiev, Microscopic Mechanics of Hairpin DNA Translocation through Synthetic Nanopores. Biophys. J. 96, 593–608 (2009).

15. U. Essmann et al., A smooth particle mesh Ewald method. J. Chem. Phys. 103, 8577–8593 (1995).

16. G. J. Martyna, D. J. Tobias, M. L. Klein, Constant pressure molecular dynamics algorithms. J. Chem. Phys. 101, 4177–4189 (1994).

17. H. J. C. Berendsen, J. P. M. Postma, W. F. van Gunsteren, A. DiNola, J. R. Haak, Molecular dynamics with coupling to an external bath. J. Chem. Phys. 81, 3684–3690 (1984).

18. D. B. Wells, V. Abramkina, A. Aksimentiev, Exploring transmembrane transport through α-hemolysin with grid-steered molecular dynamics. J. Chem. Phys. 127, 125101 (2007).

19. A. Aksimentiev, K. Schulten, Imaging α-Hemolysin with Molecular Dynamics: Ionic Conductance, Osmotic Permeability, and the Electrostatic Potential Map. Biophys. J. 88, 3745–3761 (2005).

20. J. Wilson, K. Sarthak, W. Si, L. Gao, A. Aksimentiev, Rapid and Accurate Determination of Nanopore Ionic Current Using a Steric Exclusion Model. ACS Sens. 4, 634–644 (2019).

21. M. P. Eastwood, C. Hardin, Z. Luthey-Schulten, P. G. Wolynes, Evaluating protein structure-prediction schemes using energy landscape theory. IBM J. Res. Dev. 45, 475–497 (2001).

22. C. G. Rodrigues, D. C. Machado, S. F. Chevtchenko, O. V. Krasilnikov, Mechanism of KCl enhancement in detection of nonionic polymers by nanopore sensors. Biophys. J. 95, 5186–5192 (2008).

23. W. Hu, Z. Y. Kan, L. Mayne, S. W. Englander, Cytochrome c folds through foldon-dependent native-like intermediates in an ordered pathway. Proc. Natl. Acad. Sci. U.S.A. 113, 3809–3814 (2016).

24. J. A. Ihalainen et al., α-Helix folding in the presence of structural constraints. Proc. Natl. Acad. Sci. U.S.A. 105, 9588 (2008).

25. J. H. Werner, R. B. Dyer, R. M. Fesinmeyer, N. H. Andersen, Dynamics of the Primary Processes of Protein Folding:? Helix Nucleation. J. Phys. Chem. B 106, 487–494 (2002).

26. E. A. Gooding et al., The effects of individual amino acids on the fast folding dynamics of α-helical peptides. Chem. Comm. 10.1039/B511072F, 5985-5987 (2005).

27. M. M. Krishna, Y. Lin, J. N. Rumbley, S. W. Englander, Cooperative omega loops in cytochrome c: role in folding and function. J. Mol. Biol. 331, 29–36 (2003).

28. H. Maity, M. Maity, S. W. Englander, How cytochrome c folds, and why: submolecular foldon units and their stepwise sequential stabilization. J. Mol. Biol. 343, 223–233 (2004).

29. S. van Dorp, U. F. Keyser, N. H. Dekker, C. Dekker, S. G. Lemay, Origin of the electrophoretic force on DNA in solid-state nanopores. Nat. Phys. 5, 347–351 (2009).

